# Novel mechanisms of MITF regulation and melanoma predisposition identified in a mouse suppressor screen

**DOI:** 10.1101/2023.08.04.551952

**Authors:** Hong Nhung Vu, Matti Már Valdimarsson, Sara Sigurbjörnsdóttir, Kristín Bergsteinsdóttir, Julien Debbache, Keren Bismuth, Deborah A. Swing, Jón H. Hallsson, Lionel Larue, Heinz Arnheiter, Neal G. Copeland, Nancy A. Jenkins, Petur O. Heidarsson, Eiríkur Steingrímsson

## Abstract

MITF, a basic-Helix-Loop-Helix Zipper (bHLHZip) transcription factor, plays vital roles in melanocyte development and functions as an oncogene. To explore MITF regulation and its role in melanoma, we conducted a genetic screen for suppressors of the Mitf-associated pigmentation phenotype. An intragenic Mitf mutation was identified, leading to termination of MITF at the K316 SUMOylation site and loss of the C-end intrinsically disordered region (IDR). The resulting protein is more nuclear but less stable than wild-type MITF and retains DNA-binding ability. Interestingly, as a dimer, it can translocate wild-type and mutant MITF partners into the nucleus, improving its own stability and ensuring an active nuclear MITF supply. Interactions between K316 SUMOylation and S409 phosphorylation sites across monomers largely explain the observed effects. Notably, the recurrent melanoma-associated E318K mutation in MITF, which affects K316 SUMOylation, also alters protein regulation in concert with S409, unraveling a novel regulatory mechanism with unexpected disease insights.

## Introduction

Transcription factors play a crucial role in gene regulation, and most of them have large unstructured domains termed intrinsically disordered regions (IDRs) in addition to their DNA-binding domains^1^. Due to the lack of tools and links to phenotypes, understanding the structure-function relationships of IDRs and their specific contributions to *in vivo* activity and disease has proven challenging. The basic Helix-Loop-Helix-leucine zipper (bHLHZip) transcription factor MITF is the master regulator of melanocyte development and pigmentation. It also plays a critical role in melanoma, a highly aggressive skin cancer originating from melanocytes^2, 3^. Importantly, MITF protein activity can be modulated either transiently through environmental signals or permanently by mutations leading to critical effects on the phenotype. In melanoma, MITF activity mediates phenotype plasticity such that high MITF activity promotes differentiation and proliferation, whereas low MITF activity results in a stem cell-like phenotype and enhances migration^2^. MITF binds to E-(CACGTG) and M- (TCATGTG) box motifs as a homodimer or as a heterodimer with its closest relatives, TFE3, TFEB, and TFEC^4, 5^. A unique 3-amino acid sequence in the zipper domain restricts dimerization of these proteins such that they do not dimerize with other bHLHZip proteins^6–8^. Outside the bHLHZip domain, MITF consists of N-terminal and C-terminal IDRs, located on either side of the bHLHZip DNA-binding and dimerization domain, largely of unknown function.

Importantly, multiple phosphorylation sites have been mapped in the IDRs of MITF^9^, and some (including S69, S73, and S173) have been suggested to lead to nuclear export or retention of MITF in the cytoplasm, some (S73 and S409) to affect transcription regulation and other sites have been proposed to affect protein stability (S73, S397, S401, S405, and S409) (Figure 1A)^9^. Interestingly, the S73 and S409 residues have been shown to be priming sites for GSK3β-mediated phosphorylation of downstream residues (S69 in the case of S73 and S397, S401 and S405 in the case of S409^10, 11^. MITF has also been shown to be SUMOylated at K182 and K316^12, 13^ and potentially ubiquitinylated at K201 and K265^14, 15^. However, the biological function of the different post-translational modifications (PTMs) is largely unknown.

**Figure 1:**
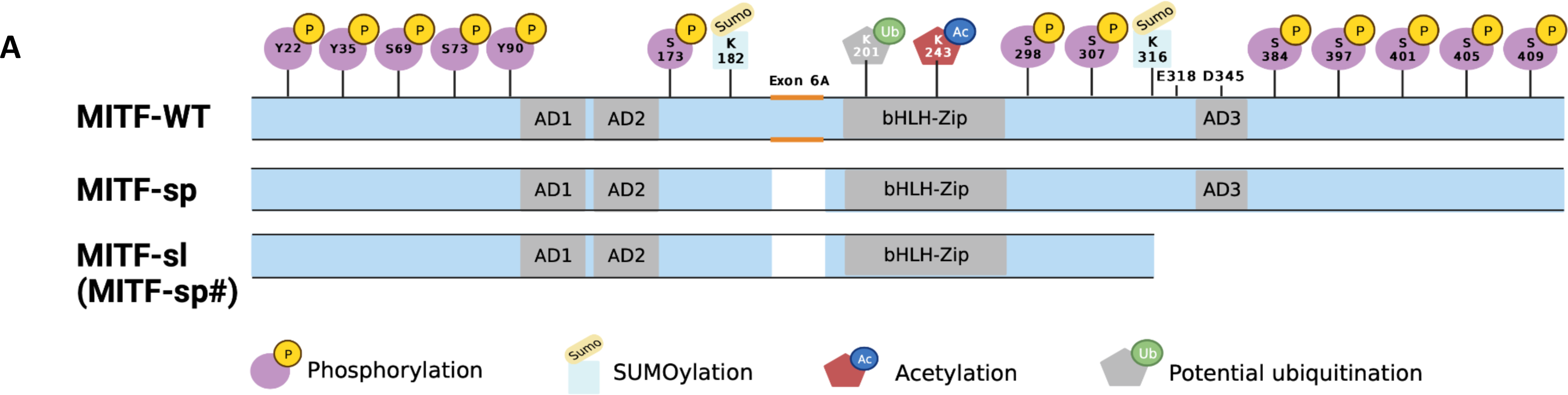
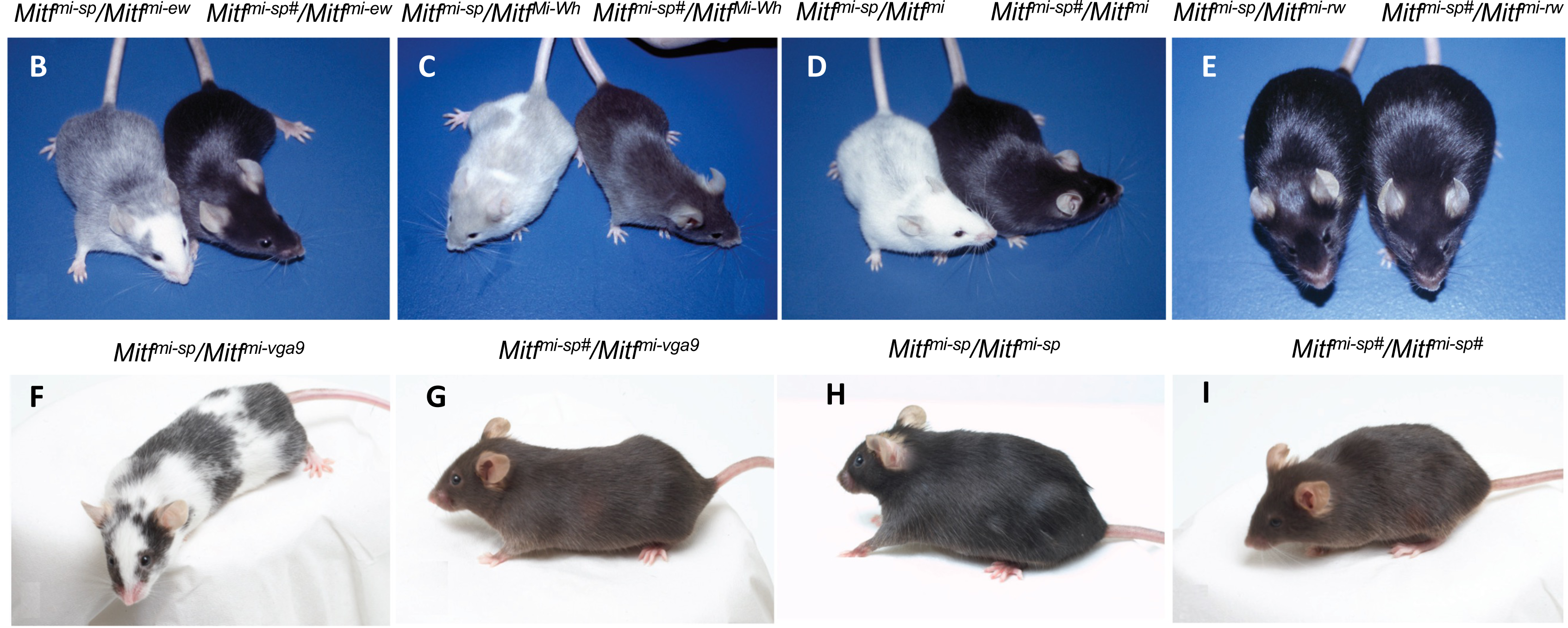
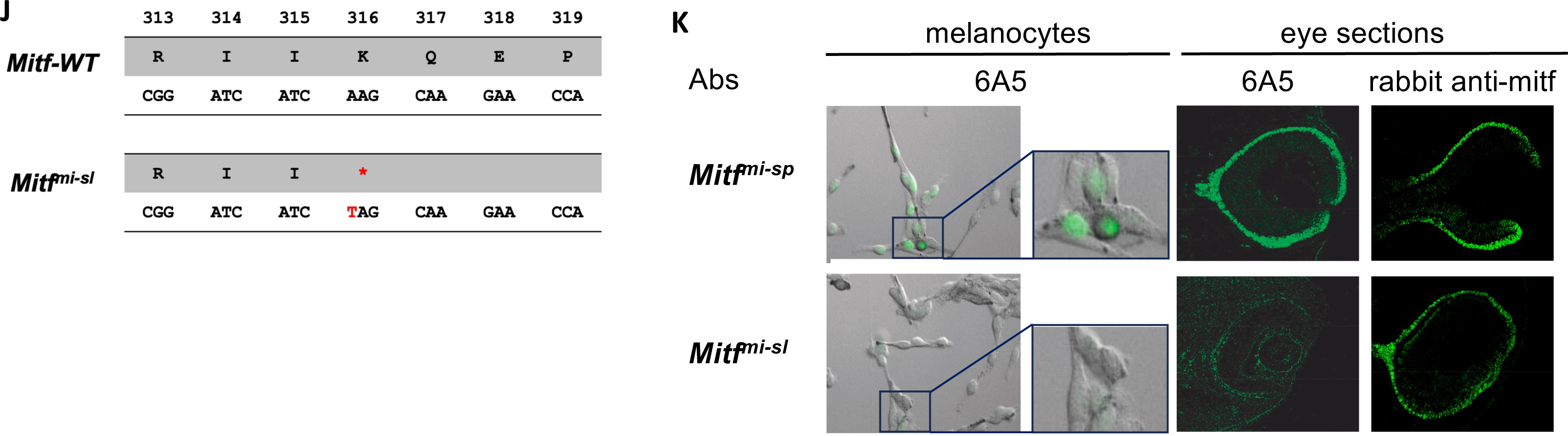
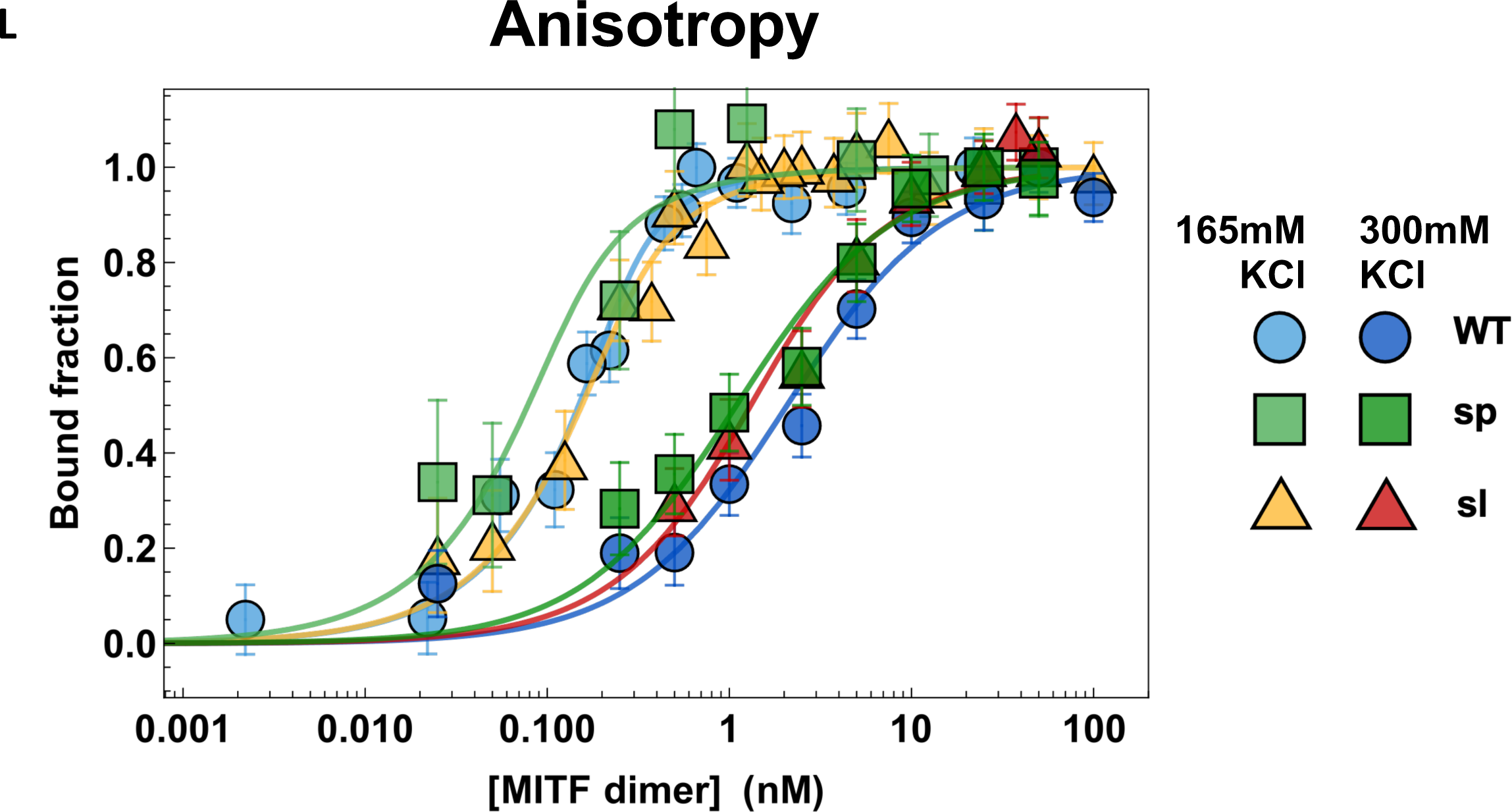
Coat color phenotypes and molecular alteration associated with the induced *Mitf^mi-sp#^* suppressor mutation (*Mitf^mi-sl^*) (A) Graphical depiction of the MITF-WT, MITF-sp, and MITF-sl (MITF-sp#) proteins indicating the domains affected. Also shown are the post-translational modifications that have been reported in MITF. (B) NAW-*Mitf^mi-ew^*/B6-*Mitf^mi-sp^* and NAW-*Mitf^mi-ew^*/B6-*Mitf^mi-sp#^* compound heterozygotes. (C) B6-*Mitf^Mi-Wh^*/B6-*Mitf^mi-sp^* and B6-*Mitf^Mi-Wh^*/B6-*Mitf^mi-sp#^* compound heterozygotes. (D) B6-*Mitf^mi-sp^*/B6-*Mitf^mi^ and B6-Mitf^mi-sp#^*/B6-*Mitf^mi^* compound heterozygotes. Notice the dramatic suppression of the phenotype from near-white to black coat color. (E) B6-*Mitf^mi-sp^*/B6-*Mitf^mi-rw^* and B6-*Mitf^mi-sp#^*/B6-*Mitf^mi-rw^* animals. (F) *B6-Mitf^mi-sp^*/B6-*Mitf^mi-vga9^*. (G) *B6-Mitf^mi-sp#^*/B6-*Mitf^mi-vga9^*. (H) B6-*Mitf^mi-sp^*/*Mitf^mi-sp^*. (I) B6-*Mitf^mi-sp^ ^#^*/*Mitf^mi-sp#^*. (J) Graphical depiction of the *Mitf^mi-sl^* mutation. (K) Antibody staining of melanocytes and eye sections from *Mitf^mi-sp^* and *Mitf^mi-sl^*tissues. The antibodies are 6A5, which recognizes the C-end of MITF, and a polyclonal rabbit anti-MITF antibody. (L) DNA binding curves of recombinantly expressed human MITF-WT, MITF-sp, and MITF-sl proteins to M-box probe measured by Fluorescence anisotropy at 165 mM KCl and 300 mM KCl. MITF-WT protein in light and dark blue circles, MITF-sp light and dark green boxes, and MITF-sl in yellow and red triangles. Error bars represent two standard deviations of fit error at each point.

Individuals carrying the E318K germline mutation in MITF are predisposed to melanoma^16, 17^. Carriers of a single allele have increased nevus count and are predisposed to melanoma; one of the initial reports also showed an increased risk of renal cell carcinoma ^16^, and a link to uterine carcinosarcoma has also been described^18^. The E318K mutation abolishes SUMOylation of the MITF protein at K316^16, 17, 19^, and ChIP-seq studies have shown that the MITF-E318K protein has increased occupancy at known MITF-target genes compared to the wild type protein but also binds to an increased number of genes.

However, transcriptomic studies did not reveal major changes in gene expression^16, 17^. Mice carrying the E318K mutation exhibited slightly reduced pigmentation in both homo- and heterozygous conditions, whereas *Mitf^E318K^*/+; *Braf^V600E^*/+ mice had an increased number of nevi. While the E318K mutation did not affect the latency or morphology of *Braf^V600E^* tumors, it accelerated tumor initiation in *Braf^V600E^*/+; Pten-/-mice^19^. Currently, it is not understood how the E318K mutation affects protein function or how it predisposes to melanoma.

Due to its obvious effects on pigmentation, MITF provides an excellent sensitized system for searching for suppressor mutations. In mice, over 40 different mutant alleles have been found in *Mitf* that can be arranged in an allelic series according to the severity of their phenotypic effects as evidenced by coat color changes^20^. At one end of the spectrum is the original and most severe allele *Mitf^mi^* (Table S1; deletion of one of four arginines in the DNA-binding domain), which leads to a white coat, severe microphthalmia, and osteopetrosis and results in death at 3-4 weeks of age. At the other end of the spectrum is the mildest *Mitf* mutation, *Mitf^mi-spotted^* (*Mitf^mi-sp^)*, which has no visible phenotype even when homozygous. The *Mitf^mi-sp^*mutation lacks the alternative 18bp exon 6A that encodes six amino acids upstream of the DNA-binding domain (Figure 1A). Interestingly, the *Mitf^mi-sp^* allele induces a white spotting phenotype when combined with other mutations at the locus^21, 22^. For example, when the *Mitf^mi-sp^* allele is mated to the original *Mitf^mi^* mutation, the offspring exhibit a white coat with occasional grey areas and no microphthalmia. The intermediate coat pigmentation alterations obtained in compound heterozygotes with *Mitf^mi-sp^* made this allele ideal for an N-ethyl-N-nitrosourea (ENU) mutagenesis screen for dominant suppressors or enhancers of the *Mitf* phenotype. Using this approach, we expected to find mutations in novel genes participating in the molecular pathways through which *Mitf* regulates pigment cell development and melanogenesis. We isolated a mutation that suppressed the *Mitf* phenotype, but intriguingly, it is a derivative of the *Mitf^mi-sp^*allele that lacks 104 residues of the carboxyl end (C-end). The induced *Mitf* suppressor mutation highlights the critical role of the IDR at the C-end of MITF in determining its stability, subcellular location, and transcriptional activity. Importantly, it also sheds light on the molecular effects of the E318K mutation in melanoma.

## Results

### Generation and analysis of an *Mitf* suppressor mutation

To screen for dominant mutations that suppress the *Mitf* phenotype, we crossed NAW-*Mitf^mi-ew^*/*Mitf^mi-ew^*females with C57BL/6J-*Mitf^mi-sp^/Mitf^mi-sp^* males that had been treated previously with the mutagen N-ethyl-N-nitrosourea (ENU). We screened for coat pigmentation changes in the F1 offspring (Figure S1A). While *Mitf^mi-sp^* homozygotes have no visible coat color phenotype, animals homozygous for the *Mitf^mi-ew^*mutation are white, severely microphthalmic, and exhibit mild hyperostosis^23^ (Table S1). Compound heterozygotes for these two mutations have a “salt-and-pepper” body color with a white head, belly, and feet (Figure 1B, left). ENU-treated F1 *Mitf^mi-sp^*/*Mitf^mi-ew^* heterozygotes were screened for coat pigmentation changes (Figure S1A). Of 63 NAW-*Mitf^mi-ew^*/*Mitf^mi-ew^* females, less than 50% produced progeny, resulting in a total of 470 offspring. In one of the matings, a deviant offspring female, marked by a ‘#’, showed a considerably darker coat (near-black coat with pale ears, tails, and toes) compared to its littermates, suggesting the presence of a mutagenized chromosome (Figure 1B, right).

When this *Mitf^mi-ew^/Mitf^mi-sp#^*female was bred to a C57BL/6J*-Mitf^mi-ew^*/*Mitf^mi-ew^*male, two classes of offspring resulted: white microphthalmic mice of the genotype *Mitf^mi-ew^*/*Mitf^mi-ew^* and mice of the genotype *Mitf^mi-ew^*/*Mitf^mi-sp#^* with the darkly pigmented phenotype of their mother (Figure S1B, right). This confirmed that the ‘#’ mutation altering the *Mitf^mi-ew^*/*Mitf^mi-sp^* phenotype is dominant, at least for the combination of these two alleles. Also, because the above crosses did not yield mice with the phenotype expected for *Mitf^mi-ew^/Mitf^mi-sp^* mice, the novel suppressor mutation is likely closely genetically linked with *Mitf^mi-sp^* or lies within the gene itself rather than on a different chromosome. Crossing the near-black *Mitf^mi-ew^*/*Mitf^mi-sp#^* mice to C57BL/6J animals only resulted in black offspring.

When the near-black *Mitf^mi-ew^/Mitf^mi-sp#^* mice were mated to white microphthalmic *Mitf^Mi-Wh^*/*Mitf^Mi-Wh^* homozygotes (Table S1), there were again two classes of offspring: the expected white mice with average eye size (*Mitf^mi-ew^*/*Mitf^Mi-Wh^* heterozygotes) and “steel”-colored mice with pale ears, tail, toes, and a belly spot (*Mitf^Mi-Wh^*/*Mitf^mi-sp#^* animals, Figure 1C, right). The coat color of the latter animals was much darker than that of the corresponding *Mitf^Mi-Wh^*/*Mitf^mi-sp^* animals (Figure 1C, left). The color was even darker than *Mitf^Mi-Wh^/Mitf-WT* mice, suggesting that the new mutant represents a gain-of-function compared to the wild type. Similar effects were also seen when animals carrying the new *Mitf^mi-sp#^* mutation were crossed to the dominant-negative *Mitf^mi^* (Figure 1D), *Mitf^Mi-or,^* and *Mitf^Mi-b^* mutations (Figures S1C-E) or the null mutation *Mitf^mi-vga^*^9^ (Figures 1F and 1G; Table S1). These observations showed that the suppressing effects of the new mutation were not restricted to the *Mitf^mi-ew^* allele and did not depend on the genetic background of the alleles tested (compare Figure 1B to S1B and S1C to S1D). However, the new mutation did not affect the coat color of the recessive *Mitf^mi-rw^* allele when compared to *Mitf^mi-rw^/Mitf^mi-sp^*animals (Figure 1E), reflecting the fact that the latter animals are already black and no further improvement in coat color is possible. Similarly, no effects were observed on eye size or bone development in any of the combinations since both phenotypes are normal in *Mitf^mi-sp^* homozygotes or their compound heterozygotes.

Intercrosses of *Mitf-WT/Mitf^mi-sp#^* heterozygotes produced two classes of offspring: normal non-agouti (black) mice and mice with a diluted “brownish” coat color in a 3 to 1 ratio (compare Figure 1H to 1I). Genotyping showed that the “brownish” animals were homozygous for *Mitf^mi-sp#^*. Thus, intriguingly, the new mutation suppresses the spotting phenotype of most combinations of *Mitf* alleles but causes the *Mitf^mi-sp#^*to express a partial loss-of-function, altering the coat color from black to brown.

### Molecular analysis of the *Mitf^mi-sp^*^#^ mutation

From the various crosses described above, it was clear that the new mutation is either tightly linked to *Mitf* on chromosome 6 or is an intragenic mutation. We, therefore, performed RT-PCR and sequencing studies of *Mitf* in total RNA isolated from homozygous *Mitf^mi-sp#^*heart and kidney. Consistent with the induced origin of the mutation, sequencing of the cDNA revealed the previously characterized *Mitf^mi-sp^*mutation (the lack of the 18 bp alternative exon)^24^. In addition, an A to T transversion was detected at nucleotide 1075 of the cDNA of the MITF-M isoform^25^, replacing the codon for K316 with a stop-codon resulting in premature truncation of the protein in the last exon, exon 9 (Figure 1J). The mutation was confirmed by genomic sequencing. The # mutation is, therefore, an intragenic re-mutation of the *Mitf^mi-sp^* allele, now termed *Mitf^mi-spotless^* (*Mitf^mi-sl^*), that leads to a protein, MITF^mi-sl^, that lacks 104 residues of the C-end, including the K316 SUMOylation site^12, 13^, a caspase cleavage site (D345)^26^, phosphorylation sites implicated in the mTOR, GSK3β, and MAP kinase signal transduction pathways (S384, S397, S401, S405, and S409)^9^ and the proposed transcription activation domain 3 (AD3)^27^ (Figure 1A).

To confirm that the C-end of MITF is missing from the *Mitf^mi-sl^* mutant, we stained primary melanocyte cultures generated from homozygous *Mitf^mi-sl^* and *Mitf^mi-sp^* embryos and eye sections from *Mitf^mi-sl^* and *Mitf^mi-sp^* mutants with the monoclonal antibody 6A5, which reacts with the C-end of MITF^28^ and should not stain cells or tissues from *Mitf^mi-sl^* animals. As shown in Figure 1K, the antibody did not give a signal in *Mitf^mi-sl^* melanocytes or eye sections, whereas *Mitf^mi-sp^* melanocytes and eye sections exhibited clear nuclear staining. In contrast, eye sections from both genotypes stained positive with a polyclonal rabbit anti-MITF antibody. This shows that the carboxyl-end (C-end) of MITF is absent from melanocytes and eyes of *Mitf^mi-sl^* homozygotes.

### *Mitf^mi-sl^/Mitf^mi-sl^* homozygotes show normal melanoblast development but delayed onset of pigmentation

To determine if the suppressor mutation affects cell proliferation, we performed a BrdU incorporation assay in the non-melanocytic HEK293 cells after overexpressing the MITF^mi-sp^ or MITF^mi-sl^ proteins (in the discussion below, we simplify the nomenclature of the mutants to MITF-sp, MITF-sl and so on). All the MITF mutants in this study were generated with the mouse MITF-M isoform. No significant difference was observed (69 ± 2% BrdU positivity in MITF-sp expressing cells versus 71 ± 2% BrdU positivity in MITF-sl expressing cells) (Figure S2A). IncuCyte live cell imaging also showed no significant changes in the proliferation of the human A375P melanoma cells (which express little endogenous MITF) upon overexpressing MITF-WT, MITF-sp or MITF-sl proteins over time (Figure S2B).

To count melanoblast numbers during development in vivo, we generated *Mitf^mi-sp^* and *Mitf^mi-sl^* homozygous embryos carrying a melanoblast marker transgene, Dct-LacZ^29^. In wild-type C57BL/6 embryos, an increased number of Dct-LacZ-labeled cells was observed with developmental time, as expected. Similar increases were seen in *Mitf^mi-sp^* and *Mitf^mi-sl^* homozygotes, although the total number of X-gal labeled cells seemed, on average, reduced compared to wild type (Figures S2C and S2D). The analysis of X-gal staining in the respective embryos suggests that there are generally fewer melanocytes in the two mutations, although there may be some region-specific differences.

In *Mitf^mi-sp^ ex vivo* neural crest cell cultures, pigmented melanocytes appeared on day 10 after explantation and increased steadily through day 17 (Figure S3A), whereas in *ex vivo* cultures from *Mitf^mi-ls^* embryos, they first emerged on day 14 and had a lower average count by day 17 than was observed in *Mitf^mi-sp^* embryos (Figure S3A). Nevertheless, in both *Mitf^mi-sp^* and *Mitf^mi-sl^*cultures, cells positive for the MITF protein (melanoblasts) were detected as early as day 1 of culture (Figure S3A, inset, for day 3 of culture), suggesting that the delay in pigmentation is not due to a delay in melanoblast appearance. Consistent with these observations, *Mitf^mi-sl^* homozygous pups develop pigmentation later than *Mitf^mi-sp^* pups (Figure S3B, the difference is particularly striking on postnatal days 2 and 3). However, this difference was only seen when the mice were homozygous for these alleles and not when the alleles were combined, for instance, with *Mitf^mi-vga9^*(Figure S3C, compare mice labeled #1 and #3) or with *Mitf-WT* (Figure S3C, compare mice labeled #2 and #4). Hence, the delayed onset of pigmentation in culture and the “brownish” color seen in adult *Mitf^mi-sl^* homozygotes suggest that the mutation leads to a partially functional protein. However, it is not caused by prolonged cell proliferation affecting differentiation initiation.

### The MITF-sl protein forms stable dimers and binds DNA

We determined the dimerization and DNA-binding ability of the MITF-sl protein. For this, we co-expressed Flag-tagged versions of the non-DNA binding mutant proteins MITF-Wh, MITF-mi, and MITF-ew (Table S1) together with the MITF-WT-GFP, MITF-sp-GFP, or MITF*-*sl*-*GFP proteins in A375P melanoma cells followed by co-immunoprecipitation (co-IP) using FLAG-antibodies. The results showed that the non-DNA binding MITF mutant proteins successfully immunoprecipitated all three GFP-labeled proteins (Figure S4A). We further confirmed the interactions between MITF-mi-Flag and GFP-tagged MITF-WT, -sp, and -sl proteins by Blue native PAGE ^30^. The results suggest that the MITF-mi protein as well as the MITF-WT, -sp, and -sl proteins can form both hetero- and homodimers (Figures S4B and S4C). We observed that in cells co-expressing MITF-mi and MITF-sl, heterodimers predominate over homodimers. Additionally, MITF-sl was found mostly as a dimer in cells expressing only MITF-sl (Figure S4B and S4C). Our findings suggest that MITF-sl may have a stronger dimerization affinity than MITF-WT.

We measured the DNA binding affinity of recombinant human MITF-WT, -sp, and -sl proteins to a fluorescently labeled M-box probe by measuring changes in anisotropy on individual molecular complexes with single-molecule spectroscopy. Quantification gave dissociation constants for all constructs at 300 mM KCl that were within one standard deviation from each other (Figures 1L, Table S2), while the affinities at 165 mM KCl (*K*_D_<250 pM) were too high to compare accurately. We also used single-molecule Förster resonance energy transfer (smFRET) and fluorescence correlation spectroscopy (FCS) as two additional and independent measures of MITF interactions with DNA (Figures S5A-D). All three methods yielded similar dissociation constants for all constructs at 300 mM KCl (Figures 1L, S5A-D and Table S2), and similar to that reported for the DNA binding domain alone^31^. From the FCS data, we observed a smaller change in diffusion time upon DNA binding of MITF-sl than -sp and -WT, consistent with its smaller size (Figure S5A, Table S2). The electrophoretic mobility shift assays (EMSA) also showed similar steady-state affinity binding to M-box DNA of mouse MITF-WT, -sp, and -sl (Figure S5E). Co-translating the non-DNA binding MITF-mi with MITF-WT, MITF-sp, and MITF-sl, thus allowing heterodimerization before EMSA showed that increasing amount of the MITF-mi protein interfered with the DNA binding of all three proteins. However, MITF-mi was more effective at interfering with MITF-WT than with either of the mutant proteins lacking exon 6A (Figure S5E), which is consistent with previous observations^6^. A slightly different picture emerged when the proteins were translated separately and subsequently mixed together before the EMSA. Again, the MITF-mi protein was more effective at interfering with the DNA binding of the MITF-WT protein than with that of the MITF-sp and MITF-sl proteins. However, it was even less effective at interfering with the DNA-binding of the MITF-sl protein (Figure S5F) than MITF-sp. This suggests that the MITF-sl homodimers are more stable than either the MITF-WT or MITF-sp homodimers and, thus, less prone to interference by a dominant negative protein such as the MITF-mi protein.

### The MITF-sl protein differentially affects gene expression

To determine if *Mitf* RNA expression was affected by the *Mitf^mi-sl^*mutation, we quantitated *Mitf* RNA expression in the heart, an organ that is not overtly involved in any of the extant *Mitf* mutant phenotypes. Quantitative RT-PCR showed that expression of *Mitf* RNA is reduced to 66% in *Mitf^mi-sp^* hearts compared to wild-type controls but remains above wild-type levels (116%) in *Mitf^mi-sl^*hearts (Figure S6). Similar observations were made when RNA obtained from the skin was quantitated (Figure S6).

To investigate the effects of the *Mitf^mi-sl^* mutation on gene expression, we induced the expression of mouse MITF proteins at an equal level in A375P cells and harvested RNA at regular intervals for qPCR analysis. The fold change of MITF target genes was first normalized to EV and then to the proportion of MITF proteins retained in the nucleus. Consistent with previous work^32, 33^, our results showed that the expression of the endogenous human *MITF* mRNA was considerably reduced over the 36 hours sampling period upon overexpressing mouse MITF-WT and MITF-sl proteins (Figure S7A). The expression of the *CDH2* and *NRP1* genes, both of which have been shown to be repressed by MITF (Dilshat et al., 2021), was also significantly reduced upon overexpression of MITF-WT and MITF-sl (Figures S7B and S7C). While MITF-WT activated the expression of *PMEL* and *TRIM63*, MITF-sl exhibited about half the activating ability of WT (Figures S7D and S7E). Interestingly, the MITF-sl protein was severely impaired in activating the expression of the pigmentation genes *TYRP1, MLANA, TYR, and DCT* (Figures S7F-I). Our results suggest that the 316-419 domain is critical for selective transcriptional activation, whereas the repressive activity of MITF remains intact, at least for the genes tested.

### The *Mitf^mi-sl^* mutation affects protein stability and localization

We also investigated the effect of the *Mitf^mi-sl^* mutation on protein stability by expressing Flag-tagged (at C-end) MITF-WT, MITF*-*sp, and MITF-sl proteins in a doxycycline (dox)-inducible vector transfected into A375P melanoma cells. Expression of the proteins was equalized by treating the cells with varying concentrations of dox for 24 hours. The cells were then treated with the translation inhibitor cycloheximide (CHX) for different periods and harvested to visualize MITF protein by Western blotting. The MITF protein is observed as two bands where the upper band is phosphorylated at S73 (hereafter referred to as pS73-MITF), and the lower band is not phosphorylated at S73 (hereafter referred to as S73-MITF)^10, 34^. The bands on the Western blot were quantitated, and the changes in protein concentration were plotted over time. This data was used to calculate protein half-life, defined as the time required to reduce the initial protein abundance to 50%. The MITF-WT and MITF-sp proteins had comparable half-lives of 3.2 hours for pS73-MITF and 1.2 hours for S73-MITF (Figures 2A-C). Critically, the MITF-sl protein was considerably less stable, with half-lives of 1.2 and 0.4 hours for the pS73 and S73 forms, respectively (Figures 2A-C). To confirm that exon 6A does not contribute substantially to MITF stability, we measured the stability of the pS73 and S73 versions of MITF-Wh with and without exon 6A and found that they were not significantly different (Figures S8A and S8B). When overexpressed in the 501Mel and SKmel28 melanoma cell lines (which express high levels of MITF), the MITF-sl protein was also less stable than the MITF-WT and MITF-sp proteins, regardless of the S73 phosphorylation status (Figures S8C-F). We also tested the stability of proteins carrying the Flag tag at the N-end or GFP tag at the C-end. In all cases, MITF-sl protein was less stable than MITF-WT and MITF-sp, regardless of the fusion tags and their location (Figures S8G-I). Our finding, therefore, suggests that the absence of the 316-419 domain significantly reduces the stability of MITF. Furthermore, in all cases, the S73 MITF (lower band) protein was degraded faster than the pS73 form (upper band). Alternatively, the S73 protein may be phosphorylated and thus become pS73 during the experiment.

**Figure 2:**
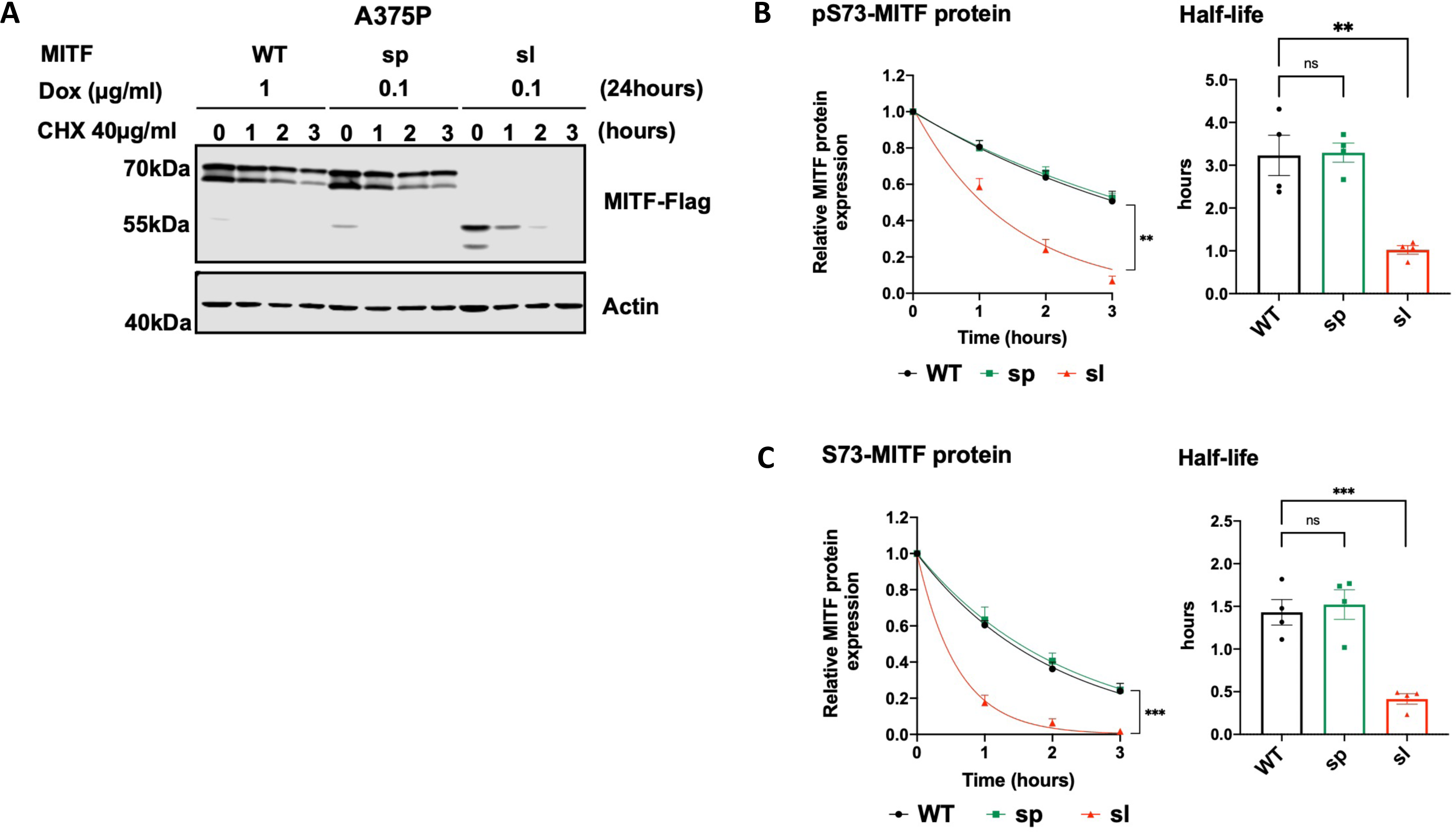
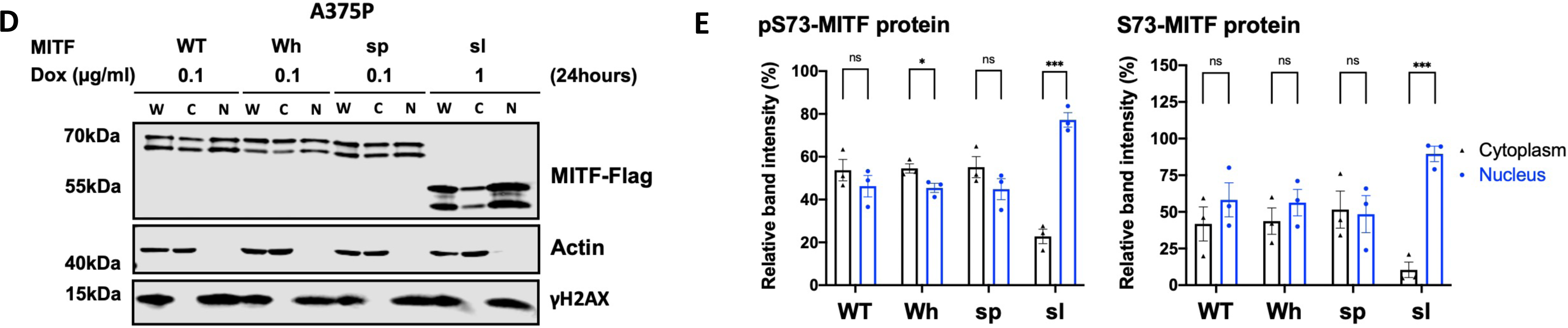
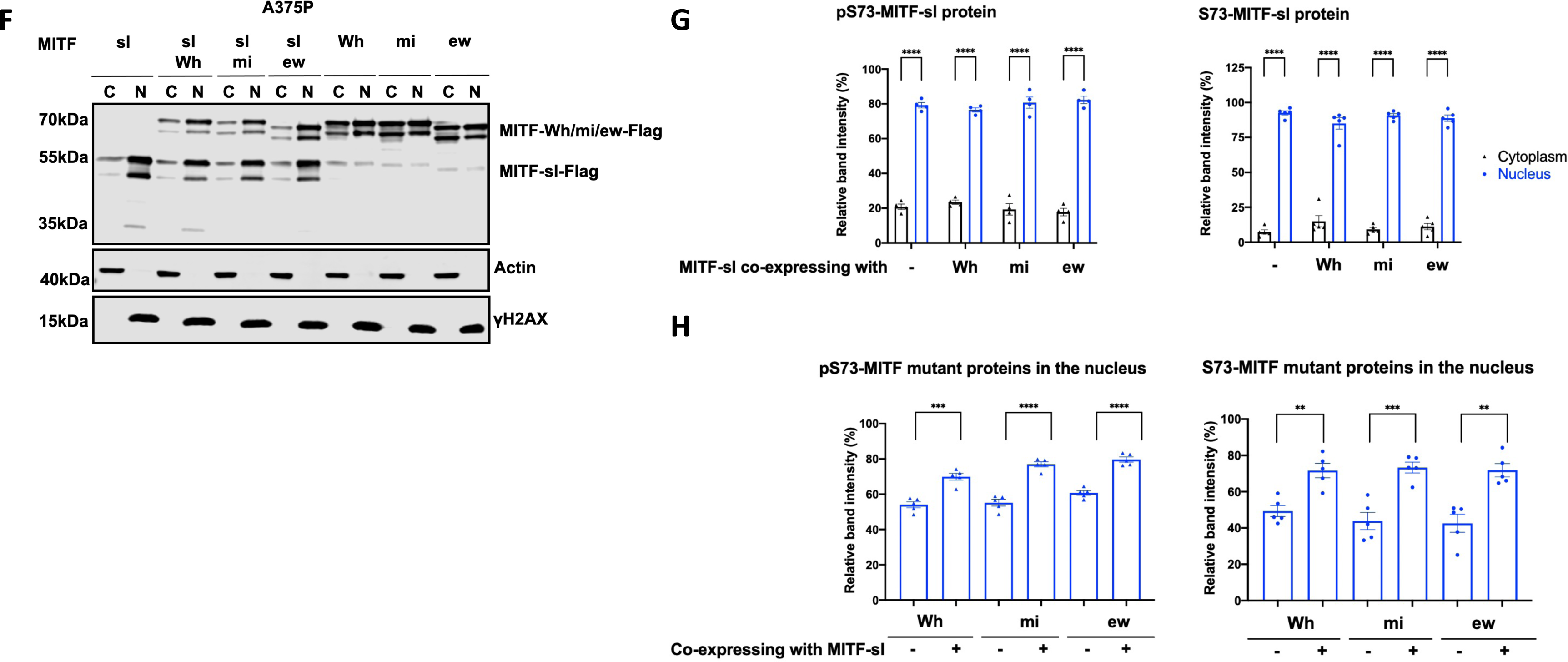
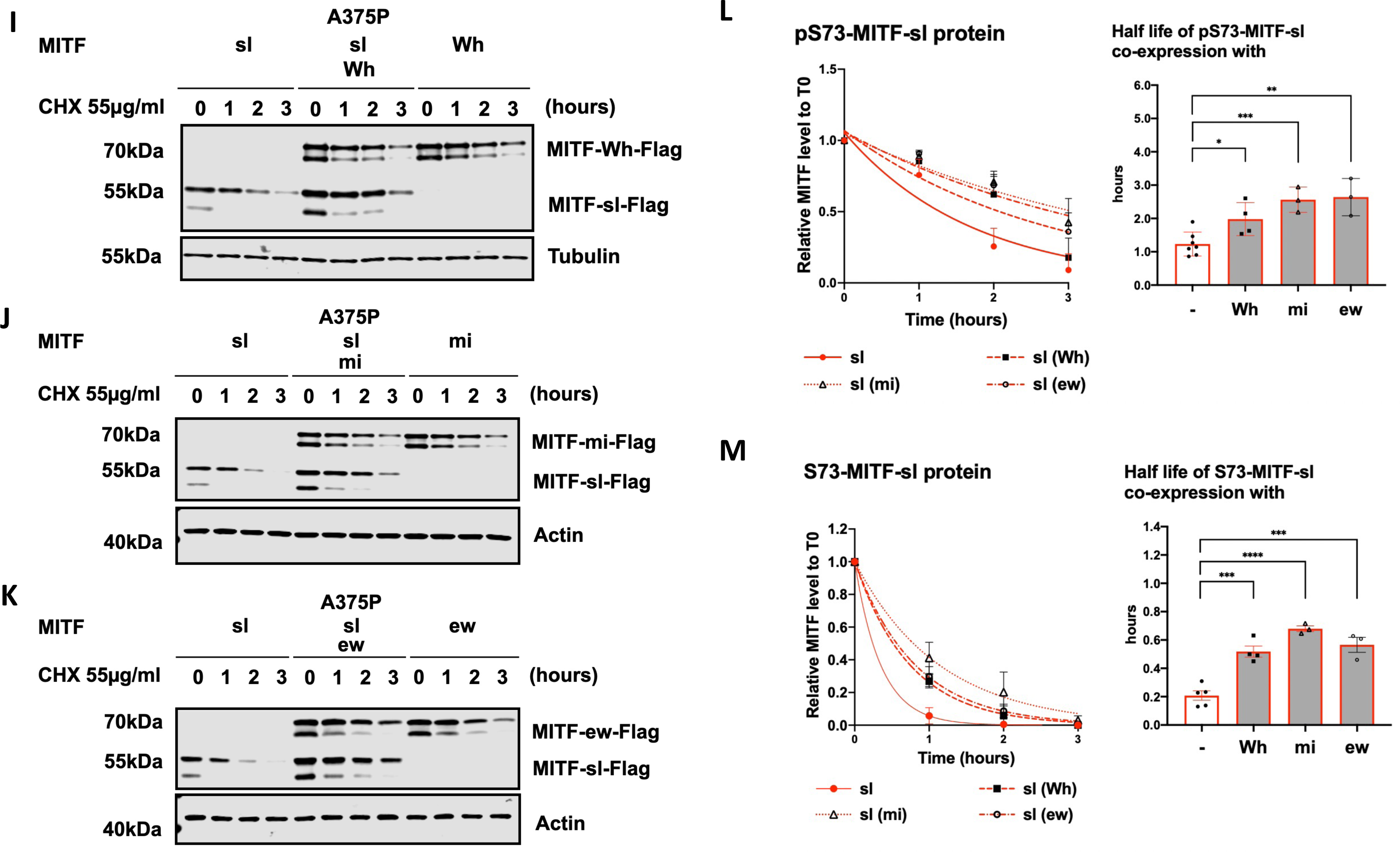
The carboxyl-domain of Mitf controls RNA and protein levels as well as its subcellular localization. **(A)** Western blot analysis of the Mitf-Flag proteins upon cycloheximide treatment. The dox-inducible A375P cells expressing the MITF-WT, MITF-sp, and MITF-sl proteins were treated with doxycycline for 24h to induce similar expression of the indicated mutant MITF proteins before treating them with 40 µg/ml cycloheximide (CHX) for 0, 1, 2, and 3 hours. The blots were stained using Flag antibody and protein quantitated using the Odyssey imager and ImageJ. Actin was used as a loading control. **(B), (C)** Non-linear regression (one-phase decay) and half-life analysis of the indicated pS73- and S73-MITF proteins over time after CHX treatment in A375P melanoma cells. The relative MITF protein levels to T0 were calculated, and non-linear regression analysis was performed. Error bars represent SEM of at least three independent experiments. Statistically significant differences (Student’s t-test) are indicated by *p< 0.05, ** p < 0.01, *** p < 0.001, **** p < 0.00, and ns not significant. **(D)** Western blot analysis of subcellular fractions isolated from A375P melanoma cells induced for 24 hours to overexpress different MITF mutant proteins. MITF-WT, MITF-Wh, MITF-sp, and MITF-sl protein in whole cell lysate (W), cytoplasmic (C), and nuclear (N) fractions were visualized using FLAG antibody. Actin and γH2AX were loading controls for cytoplasmic and nuclear fractions, respectively. **(E)** Intensities of the indicated pS73- and S73-MITF protein bands in the cytoplasmic and nuclear fraction from the western blot analysis in (E) were quantified separately with *ImageJ* software and are depicted as percentages of the total amount of protein present in the two fractions. Error bars represent SEM of three independent experiments. Statistically significant differences (Student’s t-test) are indicated by * p< 0.05, *p< 0.05, ** p < 0.01, *** p < 0.001, **** p < 0.00, and ns not significant. **(F)** Western blot analysis of subcellular fractions isolated from A375P cells transiently co-overexpressing the MITF*-*sl protein with the MITF-Wh, MITF-mi, and MITF-ew mutant MITF proteins. MITF proteins in cytoplasmic (C) and nuclear (N) fractions were visualized using FLAG antibody. Actin and γH2AX were loading controls for cytoplasmic and nuclear fractions, respectively. The MITF-sl protein migrates as a doublet at 50-55 kDa, whereas the other mutants migrate at 65-70 kDa. **(G)** The intensities of the pS73- and S73-MITF-sl protein in the cytoplasmic and nuclear fractions from western blot analysis (G) were quantified separately with ImageJ software and are depicted as percentages of the total protein present in the two fractions. Error bars represent SEM of at least three independent experiments. Statistically significant differences (Student’s t-test) are indicated by * p< 0.05, *p< 0.05, ** p < 0.01, *** p < 0.001, **** p < 0.00, and ns not significant. **(H)** Quantification of band intensities of the pS73- and S73-versions of the MITF-Wh, MITF-mi, and MITF-ew proteins as determined from western blot analysis (G) in the nuclear fractions of A375P cells transiently co-overexpressing the MITF*-*sl protein with the indicated MITF mutant proteins. The intensities are depicted as percentages of the total amount of protein present in the two fractions. Error bars represent SEM of at least three independent experiments. Statistically significant differences (Student’s t-test) are indicated by * p< 0.05, *p< 0.05, ** p < 0.01, *** p < 0.001, **** p < 0.00, and ns not significant. **(I), (J),** and **(K)** Western blot analysis of the degradation of the MITF-sl protein in the presence of non-DNA binding MITF mutations (MITF-Wh, MITF-mi, and MITF-ew). The A375P cells were transiently co-transfected with MITF-sl and either MITF-mi, MITF-ew, or MITF-Wh for 24 hours before being treated with 55 µg/ml CHX. The amount of MITF protein was then compared by western blot using FLAG antibody. Actin was used as a loading control. The band intensities were quantified using ImageJ software. **(L)** and **(M)** Non-linear regression (one-phase decay) and half-life analysis of the indicated pS73- and S73-MITF proteins over time after CHX treatment. The relative MITF protein levels to T0 were calculated, and non-linear regression analysis was performed. Error bars represent SEM of at least three independent experiments. Statistically significant differences (Student’s t-test) are indicated by * p< 0.05, *p< 0.05, ** p < 0.01, *** p < 0.001, **** p < 0.00, and ns not significant.

To determine the effect of MITF-sl on subcellular localization, we used dox-inducible A375P melanoma cells overexpressing MITF-Flag fusion proteins and performed cellular fractionation. After inducing expression of MITF for 24 hours, the nuclear and cytoplasmic fractions were separated as described^35, 36^, and MITF proteins were characterized by Western blotting. For the MITF-WT and MITF*-*sp proteins, both pS73 and S73 bands were observed at similar ratios in the nuclear and cytoplasmic fractions (Figures 2D and 2E). However, for MITF-sl, both pS73 and S73 bands were predominantly located in the nucleus (Figures 2D and 2E). The same results were observed when the MITF-WT, MITF-sp, and MITF-sl proteins were overexpressed in the 501Mel and SKmel28 melanoma cell lines (Figures S9A-D). Flag-tagging the MITF protein at the N-end or replacing the C-end Flag with GFP also resulted in a significantly increased nuclear presence of the MITF-sl protein (Figures S9E and S9F). Co-IP showed that nuclear accumulation of MITF-sl protein was not due to effects on interactions with 14-3-3 protein (Figure S9G), which has been shown to interact with MITF phosphorylated at S173 and lead to the retention of MITF in the cytosol in osteoclasts^37^. To determine if the six amino acids encoded by exon 6A were able to mediate nuclear localization, MITF lacking (-) or containing (+) this exon was transiently expressed in A375P cells. No difference was observed in the distribution of MITF between the nuclear and cytoplasmic fractions of the MITF-Wh and MITF-Wh(-) or MITF-sl(+) and MITF-sl constructs (Figures S9H and S9I). Taken together, we conclude that residues 316-419 of MITF, but not exon 6A or the tags, affect MITF subcellular localization.

### MITF^mi-sl^ translocates its Mitf partners into the nucleus and improves its own stability

Consistent with previous reports^34, 38^, stably expressed MITF-mi and MITF-ew proteins were primarily present in the cytoplasm (Figure S10A, compared with Figures 2D or 2E for MITF-WT or MITF-Wh proteins). To determine if the MITF-sl protein might affect the subcellular localization of the above non-DNA binding mutant MITF proteins, they were transiently co-overexpressed with the MITF*-*sl protein. As before, a significant portion of the pS73- and S73-MITF*-*sl proteins was observed in the nucleus (Figures 2F and 2G). In contrast to stably expressed proteins (Figure S10A), transiently expressed MITF-mi or MITF-ew showed equal distribution between cytoplasm and nucleus; no significant difference was noted between stable and transient expression of the MITF-Wh protein (Figures 2D-H). However, when co-expressed with the MITF*-*sl protein, they were all significantly translocated into the nucleus (Figures 2F-H). Similar results were observed in cells co-expressing MITF*-* sl and MITF-WT (Figures S10B-D). Our data, therefore, strongly suggest that the MITF*-*sl protein can dimerize with both mutant and WT proteins and induce nuclear localization of its partner by either translocating the dimer to the nucleus or keeping it from leaving the nucleus.

We assessed the stability of the MITF-sl protein in cells also expressing either MITF-Wh, MITF-mi, or MITF-ew. Interestingly, the stability of both pS73 and S73 MITF-sl was considerably increased in the presence of the MITF-Wh, MITF-mi, and MITF-ew proteins, with the most pronounced effect observed in cells also expressing MITF-mi and MITF-ew (around 2.5-fold increase for pS73 and 3.5-fold increase for S73) (Figures 2I-M). However, the stability of the MITF-Wh, MITF-mi, and MITF-ew dimeric partner proteins themselves remained unchanged upon co-expression of MITF-sl as compared to the condition when they were expressed in the absence of MITF-sl (Figures 2I-M). The stability of pS73 and S73 versions of MITF-sl was also significantly improved when co-expressed with MITF-WT (Figures S10E and S10F). To eliminate the possibility that we saturated the degradation machinery in the cells, we co-transfected the cells with MITF-sl-GFP and MITF-sl-Flag and measured the stability of MITF-sl-Flag protein. The results showed that the stability of MITF-sl-Flag was not affected by the presence of MITF-sl-GFP (Figure S10G). Taken together, our data suggest that the MITF-sl protein forms dimers with the MITF-Wh, MITF-mi, and MITF-ew proteins which then drags them into the nucleus or prevents them from leaving the nucleus, leading to increased stability of the MITF-sl protein itself (without, however, changing the stability of the partner proteins). On balance, this may increase the formation of DNA-binding MITF-sl homodimers after dissociation from their dimeric partners, and so explain the genetic suppression effect observed *in vivo*. The effects on protein stability and gene expression changes induced in the presence of MITF-sl alone might explain the hypomorphic effect in *Mitf^mi-sl^* homozygotes. That this hypomorphic effect is not seen in compound heterozygotes with the non-DNA binding mutants may be due to the balance between nuclear import/export, effects on stability, and the rate of dissociation of MITF-sl from its dimeric partner and subsequent effects on transcription.

### Effects on nuclear localization and stability are encoded in the carboxyl-domain

To determine which regions within the C-end of MITF contain its nuclear retention properties, we generated truncated versions of MITF-sp with Flag-tag fusion at the C-end in our inducible vector system (schematic diagram in Figure 3A). The S73-MITF-WT and S73-MITF-sp-Δ326-377 proteins are distributed equally between the cytoplasmic and nuclear fractions; the pS73-MITF-sp-Δ326-377 was slightly more cytoplasmic (Figures 3B and 3C). In contrast, a significant portion of the MITF-sp-326* and MITF-sp-378* proteins was present in the nuclear fraction (Figures 3B and 3C), suggesting that the 378-419 domain, including the phosphorylation sites indicated in Figure 3A, plays an essential role in controlling the nuclear localization of MITF. Interestingly, the non-phosphorylatable alanine mutation at S409 led to slightly more nuclear localization of the pS73 MITF form, whereas the single S384A, S397A, S401A, and S405A mutations did not alter MITF nuclear localization (Figure S11A and S11B). However, the quadruple S397/401/405/409A mutation in MITF-sp (MITF-sp-4A) or MITF-WT (MITF-WT-4A) led to increased nuclear localization of the respective proteins (Figure S11A and S11B). This suggests that the phosphorylation cascade at the C-end may be involved in the cytoplasmic retention of MITF but that other elements within the C-end must also be involved.

**Figure 3:**
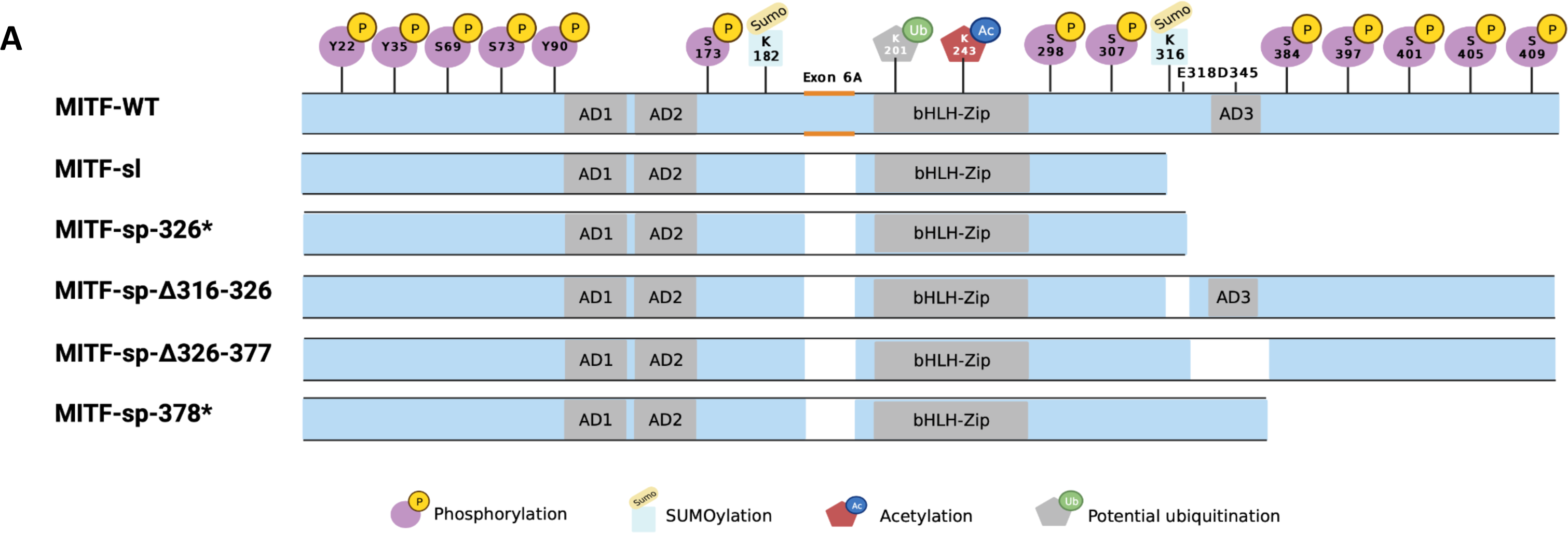
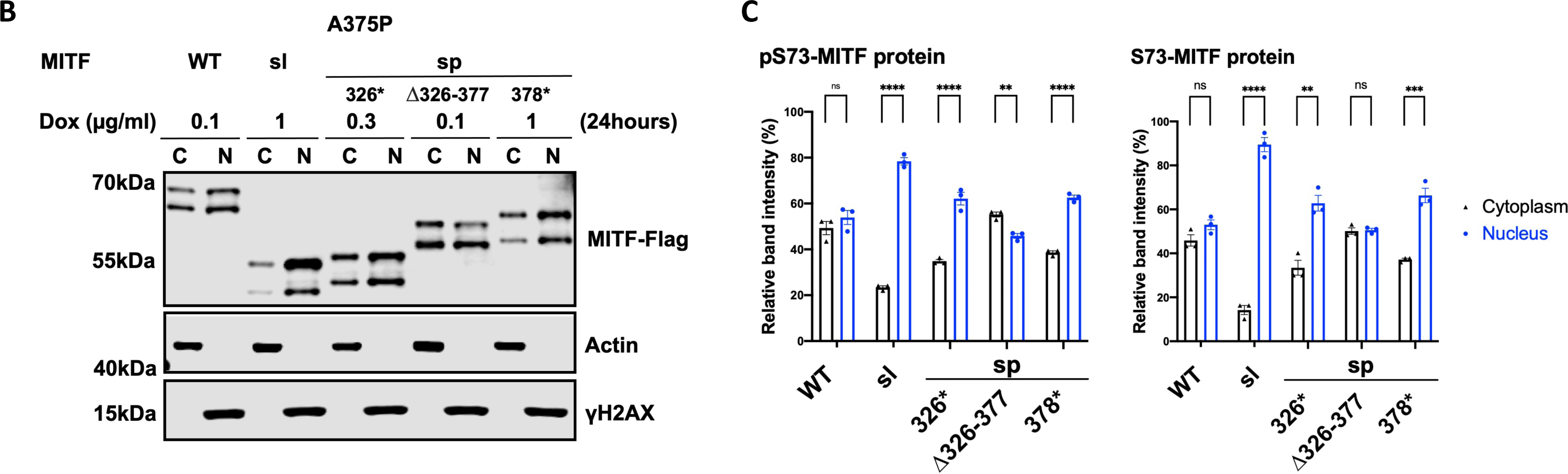
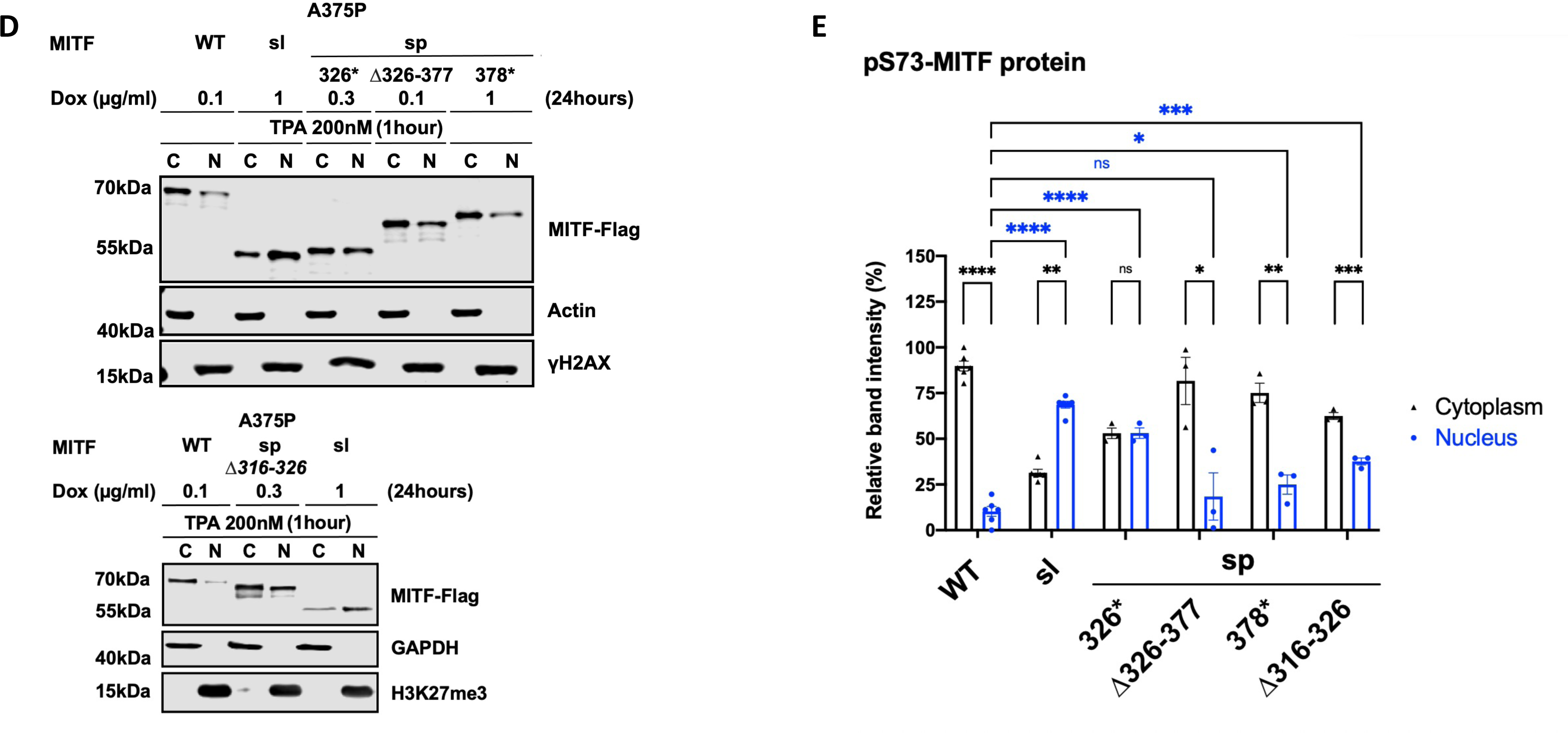
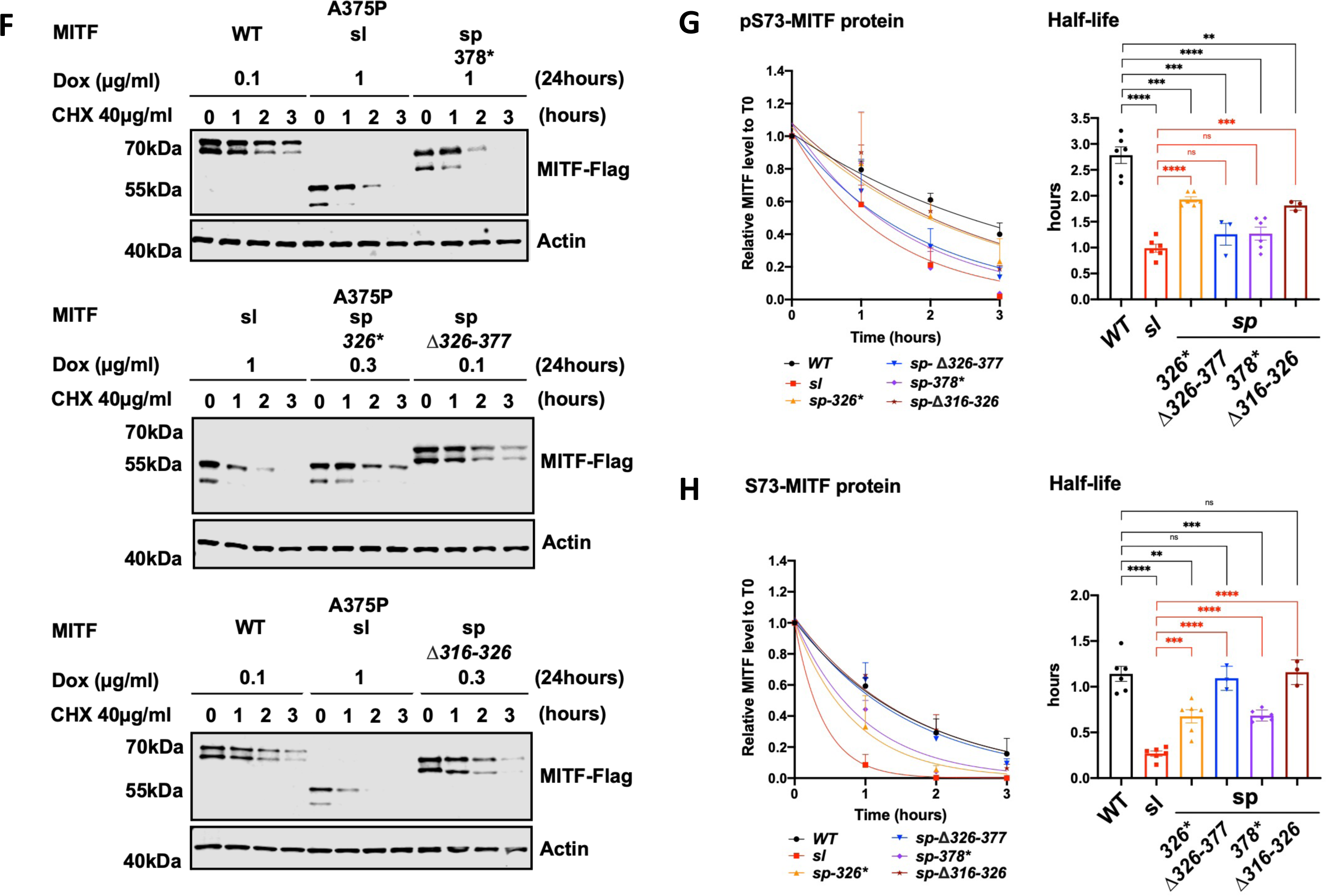
The carboxyl-domains of Mitf control its nuclear localization and stability. **(A)** Schematic of MITF-sp truncation constructs. C-term truncations were generated by introducing stop codons at position Q326 or L378 or by deleting fragments 326-377 or 316-326. MITF-sp-326* introduces a stop-codon at residue 326 and therefore contains the SUMO-site at 316; MITF-sp-Δ326-377 lacks the tentative activation domain AD3; MITF-sp-Δ316-326 lacks the SUMO-site and adjacent amino acids; MITF-sp-378* lacks the series of phosphorylation sites at the carboxyl-end of the protein. **(B)** Western blot analysis of subcellular fractions isolated from A375P melanoma cells induced to overexpress the different MITF mutant proteins fusioned with Flag-tag at C terminus for 24 hours. MITF-WT, MITF-sl, MITF-sp-326*, MITFmi-sp-378*, and MITF-sp-Δ326-377 in cytoplasmic **(C)** and nuclear (N) fractions were visualized using FLAG antibody. Actin and γH2AX were loading controls for cytoplasmic and nuclear fractions, respectively. **(C)** The intensities of the indicated pS73 MITF and S73 MITF proteins from the cytoplasmic and nuclear fractions of the western blot analysis in (B) were quantified separately with ImageJ software and are depicted as percentages of the total amount of protein present in the two fractions. Error bars represent SEM of three independent experiments. Statistically significant differences (Student’s t-test) are indicated by * p< 0.05, *p< 0.05, ** p < 0.01, *** p < 0.001, **** p < 0.00, and ns not significant. **(D)** Western blot analysis of subcellular fractions isolated from A375P melanoma cells induced for 24 hours to overexpress the different MITF mutant proteins before treatment with TPA at 200nM for 1 hour. MITF-WT, MITF-sl, MITF-sp-326*, MITF-sp-Δ326-377, MITF-sp-Δ316-326, and MITF-sp-378* protein in cytoplasmic (C) and nuclear (N) fractions were visualized using FLAG antibody. Actin or GAPDH and γH2AX or H3K27me3 were loading controls for cytoplasmic and nuclear fractions, respectively. **(E)** Intensities of the indicated pS73-MITF proteins from the western blot analysis in (D) in the cytoplasmic and nuclear fractions from the cell treated with TPA were quantified separately with ImageJ software and are depicted as percentages of the total amount of protein present in the two fractions. Error bars represent SEM of three independent experiments. Statistically significant differences (Student’s t-test) are indicated by * p< 0.05, *p< 0.05, ** p < 0.01, *** p < 0.001, **** p < 0.00, and ns not significant. **(F)** Western blot analysis of the MITF proteins from dox-induced A375P cells after treating them with 40 µg/ml CHX for 0, 1, 2, and 3 hours. The MITF proteins were visualized by western blot using FLAG antibody. Actin was used as a loading control. The band intensities were quantified using ImageJ software. **(G)** and **(H)** Non-linear regression (one-phase decay) and half-life analysis of the indicated pS73- and S73-MITF proteins over time after CHX treatment. The MITF protein levels relative to T0 were calculated, and non-linear regression analysis was performed. Error bars represent SEM of at least three independent experiments. Statistically significant differences (Student’s t-test) are indicated by * p< 0.05, *p< 0.05, ** p < 0.01, *** p < 0.001, **** p < 0.00, and ns not significant.

The MITF-sl protein was more nuclear than the MITF-sp-326* and MITF-sp-378* proteins (Figures 3B and 3C), suggesting that the 316-326 domain must also be involved in regulating nuclear localization. However, the MITF-sp-Δ316-326 construct, which lacks the K316 SUMO-site and adjacent residues, did not alter the cytoplasmic-nuclear distribution of MITF (Figures S11C and S11D). This suggests that residues 326-419 contain a major signal for mediating nuclear export of MITF and that residues 316-326 also contribute. Intriguingly, the MITF-mi and MITF-ew proteins containing the 316*, 378*, or Δ316-326 mutations were more nuclear than their full-length counterparts (Figure S11E and S11F).

Previous work has shown that treatment with 12-*O*-tetradecanoylphorbol-13-acetate (TPA), an agent known to induce ERK kinase activity, leads to phosphorylation of S73 of MITF and shifts the protein to the cytoplasm^10^. Consistent with that, TPA treatment promoted S73 phosphorylation (as seen by the almost exclusive presence of the upper MITF-band) of MITF-WT and shifted the protein out of the nucleus (Figures 3D and 3E). The MITF-sl protein was also phosphorylated at S73 but, as before, it mostly stayed in the nucleus. The MITF-sp-Δ326-377, MITF-sp-Δ316-326, and MITF-sp-378* proteins were also phosphorylated at S73, but a large proportion of these proteins was located in the cytoplasm after TPA treatment. The MITF-sp-326* protein was equally distributed between the two compartments (Figures 3D and 3E). These data suggest that MITF has more than one nuclear export domain, one involving S73 and S69 and another located in the C-end, which may act independently. The LocNES algorithm^39^ predicts a couple of nuclear export signals (NESs) in the C-end of MITF spanning residues 350-366 (NES1) and 374-388 (NES2). To test their role, we generated a fusion of MITF-sl to either NES1 or NES2 or both NES1 and NES2 (schematic diagram in Figure S11G) in our inducible vector system and performed cell fractionation. All the fusions slightly increased the proportion of MITF-sl in the cytoplasm, regardless of S73 phosphorylation status (Figures S11H and S11I). Upon TPA treatment, pS73-MITF-sl-NES1, -NES2, and NES1-NES2 were significantly exported to the cytoplasm, as opposed to pS73-MITF-sl (Figures S11J and S11K). Our findings suggest that NES1 and NES2 are involved either in nuclear export of MITF or in blocking its import.

To determine which regions within the C-end of MITF are essential for mediating effects on stability, we performed protein stability assays using the MITF deletion constructs in the presence of CHX. The results showed that, again, the pS73 form of MITF-WT, MITF-sl, MITF-sp-326*, MITF-sp-Δ316-326, and MITF-sp-378* proteins was considerably more stable than the corresponding S73 proteins (Figures 3F-H). It also showed that the MITF-sp-326*, MITF-sp-378*, and MITF-sp-Δ316-326 proteins were less stable than the MITF-WT protein. However, MITF-sl still showed the most rapid degradation upon CHX treatment of all proteins tested (Figures 3F-H). The results suggest that the 316-326 and 378-419 domains are important for nuclear localization and MITF stability. The effects of the 378-419 domain on MITF localization are not due to a single phosphorylation site at the C-end since S384A, S397A, S401A, S405A, and S409A did not significantly affect the stability of MITF-sp, nor did their combination in the 4A mutant construct (Figures S11L and S11M).

To further investigate the role of the 316-419 domain in mediating MITF protein stability, we determined the stability of the non-DNA binding MITF-mi, MITF-mi-316*, MITF-ew, and MITF-ew-316* proteins. Although the pS73-MITF-mi and pS73-MITF-ew proteins were slightly more stable than pS73-MITF-WT, the stability of S73-MITF-mi and S73-MITF-ew did not significantly differ from MITF-WT (Figures S11N and S11O). Meanwhile, the double mutant proteins (i.e., MITF-mi-316* and MITF-ew-316*) exhibited increased presence in the nucleus (Figure S11E) yet had similar stability as MITF-WT (Figures S11P and S11Q). This suggests that the ability to bind to DNA in concert with MITF C-end might be important for controlling MITF stability and triggering the degradation process.

### MITF is mainly degraded through a ubiquitin-mediated proteasome pathway in the nucleus

To determine which degradation pathway is responsible for MITF degradation, we treated the cells with the ubiquitin-proteasomal inhibitor MG132 and the lysosomal inhibitor Baf-A1 together with CHX for 3 hours. Treatment with MG132 and CHX increased the stability of the MITF-WT, MITF-sp, and MITF-sl proteins, whereas Baf-A1 and CHX treatment did not (Figures 4A and 4B). Treatment with MG132 or Baf-A1 without CHX showed a significant increase in the intensity of the pS73 band of the MITF-sp and MITF-sl proteins, though not MITF-WT protein (Figures 4C and 4D). Interestingly, the S73 bands of MITF-WT, MITF-sp, and MITF-sl showed a considerable increase after MG132 treatment (approximately 2.4-, 2.7-, and 6.7-fold increase respectively), but the increase was much less pronounced or even non-significant (in the case of S73-MITF-sl) upon Baf-A1 treatment (Figure 4D). This suggests that the ubiquitin-proteasome pathway is the primary degradation machinery for MITF.

**Figure 4:**
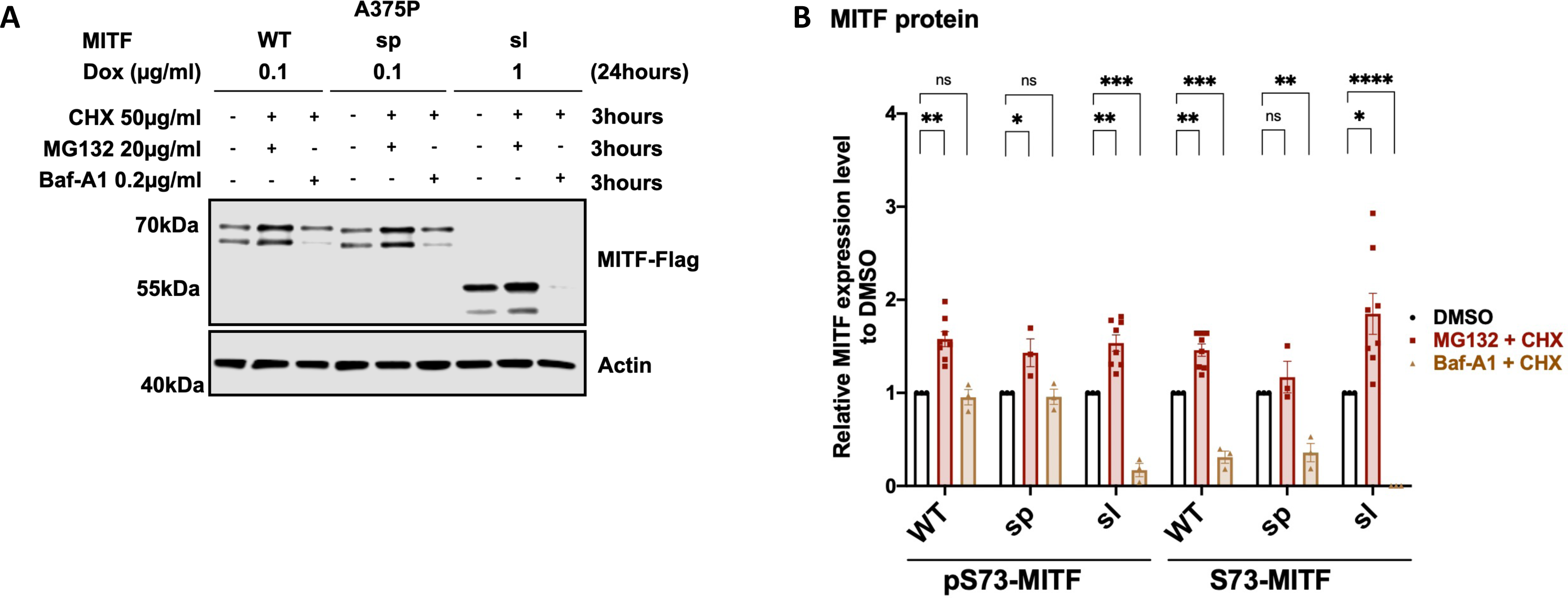
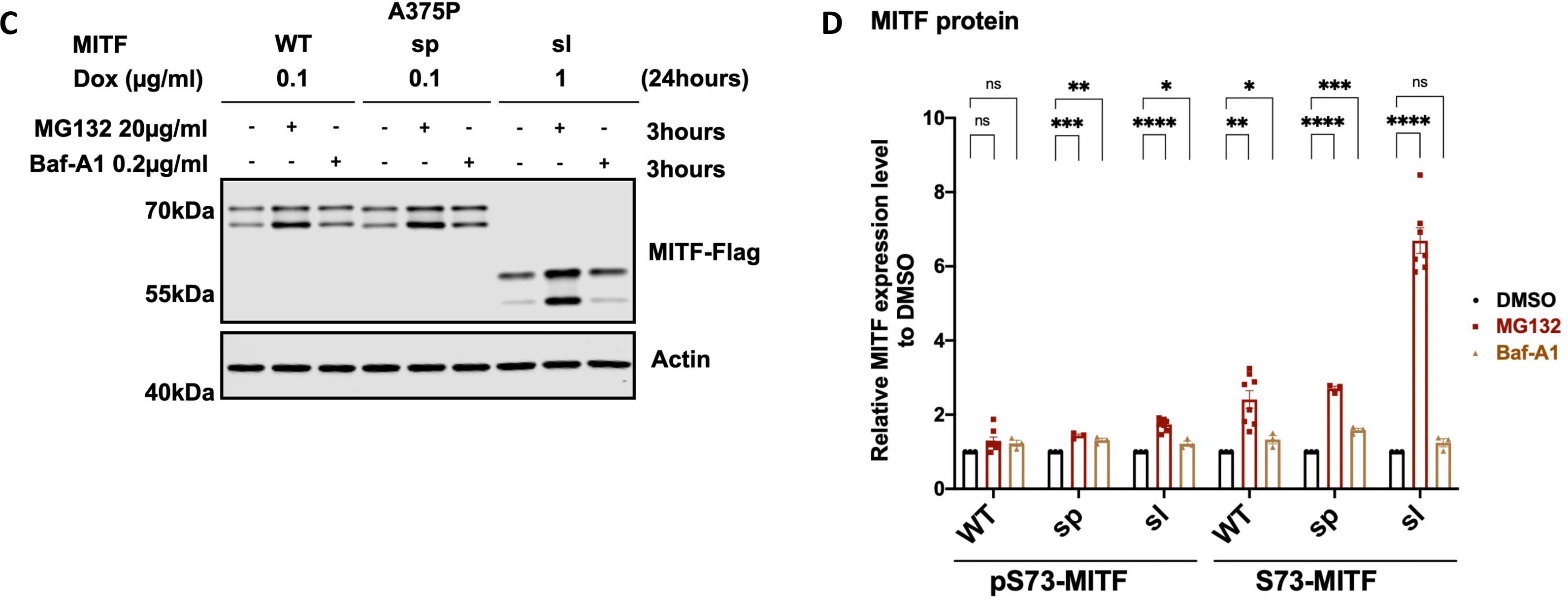
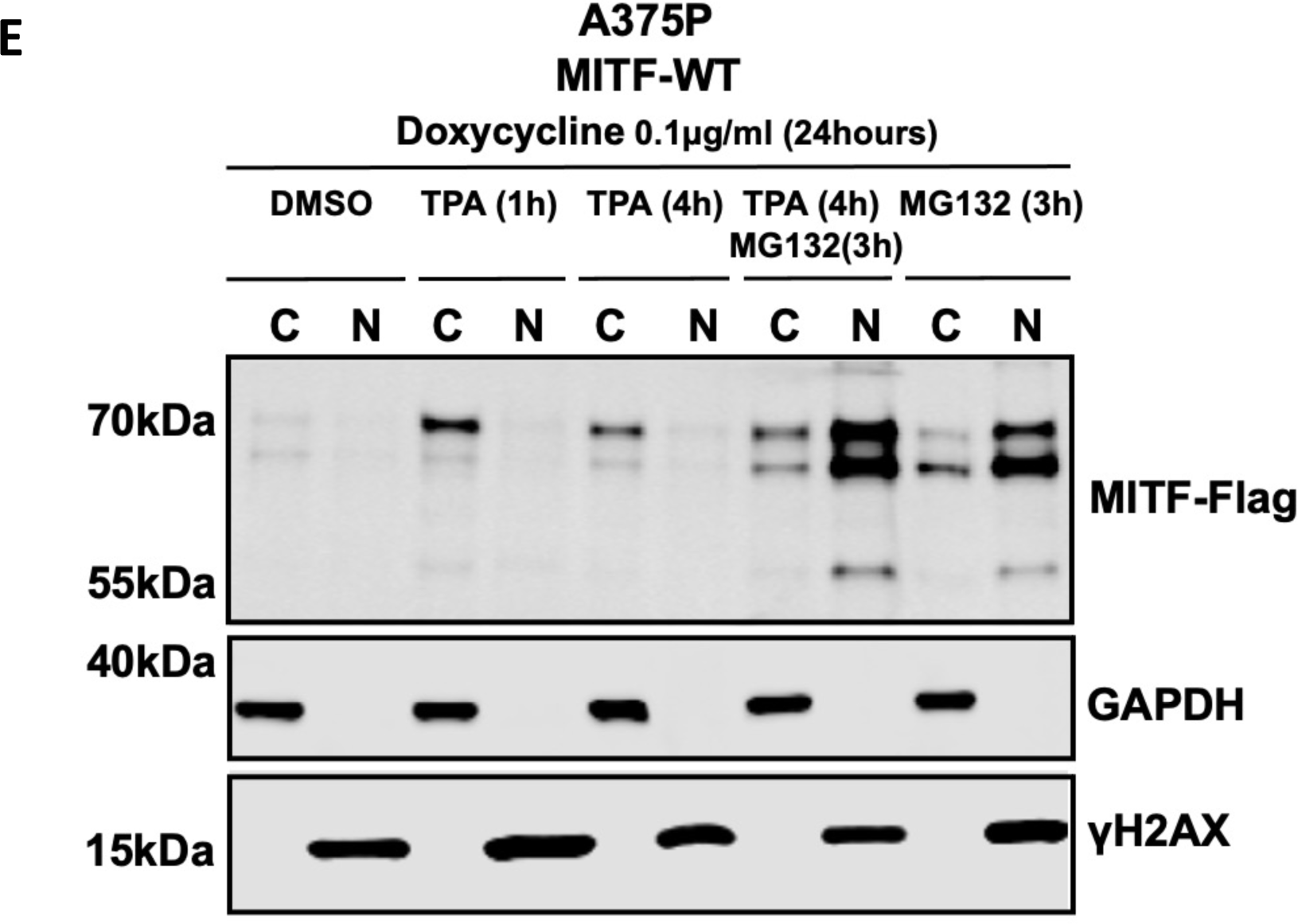
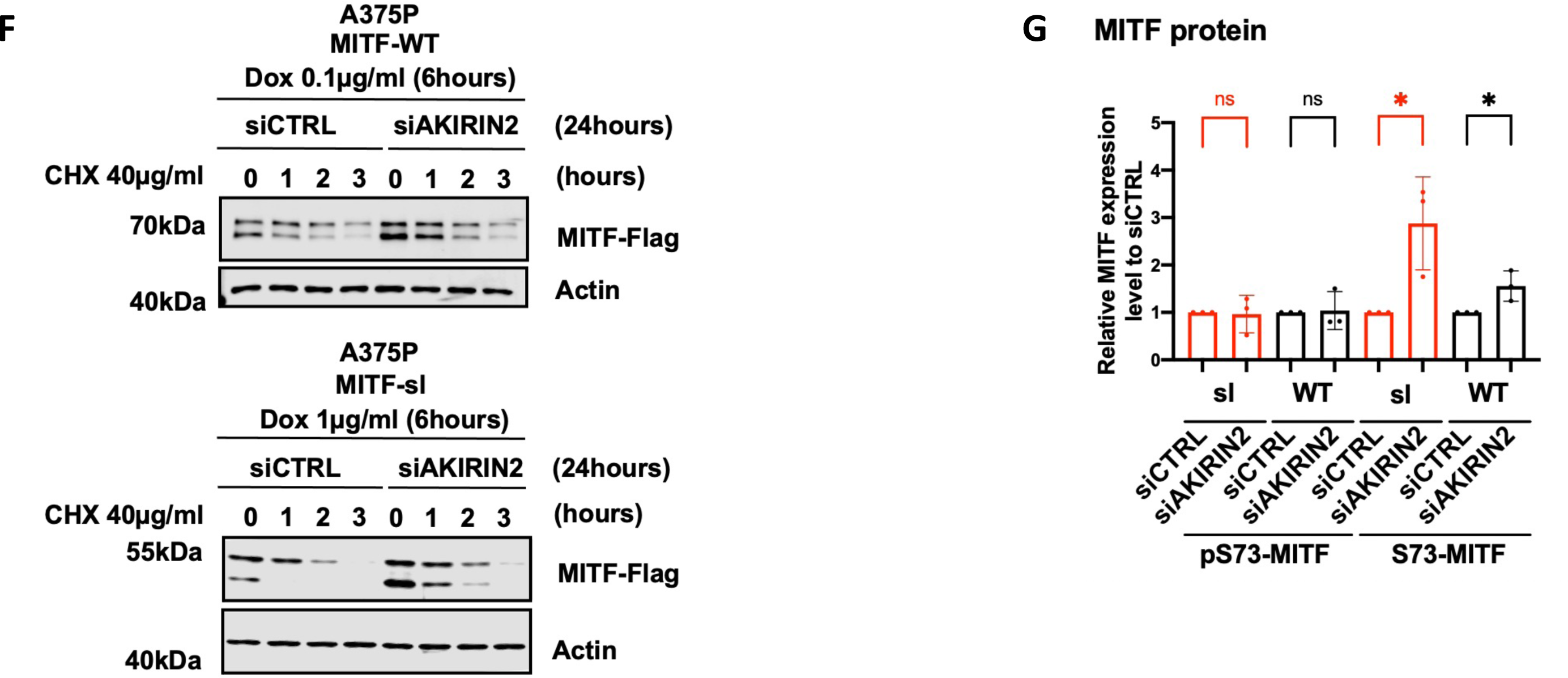
MITF is mainly degraded through the proteasome pathway in the nucleus. **(A)** Western blot analysis of the MITF-WT, MITF-sp, and MITF-sl proteins. Expression was induced for 24 hours in A375P cells treated with 50 µg/ml CHX in the presence of either DMSO or 40 µg/ml MG132 or 0.1 µg/ml Baf-A1 for 3 hours. The MITF protein was then visualized by western blot using FLAG antibody. Actin was used as a loading control. The band intensities were quantified using ImageJ software. **(C)** Western blot analysis of the MITF-WT, MITF-sp, and MITF-sl proteins. Expression was induced for 24 hours in A375P cells treated with either DMSO or 40 µg/ml MG132 or 0.1 µg/ml Baf-A1 for 3 hours. The MITF protein was then visualized by western blot using FLAG antibody. Actin was used as a loading control. The band intensities were quantified using ImageJ software. **(B)** and **(D)** The indicated pS73- and S73-MITF protein band intensities from western blot analysis (A) and (C), respectively, were quantified separately with ImageJ software and are depicted relative to DMSO. Error bars represent SEM of at least three independent experiments. Statistically significant differences (Student’s t-test) are indicated by * p< 0.05, *p< 0.05, ** p < 0.01, *** p < 0.001, **** p < 0.00, and ns not significant. **(E)** Western blot analysis of subcellular fractions isolated from A375P melanoma cells induced to overexpress MITF-WT protein before treating with either 200nM TPA for 1 or 4 hours or 40 µg/ml MG132 for 3 hours or 200nM TPA for 1 hour and then adding 40 µg/ml MG132 for the next 3 hours. MITF-WT protein in cytoplasmic (C) and nuclear (N) fractions were visualized using FLAG antibody. GAPDH and γH2AX were loading controls for cytoplasmic and nuclear fractions, respectively. **(F)** Western blot analysis of the stability of the MITF-WT and MITF-sl mutant proteins after knocking down *AKIRIN2*, a key regulator of the nuclear import of proteasomes, for 24 hours and then inducing MITF expression using dox for 6 hours. The inducible A375P cells were treated with 40 µg/ml CHX for 0, 1, 2, and 3 hours. The MITF proteins were then visualized by western blot using FLAG antibody. Actin was used as a loading control. The band intensities were quantified using ImageJ software. **(G)** The intensities of the indicated pS73- and S73-MITF protein bands were quantified from western blot analysis in (F) with ImageJ software and are depicted as relative protein expression to DMSO. Error bars represent SEM of three independent experiments. Statistically significant differences (Student’s t-test) are indicated by * p< 0.05, *p< 0.05, ** p < 0.01, *** p < 0.001, **** p < 0.00, and ns not significant.

To test whether the proteasomal degradation pathway takes place predominantly in the nucleus or the cytoplasm, we treated MITF-WT expressing cells with both MG132 and TPA. As shown in Figure 4E, TPA treatment significantly increased the total MITF protein compared to vehicle controls, suggesting that shutting the protein out of the nucleus increases stability. Treating the cells for 3 hours with MG132 in the presence of TPA revealed a significant increase of MITF, primarily in the nucleus (Figure 4E).

Dox-inducible A375P melanoma cells expressing MITF-WT, MITF-sp, and MITF-sl were exposed to CHX and the nuclear export inhibitor leptomycin B (LMB)^40^ for different time points before harvesting for Western blotting. The results showed that the stability of pS73-MITF-WT and pS73-MITF-sp was significantly reduced upon LMB treatment, whereas the stability of S73 was not changed, and the stability of both pS73- and S73-MITF-sl was decreased upon LMB treatment (Figures S12A and S12B). AKIRIN2 is essential for proteasomal degradation in the nucleus, and we hypothesized that its removal would increase MITF stability^41^. We thus knocked down *AKIRIN2* in our dox-inducible A375P cells expressing MITF-WT and MITF-sl prior to CHX treatment. After treating the cells for 24 hours with siAKIRIN2, the expression of the mRNA *AKIRIN2* was significantly decreased (Figure S12C), and the expression of both S73-MITF-WT and S73-MITF-sl was significantly increased (Figures 4F, 4G, and S12D). Our results show that MITF is degraded in the nucleus through the proteasomal pathway. The increased nuclear presence of the MITF-sl protein may explain its reduced stability.

### The K316R and E318K mutations together with the S409A mutation reduce MITF stability and increase its nuclear presence

To determine if the SUMOylation site at K316 was involved in mediating MITF subcellular localization, we replaced the K316 residue with arginine in MITF-WT and MITF-sp. We also determined the effects of the E318K mutation since individuals carrying this mutation in MITF are predisposed to melanoma, and the mutation abolishes SUMOylation at K316^16, 17^. Alone, neither the K316R nor the E318K mutations altered the localization of the MITF-WT or MITF-sp proteins (Figures 5A-C). However, the double mutant proteins K316R-S409A and E318K-S409A were more nuclear, regardless of the S73-phosphorylation status (Figures 5A-C). Critically, both double mutants were able to override the effects of TPA on nuclear export, resulting in equal distribution between the nucleus and cytoplasm (Figures 5D and 5E). However, the K316R and E318K mutations did not alter the nuclear localization of MITF or nuclear export upon TPA treatment when together with the S384A, S397A, S401A, or S405A mutations, (Figures S13A and S13B). Taken together, this suggests that a specific interaction between the SUMOylation site at K316 and the phosphorylation site at S409 is important for mediating MITF export. However, since these double mutants do not fully replicate the effects of the MITF-sl protein on localization, additional regions within the C-end must be important as well.

**Figure 5:**
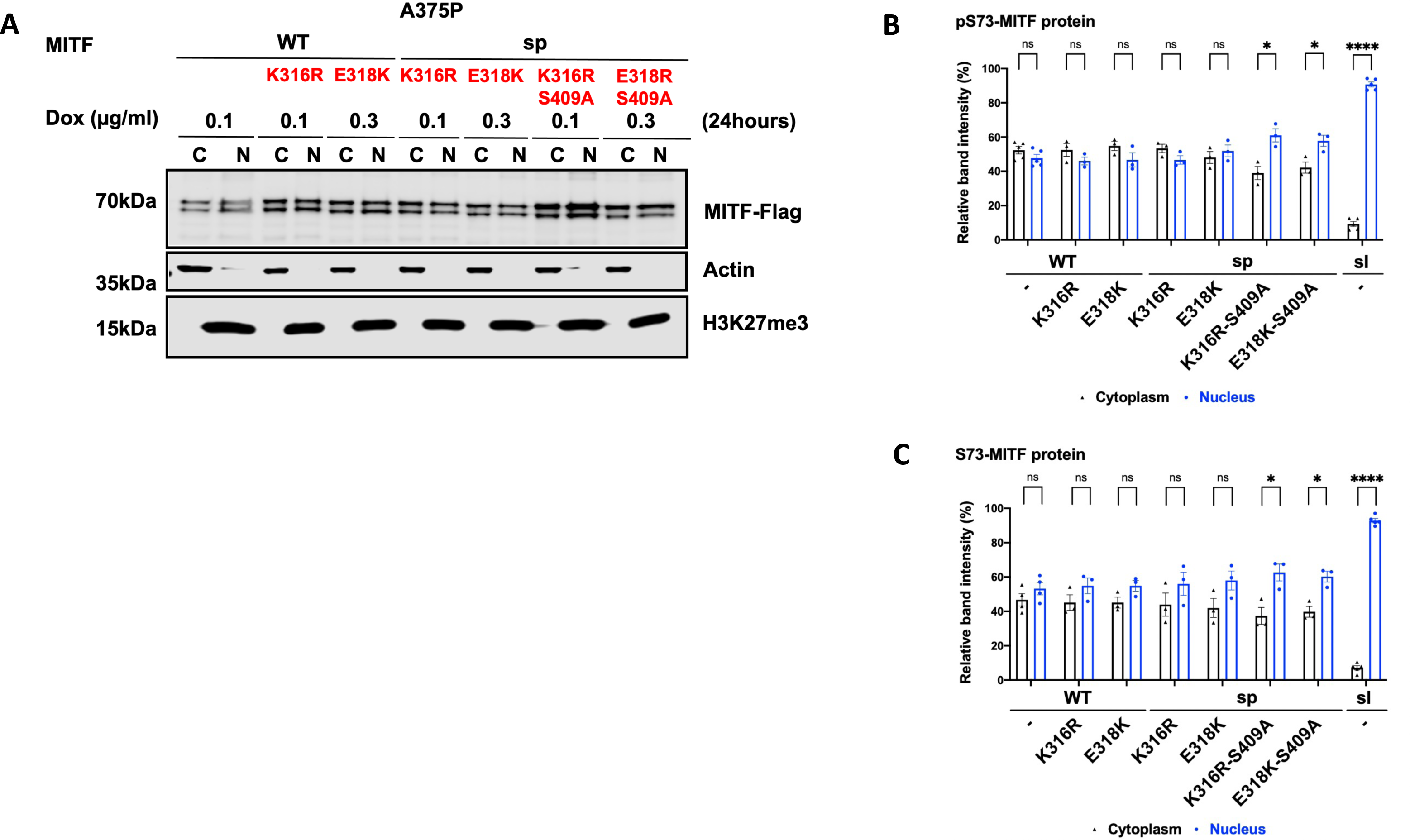
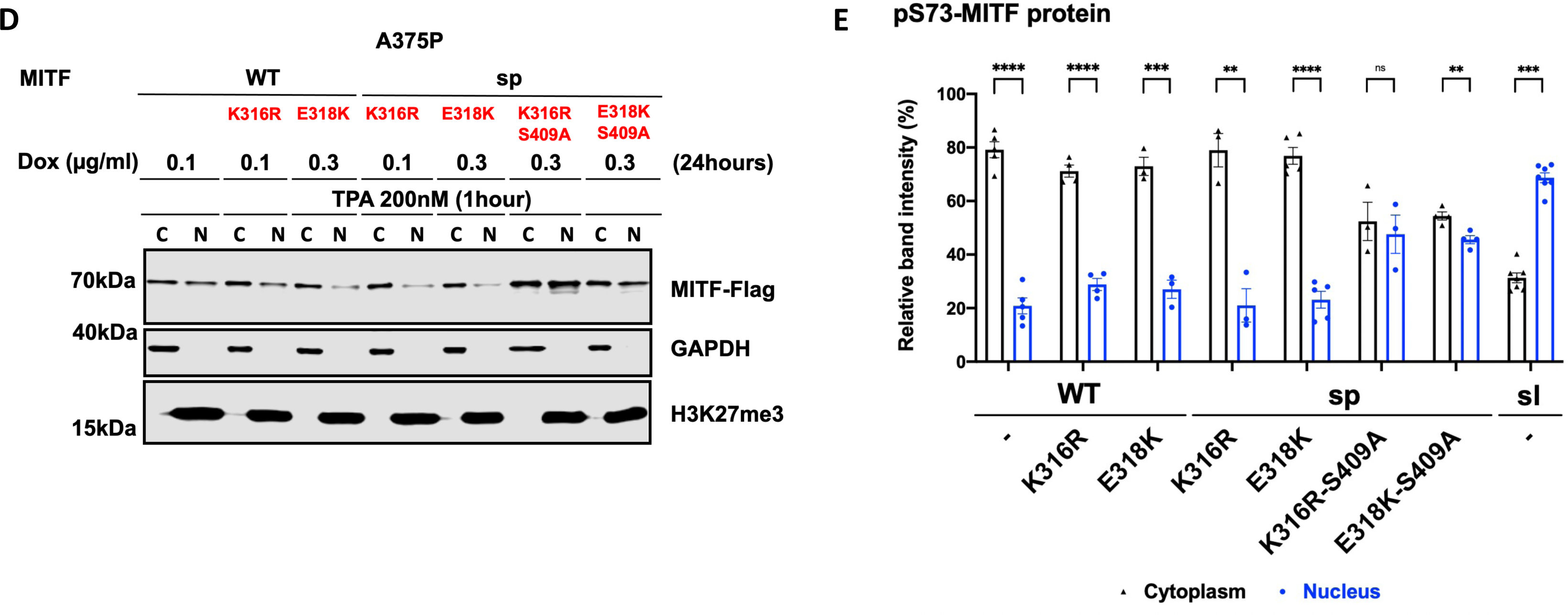
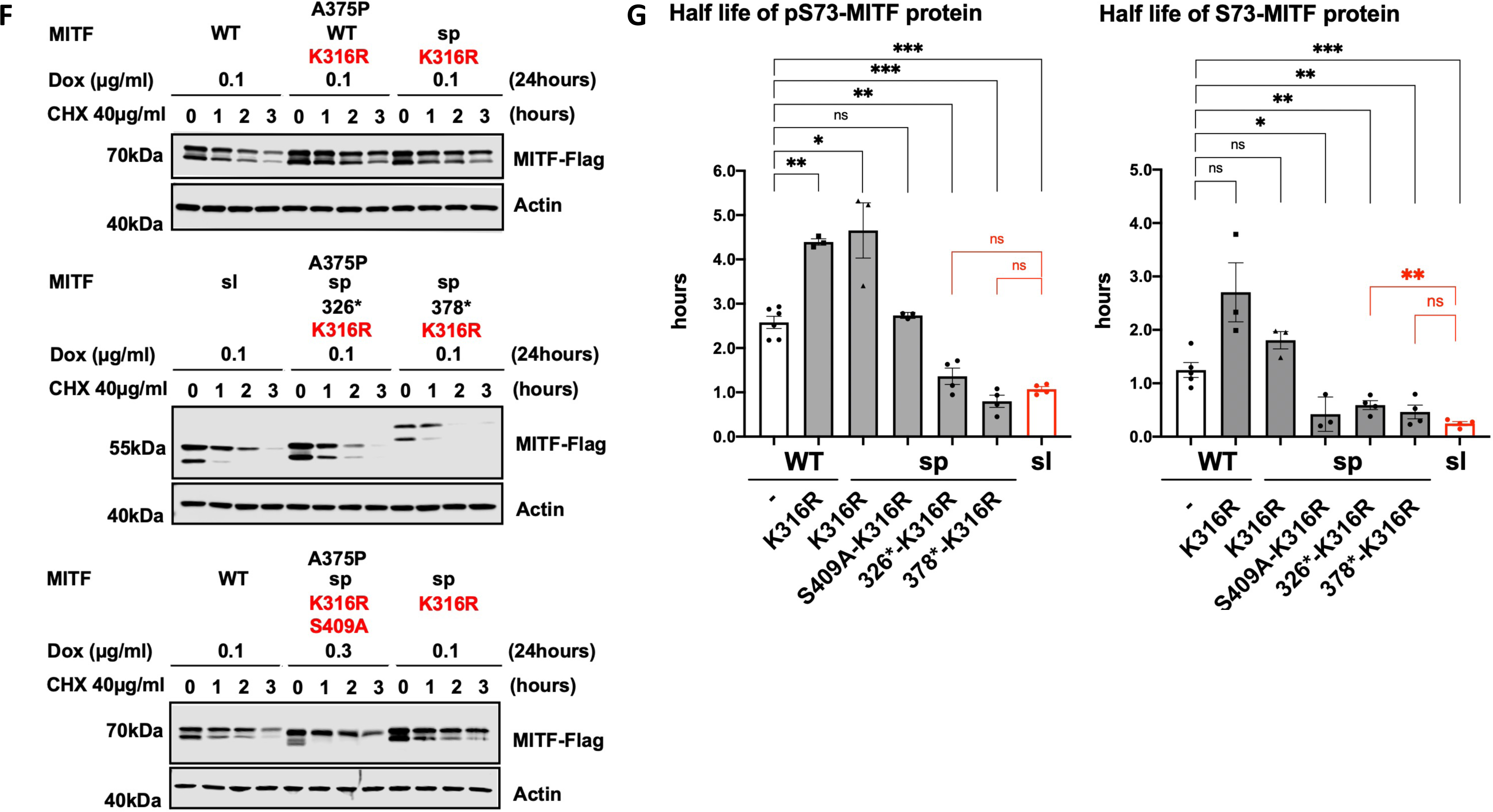
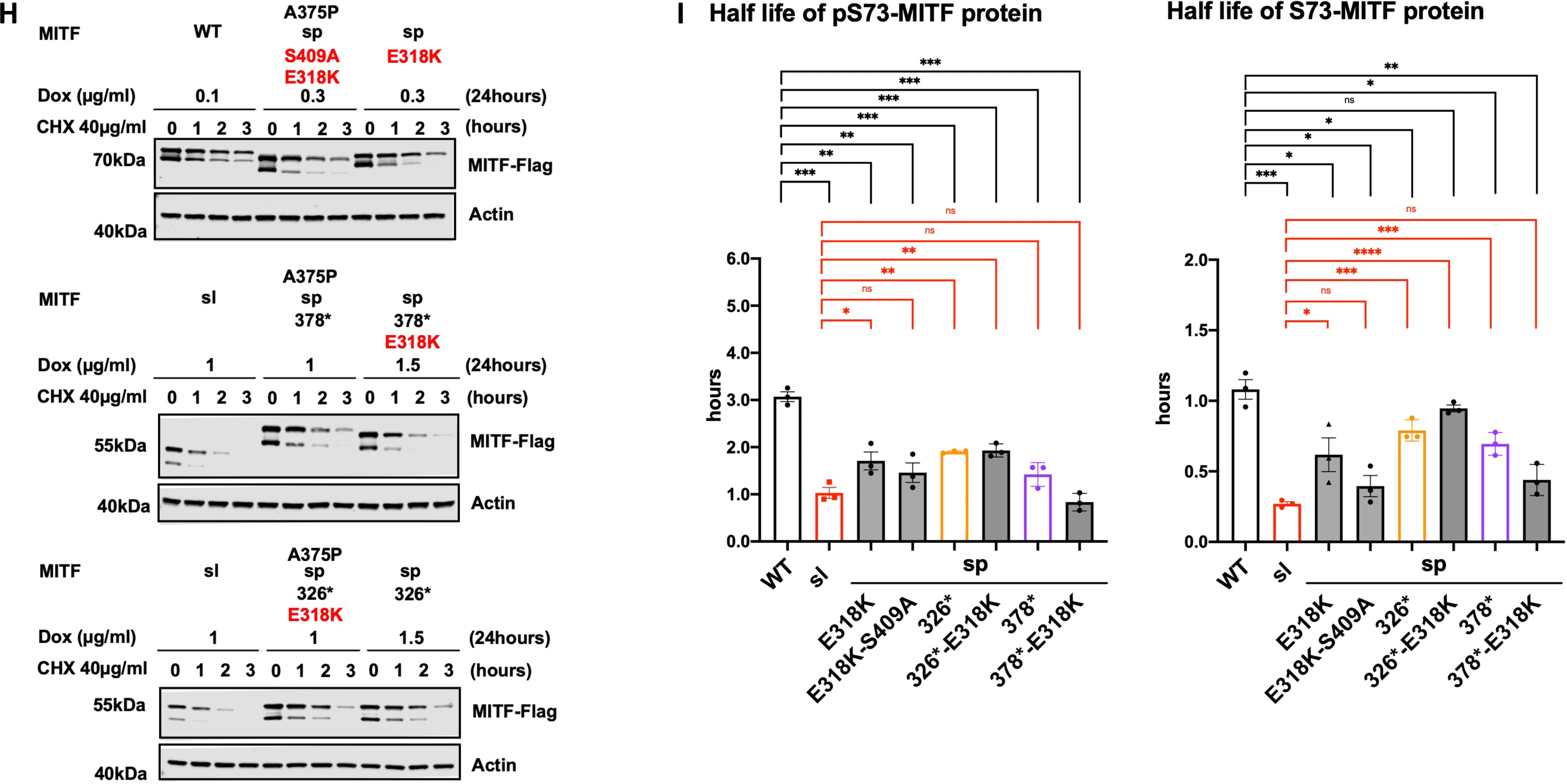
The interplay between SUMOylation at K316 and phosphorylation site at S409 in regulating MITF protein stability and localization. **(A)** Western blot analysis of subcellular fractions isolated from A375P melanoma cells induced for 24 hours to overexpress the indicated MITF mutant proteins. The MITF proteins in cytoplasmic (C) and nuclear (N) fractions were visualized using FLAG antibody. Actin or GAPDH and H3K27me3 were loading controls for cytoplasmic and nuclear fractions, respectively. **(B)** and **(C)** The intensities of the indicated pS73- and S73-MITF proteins in the cytoplasmic and nuclear fractions from western blot analysis in (A) were quantified separately with ImageJ software and are depicted as percentages of the total amount of protein present in the two fractions. Error bars represent SEM of three independent experiments. Statistically significant differences (Student’s t-test) are indicated by * p< 0.05, *p< 0.05, ** p < 0.01, *** p < 0.001, **** p < 0.00, and ns not significant. **(D)** Western blot analysis of subcellular fractions isolated from A375P melanoma cells induced for 24 hours to overexpress the indicated MITF mutant proteins before treatment with 200nM TPA for 1 hour. The mutant MITF proteins in cytoplasmic (C) and nuclear (N) fractions were visualized using FLAG antibody. GAPDH and H3K27me3 were loading controls for cytoplasmic and nuclear fractions, respectively. **(E)** The intensities of the indicated pS73-MITF proteins bands in the cytoplasmic and nuclear fractions of the western blot analysis in (D) and (F), respectively, were quantified separately with ImageJ software and are depicted as percentages of the total amount of protein present in the two fractions. Error bars represent SEM of three independent experiments. Statistically significant differences (Student’s t-test) are indicated by * p< 0.05, *p< 0.05, ** p < 0.01, *** p < 0.001, **** p < 0.00, and ns not significant. **(F)** and **(H)** Western blot analysis of the stability of the MITF proteins. The inducible A375P cells were treated with doxycycline for 24h to express the indicated mutant MITF proteins before treating them with 40 µg/ml CHX for 0, 1, 2, and 3 hours. The MITF protein was then compared by western blot using FLAG antibody. Actin was used as a loading control. The band intensities were quantified using ImageJ software. **(G) and (I)** Half-life analysis of the pS73- and S73-MITF proteins over time after CHX treatment. The MITF protein levels relative to T0 were calculated, and non-linear regression analysis was performed. Error bars represent SEM of at least three independent experiments. Statistically significant differences (Student’s t-test) are indicated by * p< 0.05, *p< 0.05, ** p < 0.01, *** p < 0.001, **** p < 0.00, and ns not significant.

To clarify the role of the SUMOylation site on MITF protein stability, we tested the stability of the MITF-WT, MITF-sp, MITF-sp-378*, MITF-sp-326* proteins, and MITF-S409A in the presence of the K316R mutation. The pS73-MITF-WT-K316R and pS73-MITF-sp-K316R proteins were significantly more stable than pS73-MITF-WT; the stability of the S73-MITF-WT-K316R and S73-MITF-sp-K316R proteins was slightly but not significantly increased (Figures 5F and 5G). Interestingly, the pS73-MITF-sl protein was significantly less stable than the pS73-MITF-sp-326* protein (Figures 3F-H), whereas the stability of MITF-sp-378*-K316R was comparable to pS73-MITF-sl (Figures 5F and 5G). This suggests that in the presence of residues 326-419, the K316R mutation increases the stability of MITF, whereas in its absence, K316R mimics the effects of the MITF-sl mutation and reduces stability. Furthermore, the double mutation K316R-S409A appears to have a specific effect on the stability of the S73-MITF-sp protein. However, the stability of pS73-MITF-sp forms remained unaffected by this mutation.

We also determined the stability of MITF-sp, MITF-sp-326*, MITF-sp-378*, and MITF-sp-S409A constructs carrying the E318K mutation. The stability of these proteins was significantly reduced in the presence of the E318K mutation (Figures 5H-I). The MITF-sp-378*-E318K protein showed similar stability as MITF-sl (Figures 5H-I), whereas MITF-sp-326*-E318K and MITF-sp-326* proteins were equally stable and both more stable than MITF-sl, regardless of phosphorylation at S73 (Figures 5H-I). The E318K-S409A mutation resulted in reduced stability of both pS73- and S73-MITF-sp. Taken together, our findings suggest that the E318K mutation reduces the stability of MITF. Interestingly, the K316R and E318K single mutations have different effects on stability even though both eliminate SUMOylation at K316. We hypothesize that the carboxyl domain (aa 378-419) of MITF interacts with the SUMO-site at K316, determining the stability of MITF.

### The carboxyl end IDRs are dynamic and are proximal to each other in the dimer

To probe the interactions between the regions containing the SUMOylation site at K316 and the phosphorylation site at S409, the distance between these domains was measured using smFRET^42^. To enable site-specific labeling, recombinant MITF constructs with only one or two native cysteines were used for inter- and intra-molecular FRET, respectively. When probed intramolecularly, the mean transfer efficiency, denoted as ⟨*E*⟩, between positions C306-C419 showed a slightly expanded chain with ⟨*E*⟩=0.30 (Figure 6A). However, when probed intermolecularly, C306-C419 showed a much higher mean transfer efficiency of ⟨*E*⟩=0.66 (Figure 6B), indicating that the S409 phosphorylation site of one chain is closer to the K316 SUMOylation region of its partner molecule than it is to its own K316 site. All intermolecular distances between different positions in the C-end region were consistently in proximity with mean transfer efficiencies of 0.48-0.85 (Figure 6C). These results suggest that crosstalk between the K316 SUMOylation and S409 phosphorylation region may be between the different monomers of MITF rather than within each monomer. Analysis of the donor fluorescence lifetimes shows that distances between and within the C-terminal IDRs are dynamic on the μs timescale, suggesting the C-terminal IDRs probably do not directly interact with each other to form stable tertiary structures (Figure 6D).

**Figure 6:**
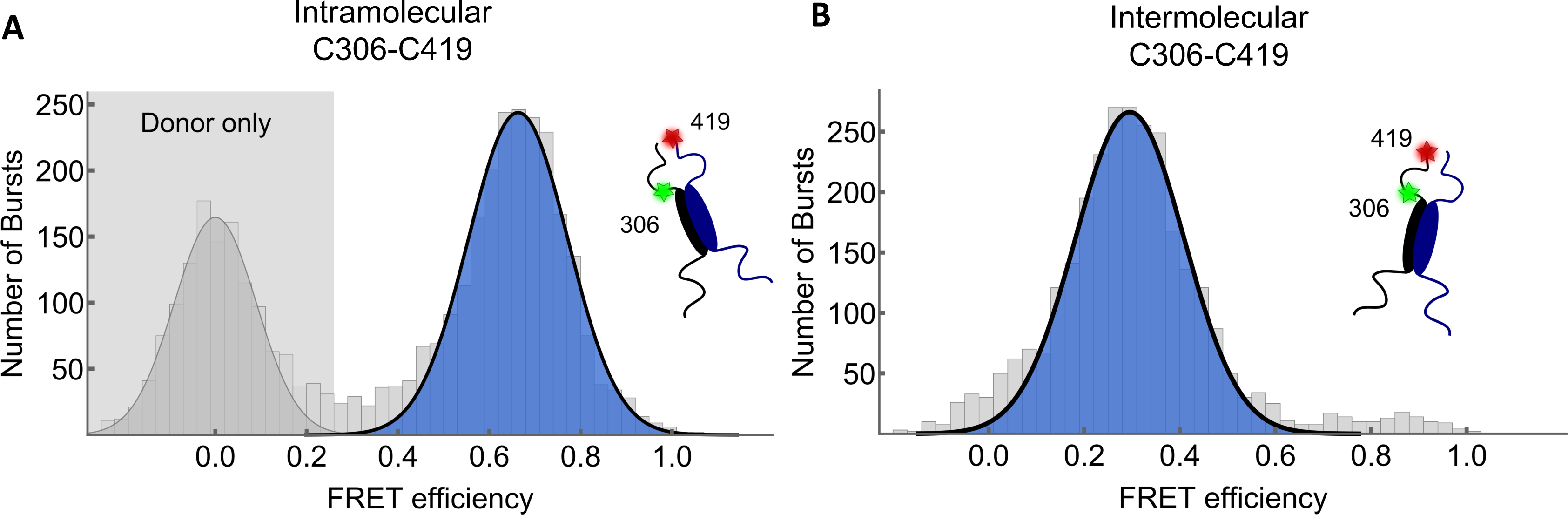
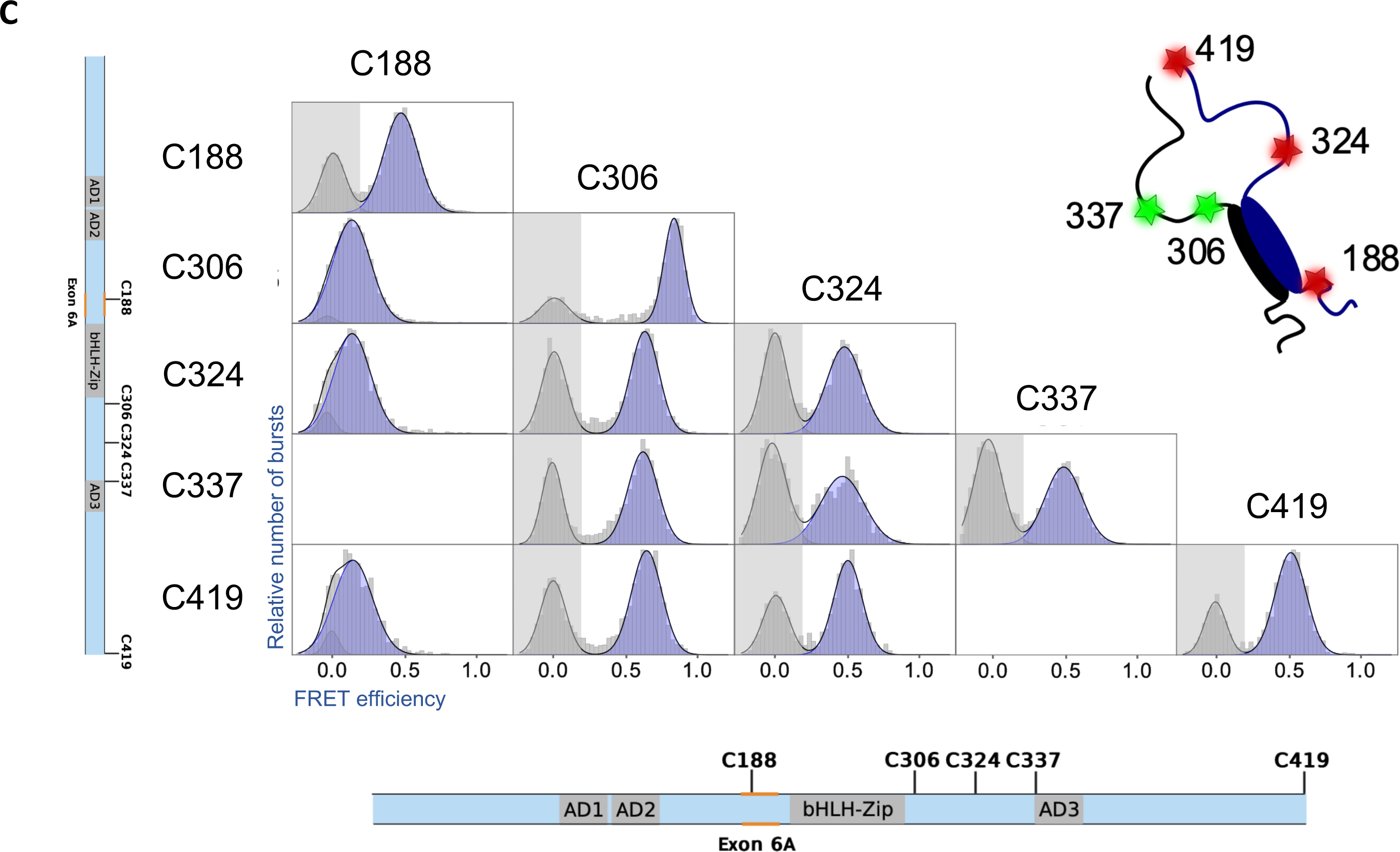
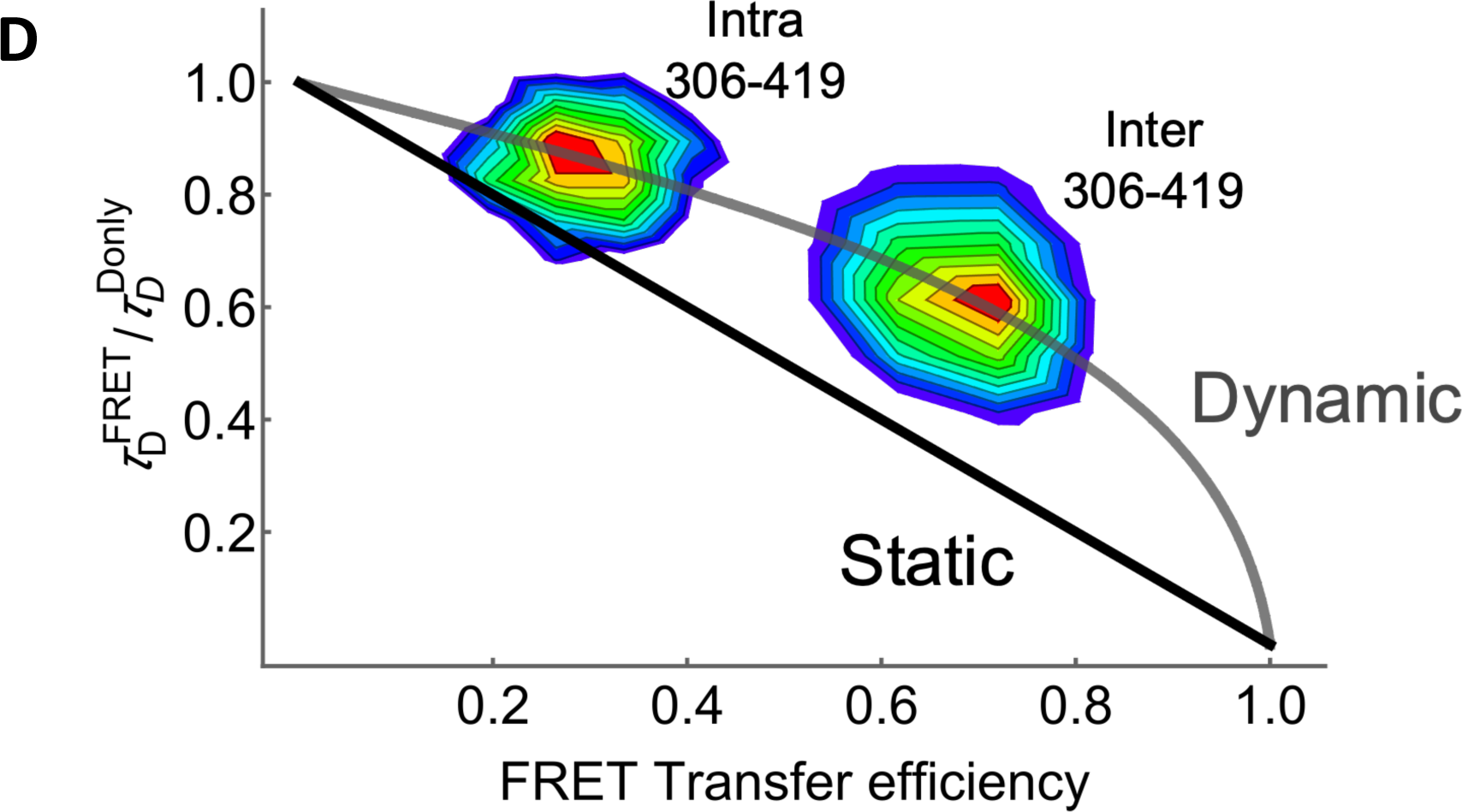
Inter- and intramolecular FRET indicates MITF C-end IDRs are proximal. **(A)** and **(B)** Single-molecule Förster resonance energy transfer histograms of dimeric MITF fluorescently labeled at residues C306 and C419. Intermolecular FRET (A), fitted Gaussian population mean, *E*=0.66, and intramolecular FRET (B), fitted Gaussian population mean, *E*=0.3. **(C)** Single-molecule Förster resonance energy transfer histograms of MITF single-labeled pairs, labeled with acceptor or donor as indicated. FRET population fitted with a Gaussian distribution in blue with donor-only events in grey. **(D)** Fluorescence lifetime analysis of inter- and intramolecular distances of the C-terminal IDR of MITF. The 2-D plot shows the lifetime of the donor in the presence of the acceptor 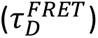 relative to the donor fluorescence in its absence 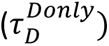, plotted against the FRET transfer efficiency for each burst. The solid black line shows the expected relationship for a static distance, while the grey line shows the expected relationship for a dynamic chain with a Gaussian distribution of distances.

## Discussion

Suppressor mutation screening is an important approach to providing valuable information about gene function, molecular pathways, and protein-protein interactions^43, 44^. Suppressor screens are commonly performed in yeast, Drosophila, and C. elegans but rarely in mice or other mammals. Here, we generated a novel intragenic suppressor mutation at the *Mitf* locus in the mouse and showed that it is a re-mutation at the *Mitf* locus, which results in a truncation of the already mutated MITF-sp protein.

In the homozygous condition, the *Mitf^mi-sl^*mutation leads to brownish coat color compared to the normal black coat of *Mitf^mi-sp^*homozygotes. However, in compound heterozygous conditions with other *Mitf* mutations, including severe dominant-negative or loss-of-function mutations, the *Mitf^mi-sl^* mutation restores the coat color phenotype compared to combinations of the same alleles with the original *Mitf^mi-sp^*mutation. At the molecular level, we show that this suppressor mutation increases the nuclear localization of the MITF-sl protein and reduces its stability. The “brownish” phenotype of *Mitf^mi-sl^* homozygotes is likely to be due to the reduced stability of the MITF-sl protein and the consequent reduction in expression of some pigmentation genes, including *Pmel*, *Tyrp1*, and *Mlana* (Figure S7)^45^. Interestingly, mutations in *Tyrp1* lead to mice with brown coat color. The total concentration of active MITF-sl protein in the nucleus at any given time will depend on the relationship between effects on nuclear import on the one hand and stability on the other hand. Expression of the MITF-partner proteins TFEB and TFE3 is limited in melanocytes, so they are likely to have negligible effects on MITF activity in the homozygous situation. Importantly, when the *Mitf^mi-sl^* mutation is combined with any of the various other *Mitf* mutations (Figures 1 and S1), its ability to dimerize and translocate its partner proteins (MITF-WT or mutant MITF) into the nucleus help to explain the suppressor effects of the *Mitf^mi-sl^* mutation. When in the nucleus, dimers between MITF-sl and any of the defective DNA-binding proteins MITF-Wh, MITF-mi, and MITF-ew slow down MITF-sl degradation but these dimers cannot bind DNA or activate gene expression^4^. Eventually, however, MITF-sl monomers will be released from their non-DNA-binding dimeric partner, thus leading to the formation of MITF-sl homodimers, which can bind DNA and regulate the expression of target genes. Here, the combined effects of nuclear import, rate of nuclear degradation, DNA binding, and dimerization properties are likely to determine the final outcome; the steady-state levels of nuclear MITF-sl are likely to be determined by the rate of heterodimer dissociation and rate of degradation. The near-normal coat color phenotype of *Mitf^mi-sl^/Mitf^mi^* compound heterozygotes suggests that together these effects result in almost full MITF activity during critical stages of melanocyte development and function. This is a novel mechanism of genetic suppression and may partly explain the normal phenotypes observed in humans carrying deleterious mutations on both alleles of genes^46^.

In addition to providing an explanation for the phenotypic outcome of the suppressor mutation, the mutation provides novel insights into how both stability and nuclear export of the MITF protein are regulated. Nuclear localization of MITF has been shown to involve a balance between import and export that depends on a number of domains, including a nuclear localization signal in the DNA-binding domain of MITF and an export signal that depends on the S69 and S73 phosphorylation sites ^10, 34^ (Figure 1A). In wild-type cells, MITF is approximately equally distributed between the nucleus and cytoplasm as determined by western blotting, although, due to differences in nuclear and cytoplasmic volumes, it is more concentrated in the nucleus than in the cytoplasm, as evidenced by immunocytochemistry^34^. Our observations show that the C-end of MITF has major effects on nuclear localization and that residues 316-326, 350-366, and 374-419 are major factors in mediating the nuclear export of MITF. Interestingly, simultaneously mutating the SUMO-site at K316 and the phosphorylation site at S409 increased the nuclear localization of MITF compared to either single mutant alone, suggesting that these two post-translational modifications are necessary for nuclear export. Importantly, the effects of the nuclear export signal mediated by the S73 and S69 phosphorylation^10^ are less efficient when missing the C-end.

Our results show that MITF is mainly degraded through a nuclear ubiquitin-proteasomal degradation pathway. Again, the effects on stability are mainly mediated by the domains encoded by residues 316-326 and 378-419 where K316 and S409 play an important role. However, since all our deletion constructs showed some effect, most regions within the C-domain seem to affect protein stability. This suggests that the entire domain may be important, as is often observed for IDRs. Since the MITF-sl protein is quickly degraded in the nucleus, it is likely that truncation at the C-end activates a degradation signal. A degron motif was recently discovered in the amino end of the A isoform of MITF^47^, but as this is not present in the melanocyte-specific M-isoform studied here, the degradation signal must be located elsewhere in the protein. The fact that the effects on nuclear localization and stability are primarily encoded by the same domains suggests that these events may be related. The effects on nuclear localization are likely dominant since the protein will be degraded by the nuclear proteasome machinery if located in the nucleus. As another layer of regulation, when in the nucleus, MITF stability will also further be regulated by other factors, including SUMOylation at K316 and phosphorylation at S409; DNA binding may also be important, potentially by mediating structural changes. However, how DNA binding contributes to MITF-sl stability is not clear. Importantly our work suggests that the interaction between the SUMOylation site at K316 and the phosphorylation site at S409 is important for regulating MITF localization and stability. Our smFRET results show that these two regions are near each other in space, suggesting that direct interactions between the different protomers are involved.

In contrast to previous literature^48^, our work shows that the S73 form of MITF-WT is much less stable than the pS73 form. In our model, there is an almost 3-fold difference between the two forms. Interestingly, the *Mitf^mi-sl^* mutation reduced the stability of both the pS73 and S73 forms of MITF about 3-fold in each case (Figures 2B and 2C), suggesting that the effects of the C-end on stability are independent of the effects of pS73. It is possible that the difference between the pS73 and S73 forms is due to continuous phosphorylation of the S73-form, possibly mediated by doxycycline treatment, thus affecting the ratio between the two forms of the protein and leading to nuclear export. The observation that pS73-MITF is exported from the nucleus^10^ suggests that the kinetics of S73 phosphorylation and dephosphorylation may determine subcellular location and thus mediate protein stability. Currently, there is limited information on the kinetics or pathways involved.

Independent reports have shown that the E318K variant in human MITF predisposes to melanoma^17, 49^. This variant alters an essential residue in the SUMOylation motif ΨKXD/E which includes K316, the actual SUMOylation site. We show that the E318K mutant protein, which cannot be SUMOylated at this site, exhibits normal nuclear localization. However, when the S409A mutation is also present, the protein is more nuclear, regardless of S73 phosphorylation status. The E318K mutation resulted in reduced MITF stability both in the presence and absence of the S409A mutation. S409 has been suggested to be phosphorylated by the MAP-kinase p90Rsk^48^ or by AKT^50^. The gain-of-function BRAF^V600E^ mutation and loss-of-function PTEN mutations might accelerate the p90Rsk or AKT kinase activity, respectively, and promote S409 phosphorylation. These effects might promote cytoplasmic retention of MITF-E318K, which subsequently would increase the stability of MITF and maintain the level of MITF protein at steady-state levels. Thus, depending on environmental signals (e.g., sun exposure), the medium-risk allele E318K^16, 17^ may mediate disease predisposition.

Based on our data, we propose a model where the two regions of the C-end of MITF, the SUMOylation site at K316 and the phosphorylation site at S409, are impacted by SUMOylation and phosphorylation, leading to effects on nuclear localization and stability. In the absence of SUMOylation at K316 and phosphorylation at S409, these residues are close in space and may collapse around the zipper domain, thus hiding nuclear export signals while at the same time exposing degradation signals.

However, SUMOylation and phosphorylation may change the conformation leading to an extended version of the C-end, thus exposing nuclear export and hiding degradation signals. Although the signals which mediate SUMOylation of K316 and phosphorylation of Ser409 are presently not clear, it has been reported that the unphosphorylated S409 MITF is required to maintain the association of MITF and PIAS3, which enables SUMOylation at K316^13, 51^. This may represent a feedback loop to limit the activity of MITF at any given time based on environmental signals. We, therefore, conclude that generating suppressor mutations in the mouse is an exciting and feasible option for studying gene function and may reveal unexpected aspects of protein function and regulation, leading to novel insights into protein activities in the living organism.

## Materials and Methods

### Mouse strains used, mutagenesis and genotyping

The following *Mitf* mutants were used in this study: C57BL/6J-*Mitf^mi-sp^*, C57BL/6J-*Mitf^mi-eyeless^ ^white^* (*Mitf^mi-ew^*), NAW-*Mitf^mi-ew^*, C57BL/6J-*Mitf^Mi-Wh^*, C57BL/6J-*Mitf^mi-red-eyed^ ^white^* (*Mitf^mi-rw^*), C57BL/6J-*Mitf^microphthalmia^* (*Mitf^mi^*), 82UT-*Mitf^mi-Oak^ ^ridge^*(*Mitf^Mi-Or^*), C57BL/6J-*Mitf^Mi-or^*, 82UT-*Mitf^mi-brownish^*(*Mitf^mi-b^*) and a [C3H/C57BL/6J]-*Mitf^mi-vga-9^*(Table 1). The *Mitf^mi-ew^* mutation arose on the NAW background^52^, and the *Mitf^Mi-Or^* mutation on the 82UT background^53^ both were subsequently backcrossed for 10 generations to C57BL/6J. To screen for Mitf suppressor mutations, homozygous B6-*Mitf^mi-sp^* males were treated four times at one-week intervals with 100 mg/kg ENU. After a 6–8-week recovery period, the males were mated to NAW-*Mitf^mi-ew^*/*Mitf^mi-ew^*females. The resulting offspring, which all should show an identical phenotype, were screened for abnormally pigmented deviates. DNA-HPLC was used to confirm the presence of the *Mitf^mi-sp^* mutation.

### Cell culture, reagents, and antibodies

Melanocyte cultures were obtained from trypsin-digested P0-P3 skin using microscopic selection of pigmented cells. Neural crest cultures were prepared following the previously described protocol ^54^. The number of pigmented melanocytes was counted for each mutant in four independent cultures. MITF-positive cells were detected by immunofluorescence using the C5 anti-MITF antibody and anti-mouse FITC secondary antibody.The cell lines HEK293, A375P (CRL-3224), and SkMel28 (HTB-72) were purchased from ATCC. 501Mel melanoma cells were obtained from the lab of Dr. Ruth Halaban (Yale University). The cells were maintained in RPMI 1640 medium (Gibco, #5240025) supplemented with 10% FBS (Gibco #10270–106) at 5% CO_2_ and 37°C in a humidified incubator. Cycloheximide (CHX – 50 mg/ml) (Sigma, #66819), 20 mg/ml MG132 (Sigma, #474790), 10 mg/ml Doxycycline (Dox – Sigma, #324285), 10 mg/ml TPA (Merck, #P1585), 1mg/ml Baf-A1 (Merk, #88899-55-2), 5 mM Leptomycin B (Merk, #L2913), 5 mM PLX4032 (Selleckchem, #S1267) stock solutions were prepared in DMSO. The primary antibodies used for all western blot (WB) experiments and their dilutions were as follows: Anti-FLAG (Sigma, #F3165) at 1:5000 dilution; Anti-β-Actin (Cell Signaling, #4970) at 1:1000 dilution; Anti-γH2AX (Abcam, #ab2251) at 1:2000 dilution, Anti-GAPDH (Cell Signaling, #5174), at 1:1000 dilution; Anti-H3K27me3 (Cell Signaling, #9733) at 1:1000 dilution; Anti-GFP (Abcam, #ab290) at 1:2500 dilution.

### Generation of plasmid constructs for stable doxycycline-inducible overexpression

Fusions of wild-type and mutant mouse MITF-M cDNA with the 3XFLAG-HA tag at the C- or N-terminus or fusion with the GFP-tag at the C terminus were generated in the piggy-bac vector pPB-hCMV1. The cDNAs were subcloned downstream of a tetracycline response element (TRE) using the Gibson Assembly Cloning kit (New England Biolabs, # E5510S). Mutations were introduced by in vitro mutagenesis using Q5 Site-directed Mutagenesis Kit (New England Biolabs, #E0554S) according to the manufacturer’s instructions.

### Generation of stable doxycycline-inducible MITF

Inducible A375P, SKmel28, and 501Mel cells were generated as described before ^55^. Briefly, the wild-type and mutant mouse MITF fusion constructs with the 3XFLAG-HA at the C- or N-terminus or fusion with GFP at C-terminus or a pPB-hCMV1-EV-3XFLAG-HA empty vector was transfected into 70–80% confluent A735P, 501Mel or SKMel28 cells using Fugene HD reagent (Promega, #E2311) together with the py-CAG-pBase and pPB-CAG-rtTA-IRES-Neo plasmids at a 10:10:1 ratio. After 48 hours of transfection, the transfected cell lines were selected with 0.5 mg/ml of G418 (Gibco, #10131-035) for two weeks. A ‘mock plate’ of no transfected cells was also included in each case. To equalize the expression of MITF proteins, the dox-inducible A375P, 501Mel, and SKmel28 melanoma cell lines were treated with varying concentrations of doxycycline, and the doxycycline concentrations leading to similar MITF protein levels were used for future experiments.

### Incucyte live cell imaging

Dox-inducible A375P melanoma cells overexpressing MITF-WT, MITF^mi-sp^, and MITF^mi-sl^ were seeded at 2000 cells per well in triplicate onto a 96-well cell culture plate (Falcon 96-well Clear Flat Bottom TC-treated Culture Microplate, CORNING, Cat#353072). Images were recorded with Incucyte S3 Live-Cell Analysis System (Sartorius, Essen BioScience) every 2 hours for four days. Collected images were then analyzed using the Incucyte software by measuring the percentage of cell confluency.

### qRT-PCR and sequencing

Total RNA was isolated from hearts of wild type and mutant mice using the RNAwiz kit (ThermoFisher, #Am1925). The RNA was reverse transcribed by SuperScript reverse transcriptase (Invitrogen), and the resulting cDNA phenol/chloroform was extracted. Alternatively, RNA was isolated using the Macherey Nagel RNAII kit. The entire *Mitf* cDNA was amplified by PCR using overlapping primers. The resulting PCR products were sequenced directly using the Big Dye Terminator Cycle Sequencing kit (ABI) and the ABI 377 sequencer.

The day before inducing MITF expression by doxycycline, cells were seeded on 12-well plates at a density of 12×10^4^ cells per well. MITF expression was induced for 6, 12, 24, and 36 hours and harvested for RNA isolation using TRIzol reagent (ThermoFisher, #15596–026) High-Capacity cDNA Reverse Transcription Kit (Applied Biosystems, #4368814) was used for cDNA synthesis according to the manufacturer’s instructions. The SensiFAST SYBR Lo-ROX Kit (Bioline, #BIO-94020) was utilized for the qRT-PCR. qRT-PCR reactions were performed using 0.4 ng/µl cDNA in triplicates. The relative foldchange in gene expression was calculated using the D-ΔΔCt method ^56^. The geometrical mean of β-actin and hARP expression was used to normalize the target gene expression.

### Subcellular fractionations

The day before inducing MITF expression by doxycycline, cells were seeded on 6-well plates at a density of 3.5×10^5^ cells per well for 24 hours, after which the cells were either directly harvested or treated with TPA at 200nM for 1 hour, 4 hours, or treated with TPA at 200nM for 1 hour and MG132 40 µg/ml for the next 3 hours in the presence of TPA before harvesting by trypsinization. Cells were washed with PBS before washing twice with swelling buffer (10 mM HEPES, pH 7.9, 1.5 mM MgCl_2_, 10 mM KCl, 0.5 mM DTT, and freshly added protease and phosphatase inhibitors). The cells were then lysed by incubation at 4°C for 15min in cell lysis buffer (10 mM HEPES, pH 7.9, 1.5 mM MgCl_2_, 10 mM KCl, 0.5 mM DTT, 0.1% NP40). Approximately 30% of the sample was collected and set aside as whole cell lysate. The remaining cell lysate was spun down at 3000 rpm for 5min at 4°C, and the supernatant was collected as the cytoplasmic fraction. At the same time, the pellet, representing the nuclear fraction, was washed with cold PBS before resuspension in RIPA buffer (20 mM Tris-HCl, pH7.4, 50 mM NaCl, 2 mM MgCl_2_, 1% (v/v) NP40, 0.5% (m/v) sodium deoxycholate, and 0.1% (m/v) sodium dodecyl sulfate, and freshly added protease and phosphatase inhibitors) for further experiments including Western blotting and immunoprecipitation.

### Immunoprecipitation

Cells were seeded on 6-well plates at a density of 3.5×10^5^ cells per well the day before transfection. The following day, FuGENE HD reagent (Promega # E2311) was used to conduct the co-transfection of MITF-WT, MITF^mi-sp^ or MITF^mi-sl^-GFP-tagged constructs together with MITF^mi^, MITF^mi-ew^, MITF^Mi-Wh^, 14-3-3-Epsilon or 14-3-3-zeta Flag-tagged proteins. After 24 hours, the cells were washed twice with ice-cold PBS and lysed by adding 200 µl of RIPA buffer with freshly added protease and phosphatase inhibitors. The cell lysate was then ready for immunoprecipitation (IP); 30% of the sample was collected as an input fraction.

For each IP sample, 20 µl of Dynabeads Protein G magnetic beads (Invitrogen, # 10004D) were washed twice with 1ml PBS using a magnetic stand before resuspending in 300 µl of PBS containing 0.01% Tween 20. The magnetic beads were then conjugated with anti-FLAG antibodies by adding 1 µg anti-FLAG antibody (Sigma, #F3165), followed by a 30-minute incubation at RT with rotation. The magnetic beads were washed twice with PBS containing 0.01% Tween 20 to eliminate non-conjugated antibodies and then resuspended with 20 µl of PBS containing 0.01% Tween 20. The IP samples were incubated with the coated beads overnight at 4°C with rotation. Samples were then placed on the magnetic stand, and supernatants were removed and saved as an unbound fraction (UnB) in each case. The beads were washed twice with 1ml PBS containing 0.01% Tween 20. The protein was eluted from the beads by incubating with 150 ng/µl 3X Flag peptide in PBS containing 0.01% Tween 20 for 30 minutes at 4°C with rotation. The samples were placed on the magnetic stand, and supernatants were saved as an immunoprecipitation fraction (IP). The collected fractions were then subjected to Western blot analysis.

### EMSA DNA binding studies

Electrophoretic Mobility Shift Assays were performed using proteins expressed in the TNT T7 Coupled Reticulocyte Lysate System (Promega, WI), according to the manufactureŕs recommendation. DNA binding reactions were performed in 10 mM Hepes (pH 7.9), 50 mM NaCl, 5 mM MgCl_2_, 0.1 mM EDTA, 2 mM dithiothreitol (DTT), 5% ethylene glycol, and 5% glycerol. Ten µL of 2X buffer were combined with 2.5 µL of TNT translated MITF protein, 1.4 ng of labeled probe (5’-AAAGTCAGTCATGTGCTTTTCAGA-3’), 4 µg of bovine serum albumin (BSA) ^57^ and water, adjusting the reaction volume to 20 µL. For supershifts, 0.5 µL of C5-monoclonal MITF antibody (Neomarkers) was added to the reaction. The samples were incubated on ice for 30 minutes to allow binding to proceed. The resulting DNA-protein complexes were resolved on 6% non-denaturing polyacrylamide gels, placed on a storage phosphor screen, and scanned on a Typhoon Phosphor Imager 8610 (Molecular Dynamics) for analysis.

### Protein degradation assay

The dox-inducible A375P cells were treated with doxycycline to express MITF WT and MITF mutant proteins for 24 hours and then treated with 40 µg/ml cycloheximide (Sigma #66819), in presence or absence of 200nM TPA, 2µM PLX, or 5nM LMB for 0, 1, 2, and 3 hours before harvesting. For protein degradation pathway analysis, the dox-inducible A375P cells were treated with doxycycline to express the respective MITF constructs for 24 hours at density of 8×10^4^ cells and then treated with either 40 µg/ml MG132 or 2 µg /ml Bafa1 in the presence or absence of 40 µg/ml CHX for 3 hours before harvesting. FuGENE HD reagent (Promega, #E2311) was used for the co-transfection. After 24 hours, the cells were treated with 55 µg/ml cycloheximide (Sigma, #66819) for 0, 1, 2, and 3 hours. The cells were finally lysed in SDS sample buffer (2% SDS, 5% 2-mercaptoethanol, 10% glycerol, 63 mM Tris-HCl, 0.0025% bromophenol blue, pH 6.8), and the expression of the MITF protein determined by Western blot using FLAG antibodies.

### Western blot analysis

For the Blue native PAGE, The A375P melanoma cells transiently co-expressed the MITF^mi^-Flag protein with either MITF-WT-EGFP, MITF^mi-sp^-EGFP, or MITF^mi-sl^-EGFP proteins or expressed MITF-WT-Flag, MITF^mi-sp^-Flag, MITF^mi-sl-^Flag, MITF^mi^-Flag in A375 cells for 24 hours before harvested by NativePAGE Sample Buffer (Thermo Scientific, #BN2003). We then performed Blue native PAGE (Wittig et al., 2006) according to the manufacturer’s instructions (Thermo Scientific, # BN1001) followed by a second dimension of SDS-PAGE. For cell lysates in SDS sample buffer, the samples were boiled for 5 minutes at 95°C. SDS-PAGE separated proteins. Proteins were then transferred to 0.2 μm PVDF membranes (Thermo Scientific, #88520). The membranes were blocked in T-TBS (20 mM Tris, pH 7.4; 150 mM NaCl; 0.01% Tween 20) containing 5% BSA. The membranes were probed with specific primary antibodies. After washing three times with T-TBS for 10 minutes each, the membrane was then incubated for 1 hour at room temperature with either DyLight 800 anti-mouse (Cell Signaling, #5257) or DyLight 580 anti-rabbit IgG (Cell Signaling, #5366) secondary antibodies (Cell Signaling Technology). The protein bands were detected using Odyssey CLx Imager (LICOR Biosciences) and Image Studio version 2.0. The band intensity was quantified using the open-access ImageJ software (https://imagej.nih.gov/ij/).

### BrdU assay

HEK293 cells were transfected with Mitf^mi-sp^ or Mitf^mi-sl^ cDNA constructs, and 18 hours after transfection, the cells were incubated in 10 µM BrdU (Sigma) for 30 min at 37 °C. The cells were fixed and double-labeled for MITF and BrdU as described ^58^. Three independent experiments were performed, each counted in triplicates.

### Generation of protein expression vectors

A bacterial codon-optimized human MITF-M+ ORF synthesized by GenScript Biotech was cloned using Gibson assembly (New England Biolabs, # E5510S) into a pET24a vector, C-terminally in frame with a 6xHistidine tagged SMT3 (SUMO) fusion protein. To obtain single cysteine mutant versions of MITF for maleimide labeling, two successive rounds of cysteine to serine mutagenesis were performed using the QuikChange Lightning Multi Site-directed Mutagenesis kit (Agilent Technologies #210513/#210515) according to manufacturer’s instructions at 1/4^th^ scale.

### Recombinant protein expression, purification, and labeling

The protein was expressed from Lemo21(DE3) *E.coli* by autoinduction at 37°C for 8-12 hours. Harvested cells were resuspended in 50 mM Tris, 10 mM EDTA, 1 mM PMSF, 1% Triton X-100, pH=8.2, lysed by sonication, and centrifuged at 40000×g for 30 minutes. The insoluble material was washed in 50 mM Tris, 150 mM NaCl, 0.05% Triton X-100, pH=8, and re-centrifuged, followed by resuspension in 8 M Urea, 500 mM NaCl, 20 mM Imidazole, 20 mM Na_2_HPO_4_, 0.1 mM PMSF, 0.1 mM TCEP, 0.01% Tween-20, pH=8.2. The resuspended pellet was centrifuged at 40000×g for 30 minutes and the supernatant was loaded onto a 5 ml HisTrap HP column (Cytiva) connected to an Äkta Pure chromatography system using a flowrate of 5 ml/min. The column was washed with a buffer containing 6 M Urea, 1 M NaCl, 25 mM Imidazole, 20 mM Na_2_HPO_4_, 0.01% Tween-20, pH = 8.2, until 280 nm UV absorption reached baseline. The column was washed with at least 4 CV of running buffer containing 5 M Urea, 500 mM NaCl, 20 mM Imidazole, 20 mM Na_2_HPO_4_, 0.01% Tween-20, pH=8.2, and the protein eluted in 4 CV of 50% running buffer, 50% elution buffer containing 2 M Urea, 500 Mm NaCl, 300 mM Imidazole, 50 mM Na_2_HPO_4_, 0.01% Tween-20, pH=7.8. Fractions containing more than 4 mg/ml of protein were pooled, treated with 10 mM of DTT at RT for 15 minutes, and then diluted with a buffer containing 3.5 M Urea, 500 mM NaCl, 50 mM Tris, 0.01% Tween-20, pH=8, to 4 mg/ml. The diluted mix was then further diluted to a Urea concentration below 1 M by addition of a buffer containing 50 mM Tris, 150 mM NaCl, 0.05% Triton X-100, pH=8. The His6-SUMO tag was enzymatically removed by addition of ULP-1 protease at a substrate-to-enzyme molar ratio of 1:200. The reaction was incubated at RT for 12-36 hours, followed by SDS-PAGE to confirm digestion and purity. The sample was precipitated from with 10% TCA, the precipitate pelleted by centrifugation, washed with EtOH, and resuspended in a minimal volume of 8 M Urea, 50 mM K_2_HPO_4_/KH_2_PO_4_, 10 mM DTT, pH=8.2, incubated for 1 hour and centrifuged at 17000×g for 5 minutes followed by application to a HiTrap Desalting column equilibrated in the same buffer containing no DTT.

For labeling, one mg of eluted protein at a concentration of 50-150 µM was incubated with a 2× molar excess of CF660R (Biotium) or Cy3B (Cytiva) overnight at RT. Protein was precipitated by adding EtOH and resuspended in 7 M GdmHCl, 100 mM DTT, 50 mM K_2_HPO_4_/KH_2_PO_4_, pH=6.7. Labeling steps were omitted for the MITF-WT, MITF-sp, and MITF-sl constructs used for DNA binding experiments. Samples were purified using RP-HPLC on a Zorbax 300SB-C3 column, and fractions corresponding to the target protein were lyophilized. Fractions were resuspended in 7 M GdmHCl, 50 mM K_2_HPO_4_/KH_2_PO_4_, pH=6.7, and analyzed by UV-Vis to determine protein concentration and degree of labeling before long-term storage at −80 °C.

### DNA probe labeling and purification

Functionalized DNA oligonucleotides were synthesized by IDT Inc, positive strand /5AmMC6/GAGATCATGTGTTGA and negative strand /5AmMC6/TCAACACATGATCTC. Both oligonucleotides were resuspended in 100 mM Sodium Bicarbonate to a concentration of 1 mM. The forward strand was incubated with 10 fold molar excess of Cy3B NHS ester overnight at RT, and the negative strand was incubated in the same way with CF660R NHS ester. Reactions were precipitated by adding 0.1 volumes of 4 M sodium acetate and 3 volumes 75% EtOH, incubated at −20°C for an hour, followed by centrifugation at 25000 ×g for 30 minutes. The pellet was resuspended in a minimal volume of DNA RP-HPLC solution A (100 mM triethylammonium acetate, pH=8, 5% acetonitrile) and purified using RP-HPLC on a ReproSilGold 200å C18 column with a linear gradient of 5% RP-HPLC solution B (acetonitrile) to 55% over 35 minutes at 1 ml/min. Fractions corresponding to labeled oligonucleotides were collected, freeze-dried, and resuspended in DNA annealing buffer (10 mM Tris, 1 mM EDTA, 50 mM NaCl, pH = 8), mixed 1:1 at a concentration of 20 μM and annealed by incubation at 95°C for 5 minutes and passive cooling to RT.

### Single-molecule fluorescence spectroscopy

Single-molecule experiments were performed at 20-23°C using a confocal MicroTime 200 instrument (PicoQuant). The donor dye was excited using a 520 nm diode laser (LDH-D-C-520, PicoQuant) at 20-30 μW of power (measured after the dichroic) using pulsed interleaved excitation at 20 MHz repetition rate ^59^. The acceptor dye was excited using a 640 nm diode laser (LDHD-C-640, PicoQuant) at 15-20 μW of power. Excitation and emission light were collected through a 60× water-immersion objective (UPLSAPO60XW, Olympus) and focused onto a 100 or 50 μm pinhole, separated by polarization, donor, and acceptor emission wavelengths. Photons were detected using four identical SPADs (SPCM-AQRG-TR, Excelitas Technologies).

For intermolecular FRET measurements, pairs of single-labeled hMITF-M+ (hMITF-WT) were mixed together in 7 M GdmHCl to a total concentration of 10 μM or greater at an acceptor-to-donor ratio of 5:1. The sample was then diluted to 100-400 nM in 10 mM Tris, 0.1 mM EDTA, 165 mM KCl, 0.001% Tween-20, pH=7.4, prior to dilution to 0.5-2 nM for experiment. All experiments using single-labeled pairs were performed using ibidi μ-slide sample chambers at a volume of 40-50 μL in 10 mM Tris, 0.1 mM EDTA, 165 mM KCl, 1 mM MgCl_2_, 143 mM 2-ME, 0.001% Tween-20, pH=7.4. Data were analyzed using Mathematica 12 and the Fretica plugin (https://schuler.bioc.uzh.ch/programs/).

For intramolecular FRET measurements, double-labeled h-MITF-M+ was diluted to a concentration of 100 nM in 10 mM Tris, 0.1 mM EDTA, 165 mM KCl, 1 mM MgCl_2_, 0.001% Tween-20, pH = 7.4, prior to dilution to ∼100 pM for the experiment in the same buffer with 143 mM 2-ME added.

DNA-binding experiments were performed using a labeled M-Box probe, prepared as described above, at a concentration of 0.5 nM in 10 mM Tris, 0.1 mM EDTA, 165 or 300 mM KCl, 1 mM MgCl_2_, 143 mM 2-ME, 0.001% Tween-20, 0.1 mg/ml BSA pH=7.4. For refolding, purified hMITF-WT, hMITF-sp, and hMITF-sl were diluted 1000-100-fold to 500 nM in the same buffer containing 14 mM 2-ME, incubated on ice for at least 30 minutes and added to the M-box probe at least 15 minutes before measurement to ensure equilibrium conditions.

### Statistical analysis

Data were analyzed with GraphPad Prism 9.0 software (San Diego, CA). The results were obtained from at least three biological replicates. An unpaired t-test was conducted to compare the two groups. A significant difference was established with * p< 0.05, *p< 0.05, ** p < 0.01, *** p < 0.001, **** p < 0.00, and ns not significant. All the data were expressed as mean ± SEM.

## Supplement Tables

**Table S1.**
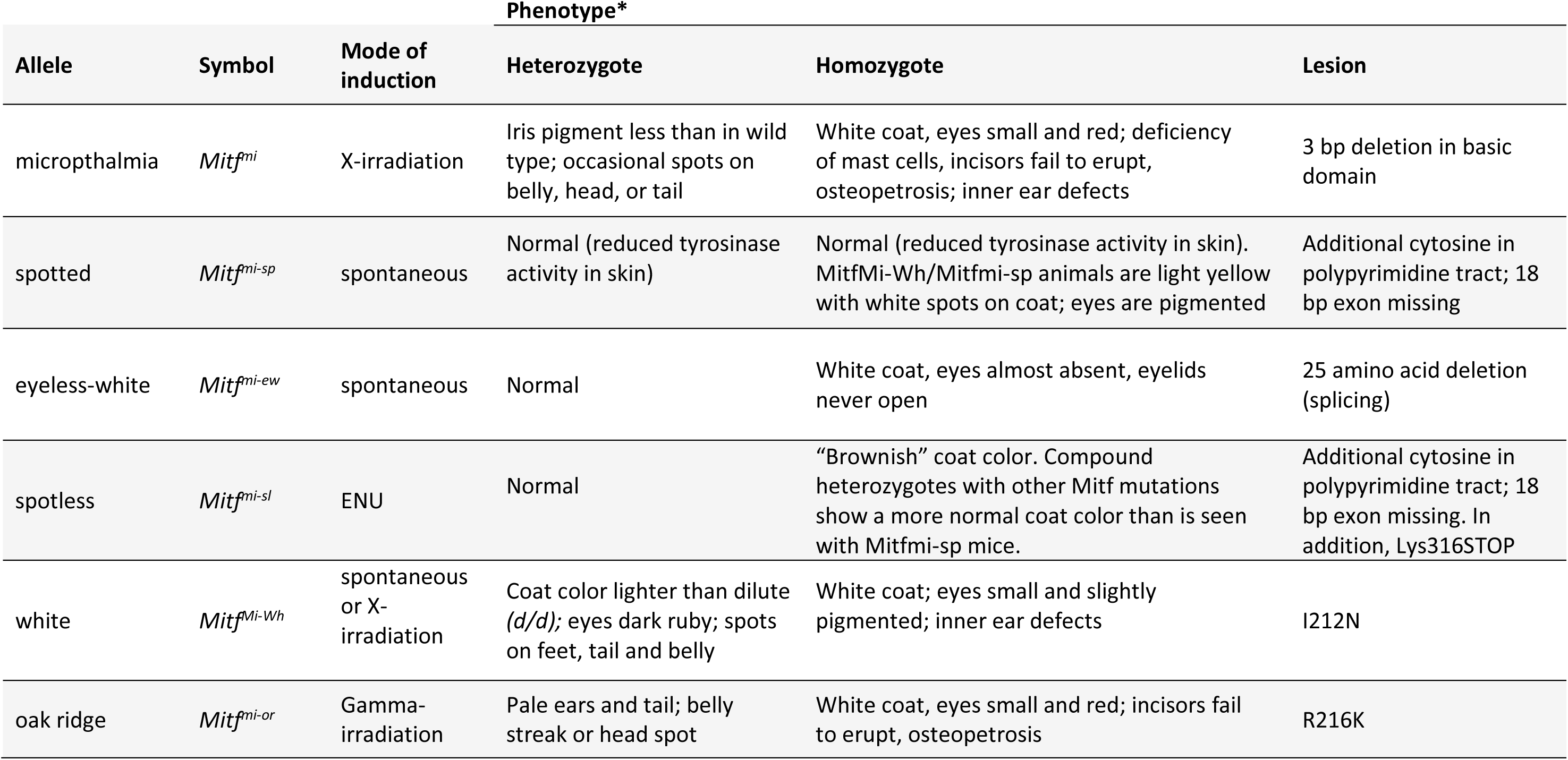
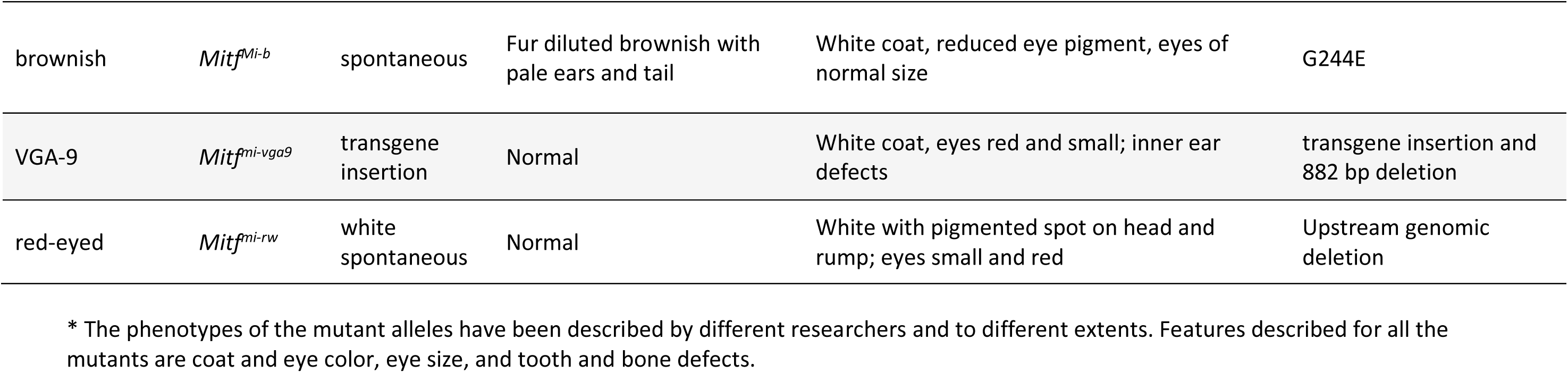
The Mitf mutants used in this study.

**Table S2.** DNA binding affinity of recombinant human MITF-WT, MITF-sp, and MITF-sl to M-box.

| Construct | Anisotropy | FCS | FRET |
| --- | --- | --- | --- |
| **hMITF-WT** | 1.9 ± 0.6 nM | 4.5 ± 1.4 nM | 1.6 ± 1.4 nM |
| **hMITF-sp** | 1 ± 0.4 nM | 5.8 ± 3.5 nM | 1.7 ± 0.9 nM |
| **hMITF-sl** | 0.9 ± 0.4 nM | 3.1 ± 1.6 nM | 1.6 ± 0.5 nM |

## Supplementary Figure Legends

**Figure S1:**
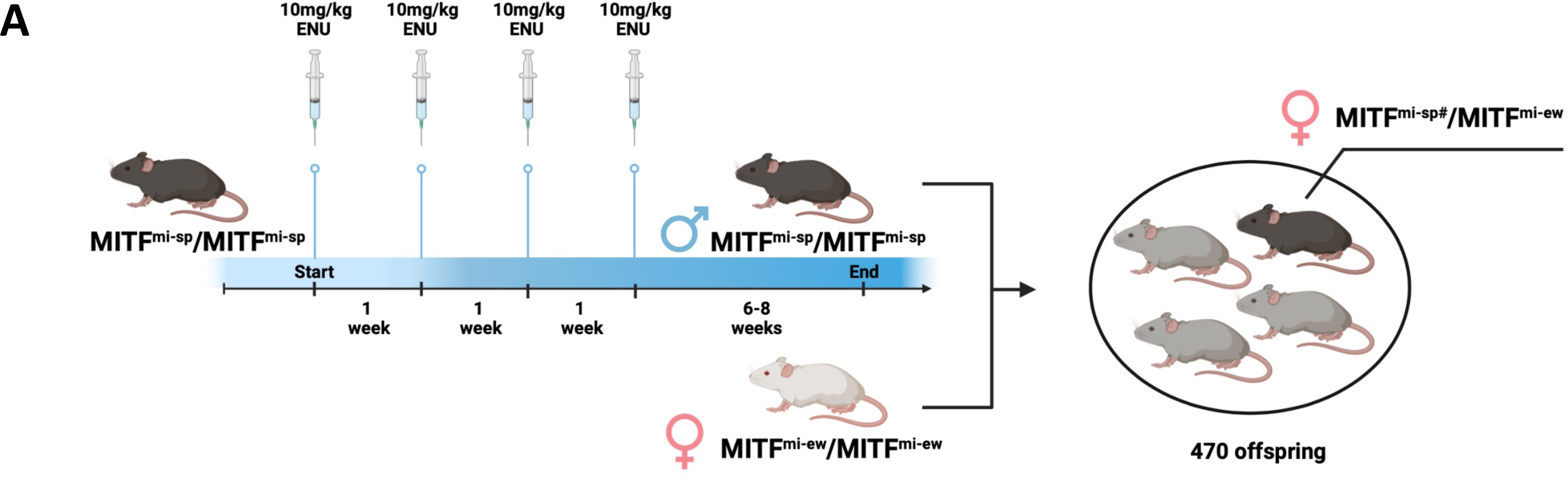
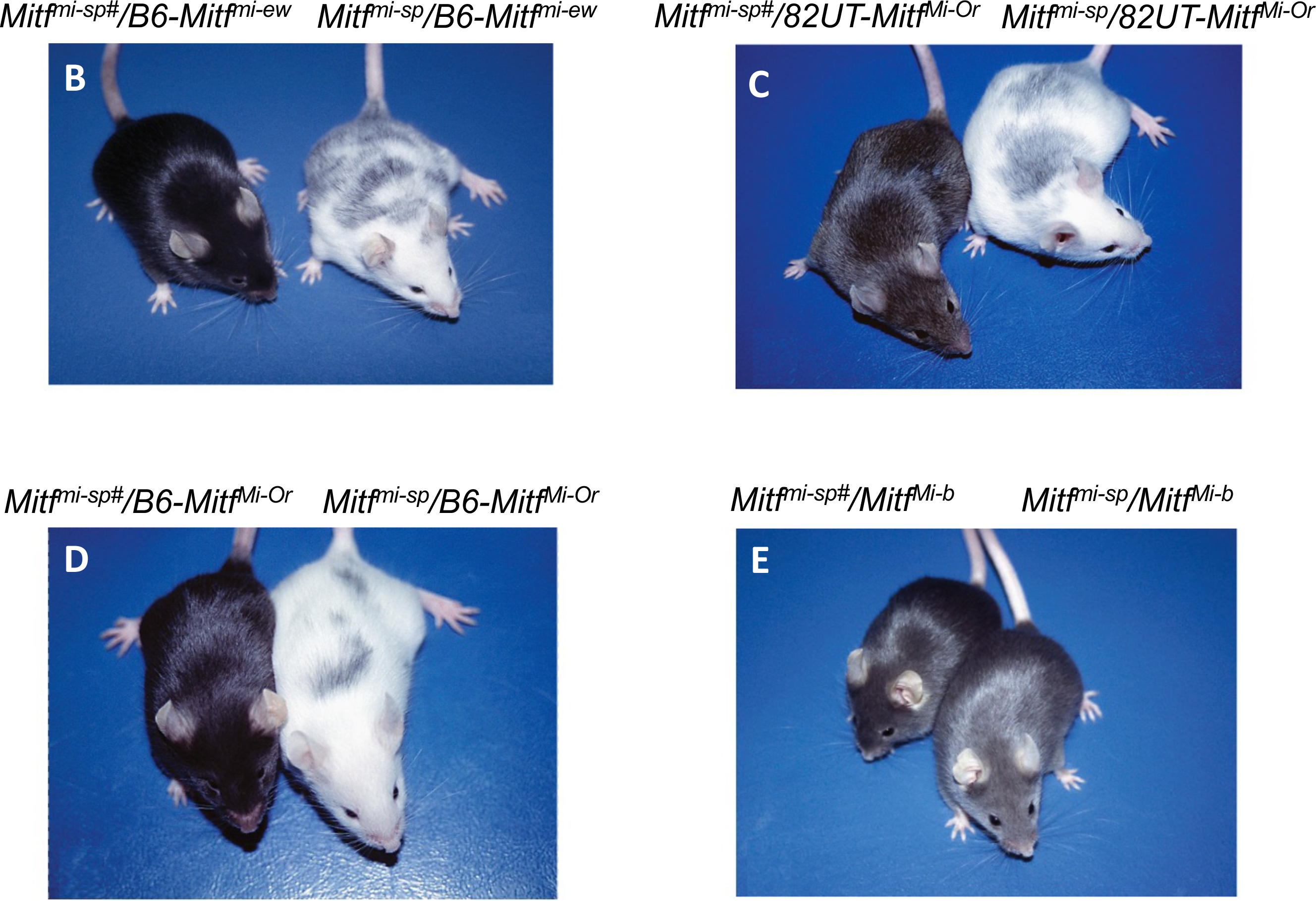
Generation and phenotypic behavior of the induced *Mitf^mi-sp#^* suppressor mutation. (A) Schematic of generation of a *Mitf* suppressor mutation in mouse. (B) B6-*Mitf^mi-ew^*/B6-*Mitf^mi-sp#^* and B6-*Mitf^mi-ew^*/B6-*Mitf^mi-sp^* compound heterozygotes. (C) 82UT-*Mitf^Mi-Or^*/B6-*Mitf^mi-sp#^* and 82UT-*Mitf^Mi-Or^*/B6-*Mitf^mi-sp^* compound heterozygotes. (D) B6-*Mitf^mi-sp#^*/B6-*Mitf^Mi-Or^ and B6-Mitf^mi-sp^*/B6-*Mitf^Mi-Or^* compound heterozygotes. (E) B6-*Mitf^mi-sp#^*/B6-*Mitf^Mi-b^* and B6-*Mitf^mi-sp^*/B6-*Mitf^Mi-b^* animals.

**Figure S2:**
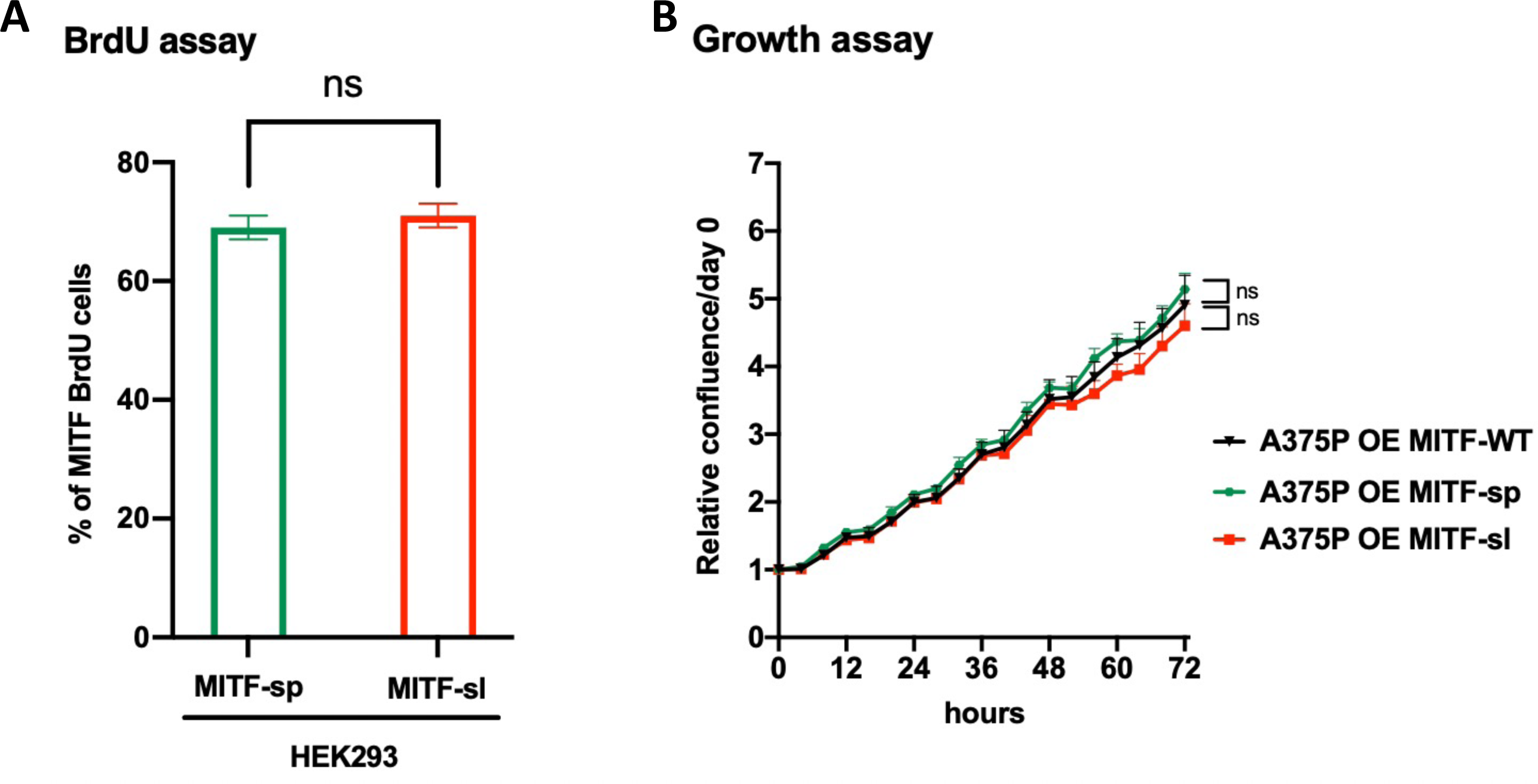
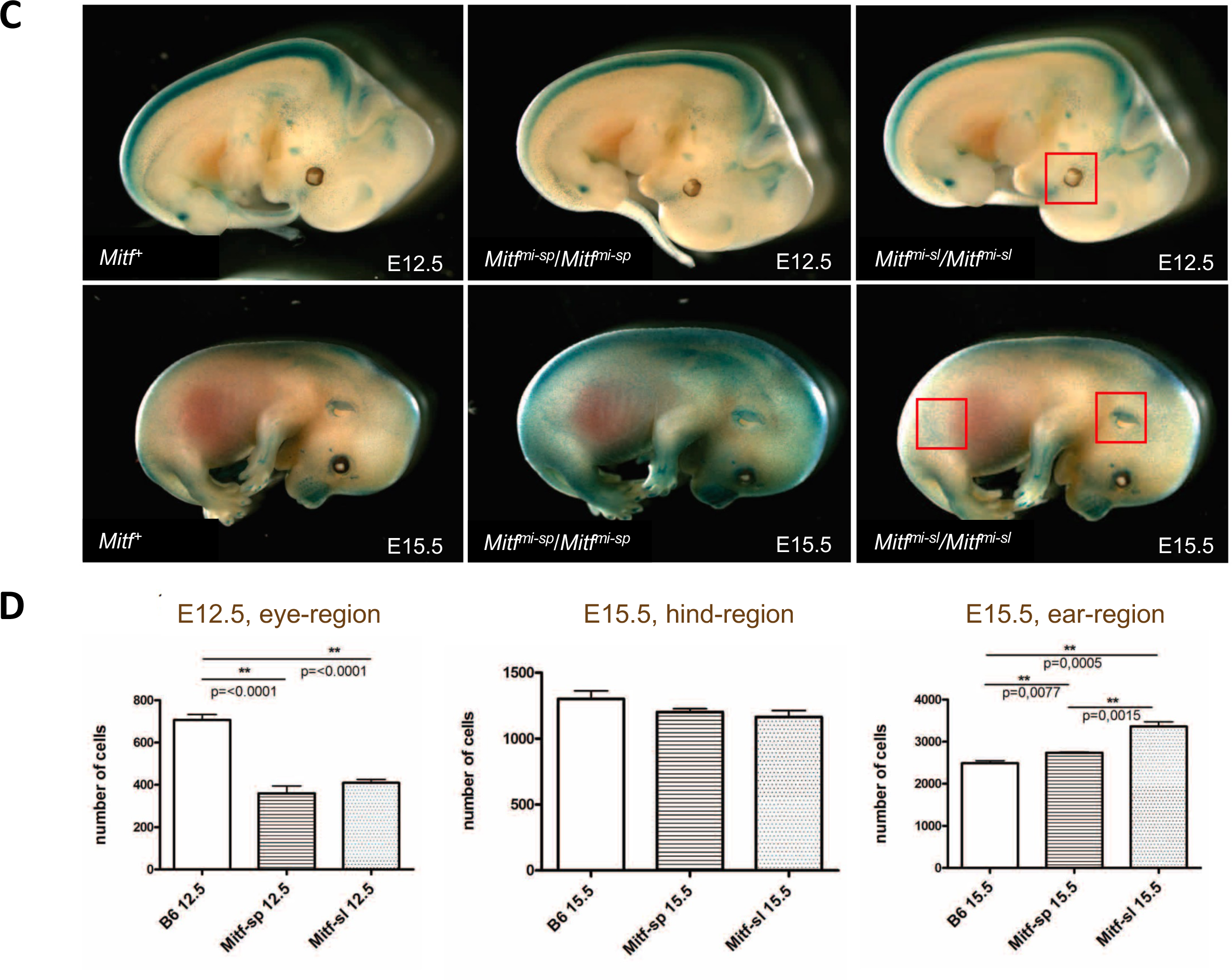
*Mitf^mi-sl^* did not affect cell proliferation or melanocyte numbers in developing embryos. (A) Percentage of BrdU-positive cells was assessed. Error bars represent SEM of three independent experiments. (B) Relative cell confluence compared to day 0 obtained from IncuCyte proliferation assay was plotted for A375P expressing Mitf *WT, Mitf^mi-sp,^ or Mitf^mi-sl^.* Error bars represent SEM of three independent experiments. **(C)** and **(D)** The Dct-LacZ transgene was crossed into *Mitf^mi-sl^* and *Mitf^mi-sp^*mice, and the number of melanocytes, as determined by the presence of X-gal-labeled blue cells, determined in selected regions in 12.5-day-old and 15.5-day-old embryos (indicated by red boxes). Statistically significant differences (Student’s t-test) are indicated by * p< 0.05, *p< 0.05, ** p < 0.01, *** p < 0.001, **** p < 0.00, and ns not significant.

**Figure S3:**
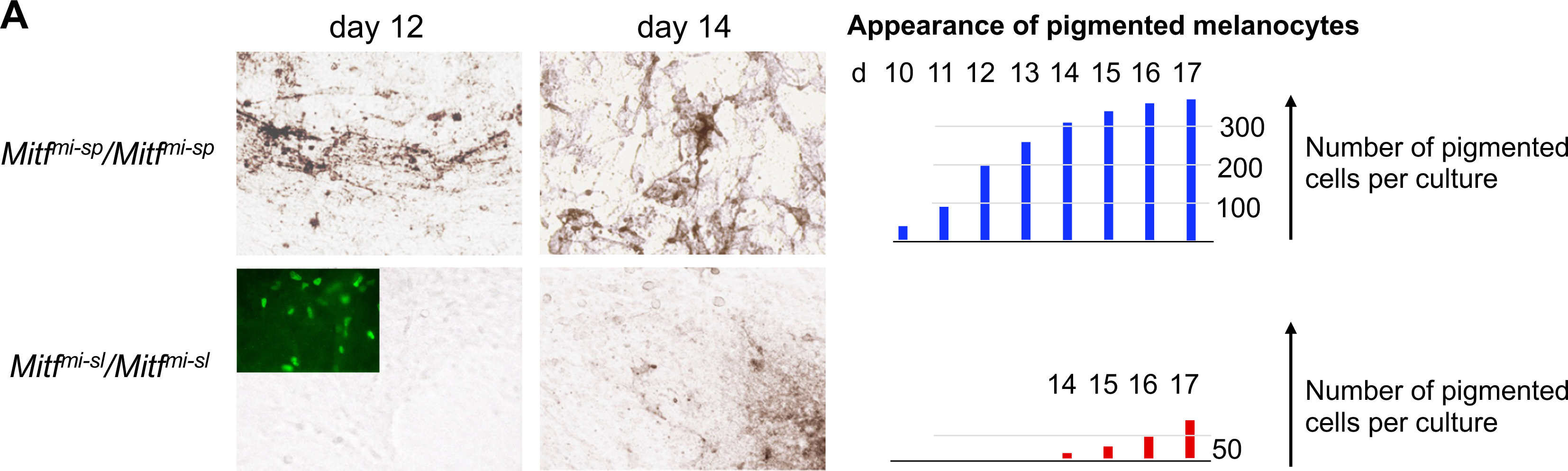
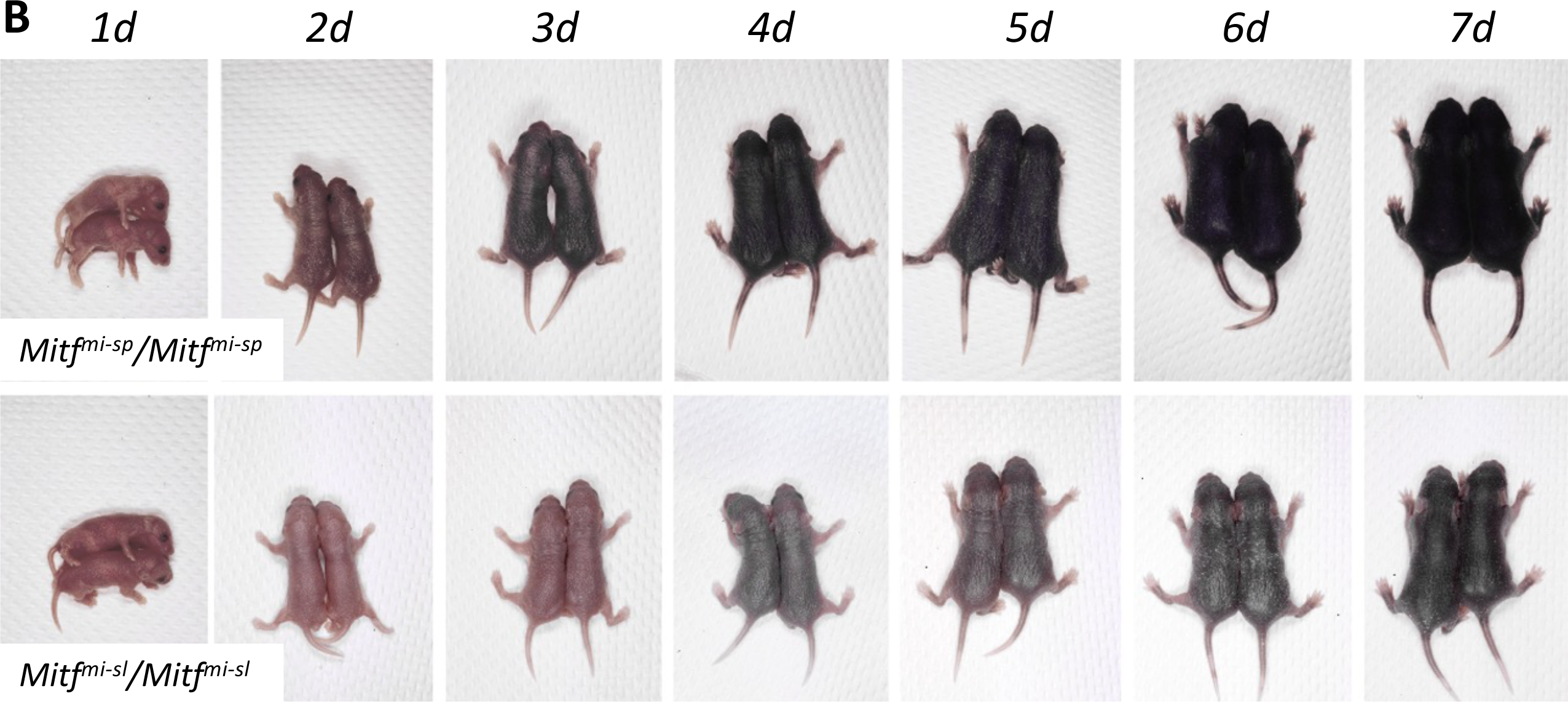
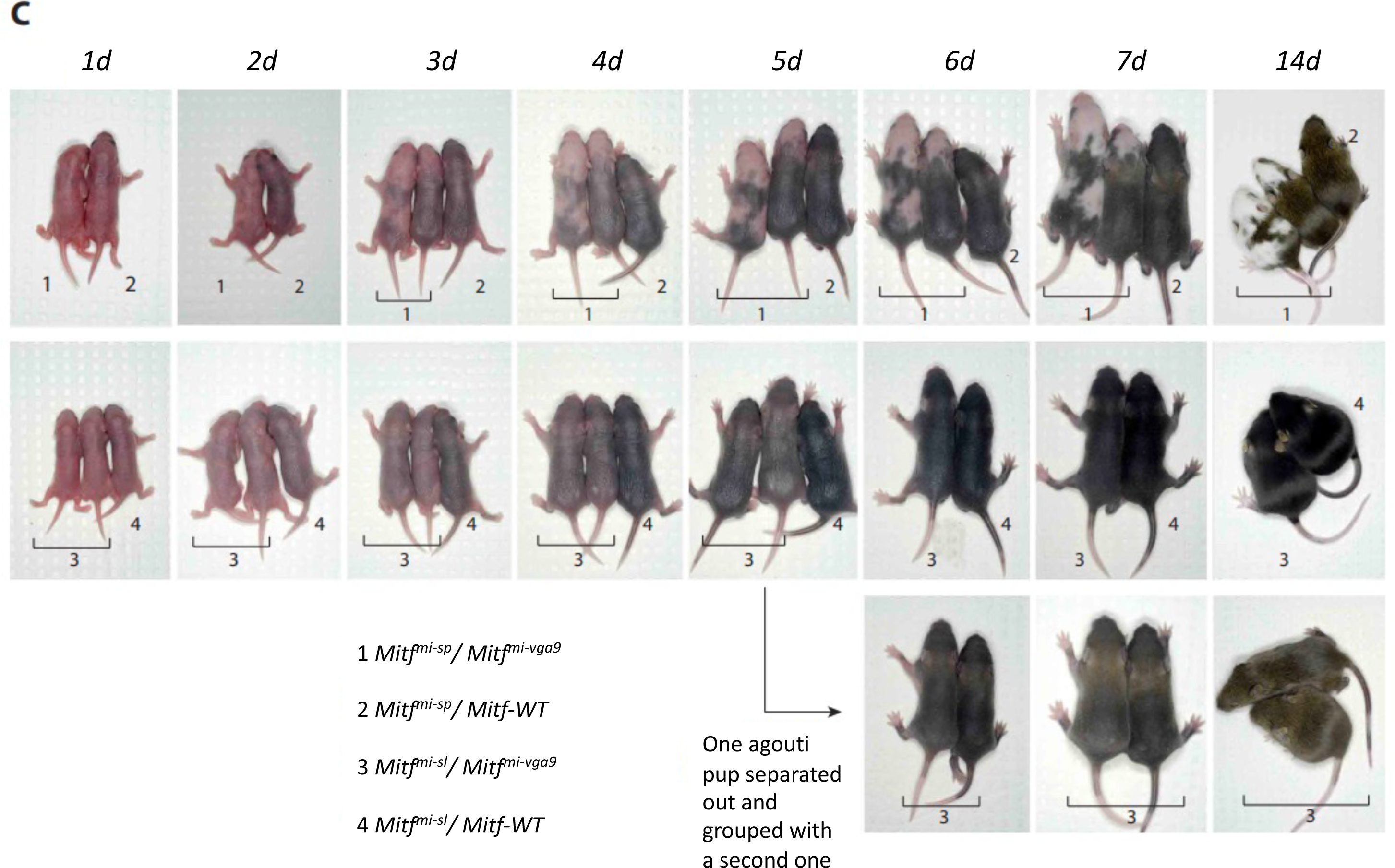
The *Mitf^mi-sl^* mutation results in delayed pigmentation. (A) Melanocytes differentiating from *ex vivo* neural crest cultures derived from *Mitf^mi-sl^* and *Mitf^mi-sp^* animals show that pigmentation is severely delayed in *Mitf^mi-sl^* homozygotes compared to *Mitf^mi-sp^*homozygotes. Nevertheless, melanoblasts are present, as evidenced by anti-MITF antibody staining (inset). (B) Homozygous *Mitf^mi-sl^* newborns show delayed onset of pigmentation compared to *Mitf^mi-sp^* homozygotes, particularly clearly visible at postnatal day-2 and 3 (p2 and p3). (C) No difference was observed in the onset of pigmentation in compound heterozygous condition with *Mitf^mi-vga9^*. Compare mice labeled #1 (*Mitf^mi-sp^*/*Mitf^mi-vga9^*) with mice labeled #3 (*Mit^fmi-sl^/Mit^fmi-vga9^*).

**Figure S4:**
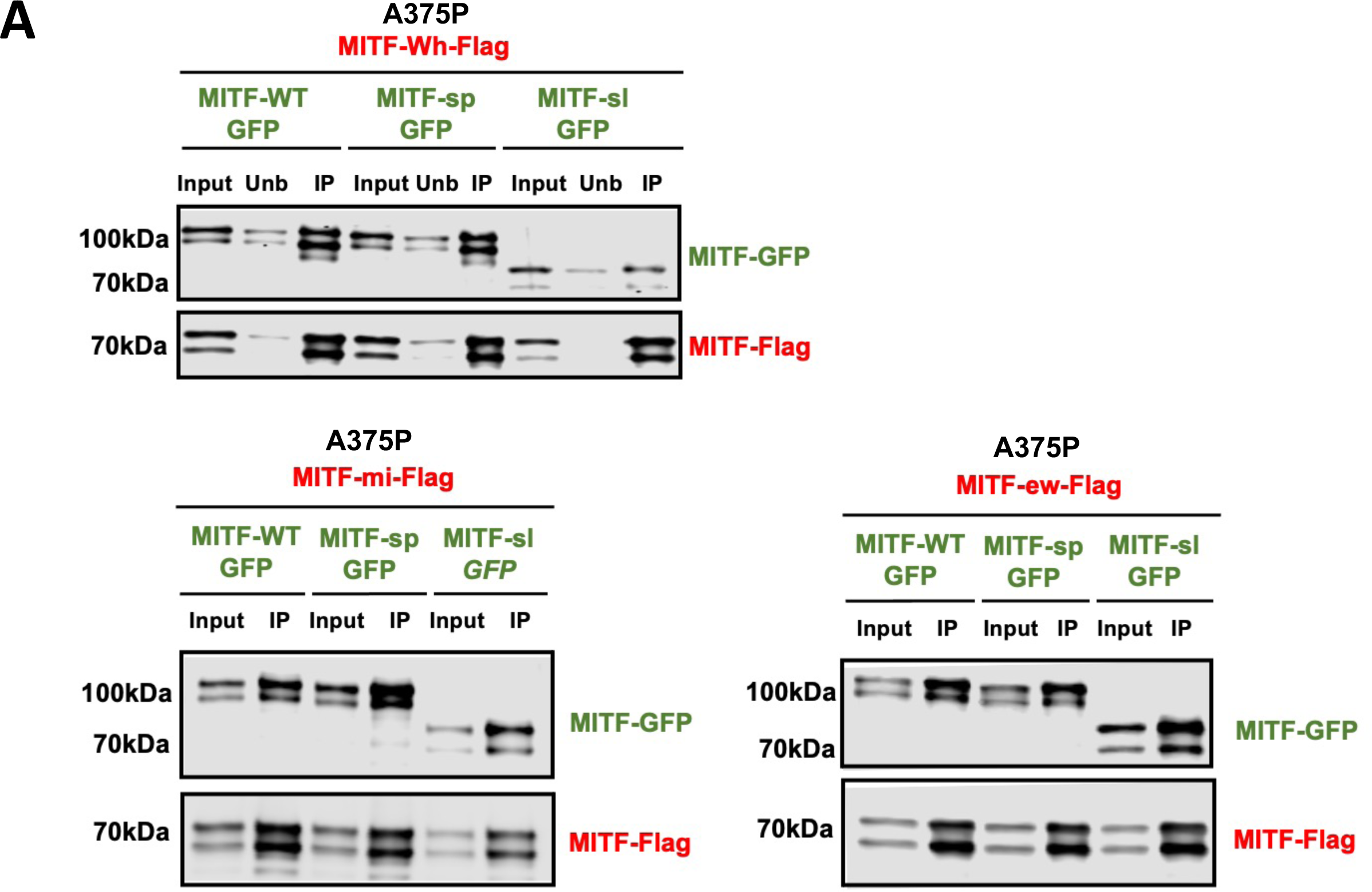
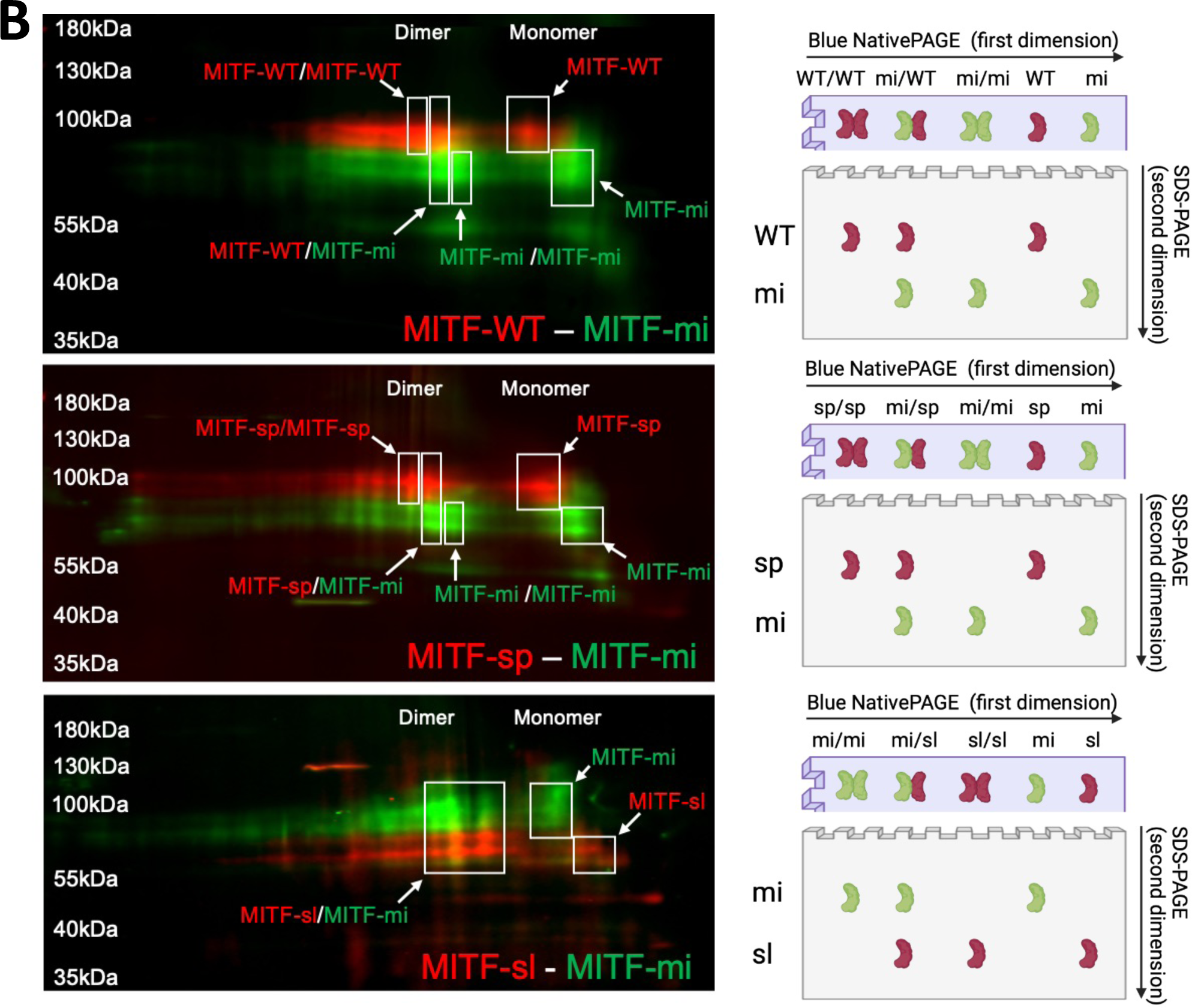
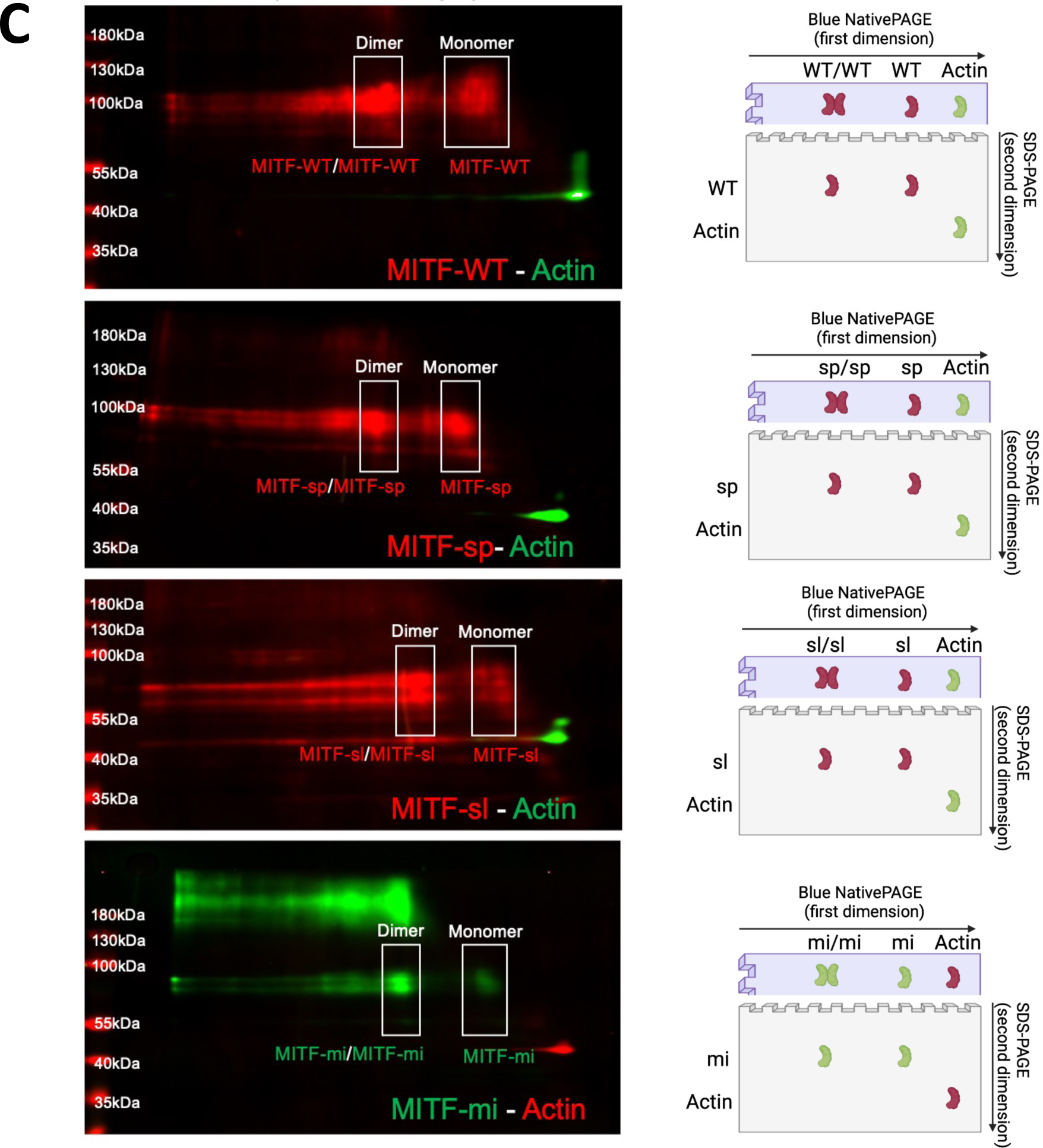
MITF-sl can dimerize with non-DNA-binding mutant MITF proteins. **(A)** Western blot analysis showing the results of a co-immunoprecipitation experiment. MITF-Wh-Flag, MITF-mi-Flag, or MITF-ew-Flag construct was cotransfected with either MITF-WT-GFP, MITF-sp-GFP, or MITF-sl-GFP construct in A375P melanoma cells. Co-immunoprecipitation (co-IP) of the whole cell lysate using FLAG-antibodies was performed, and proteins were visualized using FLAG and GFP antibodies. Input fraction (Input), Unbound fraction (Unb), and Immunoprecipitated fraction (IP) are indicated on the western blot. **(B) and (C)** Wild type and mutant MITF proteins visualized after a Blue native PAGE followed by the second dimension of SDS-PAGE. A375P melanoma cells transiently (B) co-expressing the MITF-mi-Flag protein (green) together with either MITF WT-GFP, MITF*-*sp*-*GFP, or MITF-sl-GFP proteins (red) or (C) expressing MITF-WT-Flag, MITF-sp-Flag, MITF-sl-Flag (red) or MITF-mi-Flag (green). We then performed Blue native PAGE 30 followed by a second dimension of SDS-PAGE. Actin in western blot (C) was visualized by using mouse or rabbit Actin antibodies and then anti-mouse (red) or anti-rabbit (green) secondary antibodies. The cartoon (on the left) illustrates the migration patterns of dimeric and monomeric proteins after undergoing Blue Native PAGE and SDS-PAGE, demonstrating their relative mobility and separation in the gel.

**Figure S5:**
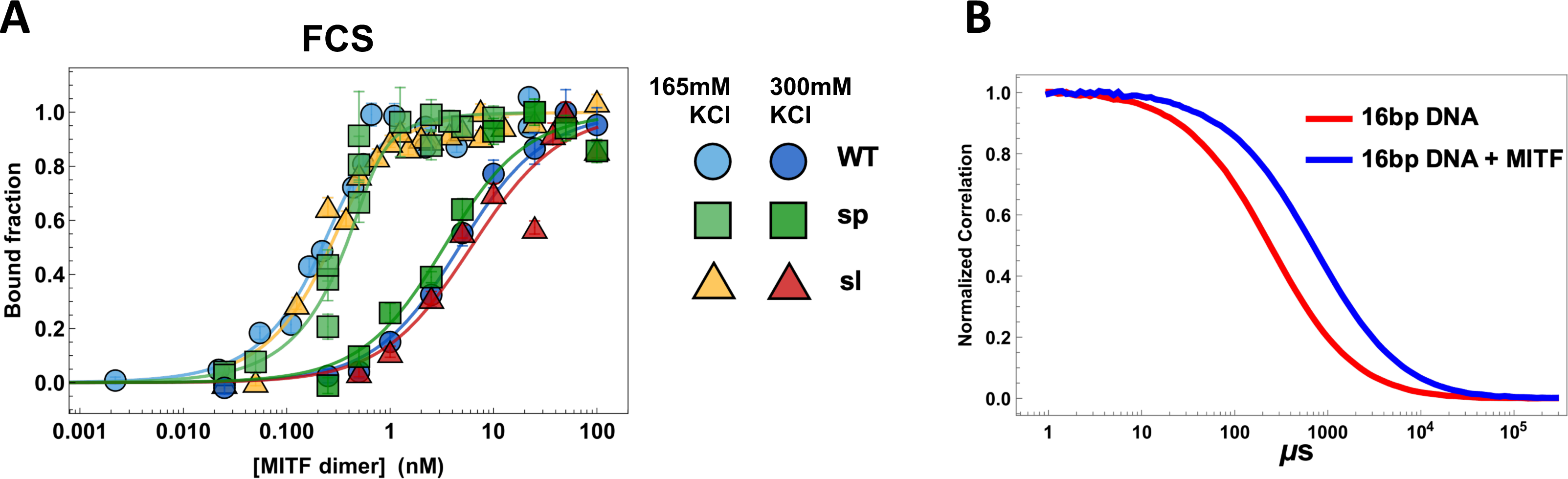
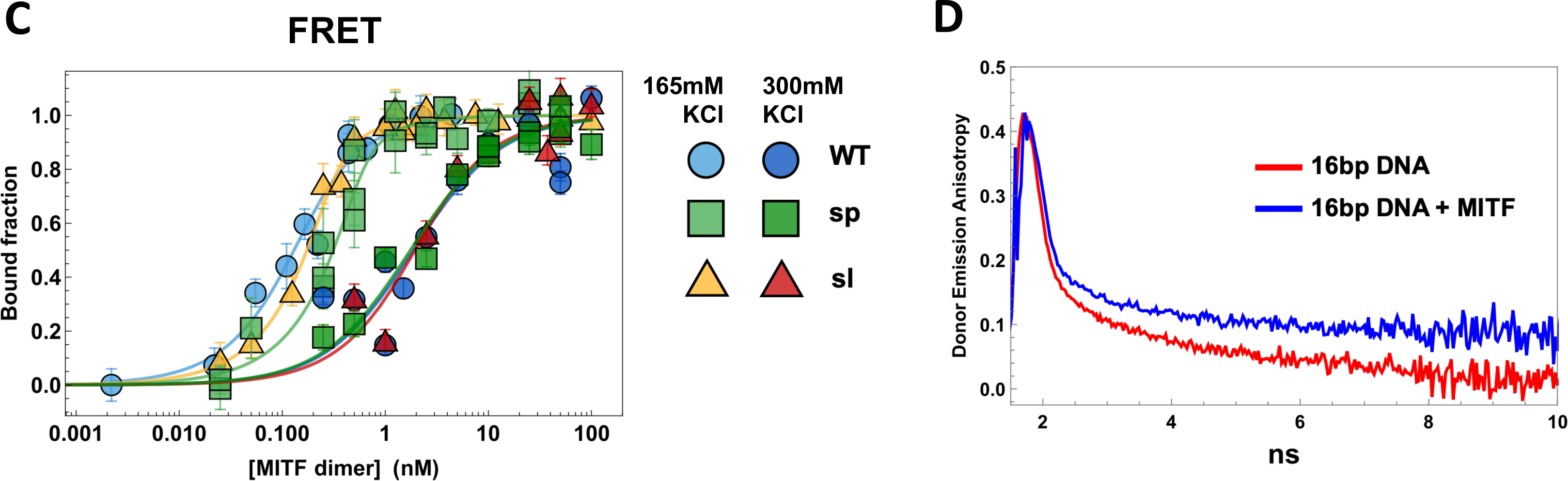
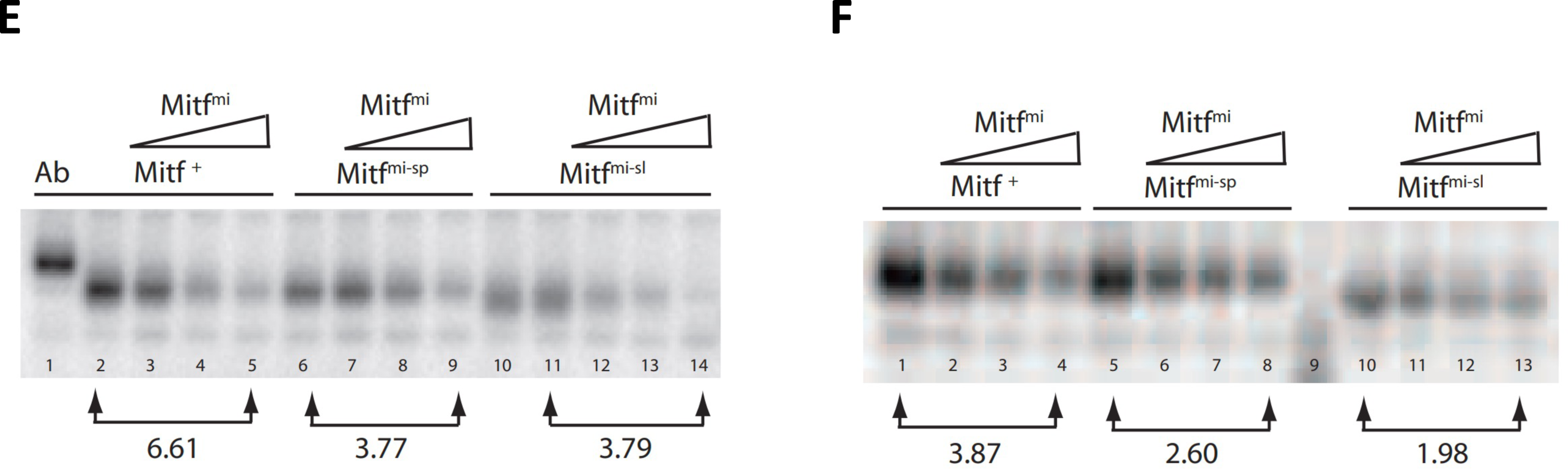
The MITF-sl protein has similar DNA binding affinity, however, prefers to form dimers compared to MITF-WT and MITF-sp. **(A)** and **(C)** DNA binding curves of recombinantly expressed human MITF-WT, MITF-sp, and MITF-sl protein to M-box probe measured by Fluorescence Correlation Spectroscopy (FCS) (A) and mean FRET (C) at 165 mM KCl and 300 mM KCl. MITF-WT protein in light and dark blue circles, MITF-sp light and dark green boxes, and MITF-sl in yellow and red triangles. Error bars represent two standard deviations of fit error at each point. **(B)** Normalized donor-acceptor fluorescence cross-correlation curves of 16bp M-Box DNA alone (red) and with 100 nM WT MITF added (blue), both at 300 mM KCl. **(D)** Inverted time-correlated donor emission anisotropy of 16bp DNA alone (red) and in the presence of 100 nM MITF WT (blue), both at 300 mM KCl. **(E)** and **(F)** Electrophoretic mobility shift assays were performed using the M-box sequence (5’-AAAGTCAGTCATGTGCTTTTCAGA-3’) as a probe. (E) MITF-WT, MITF-sp, and MITF-sl proteins were expressed using the TNT (Promega) system alone (lanes 1, 2, 6, 10, and 11) or co-expressed with the dominant negative MITF-mi protein (lanes 3-5, 7-9, and 12-14) and then incubated with the labeled probe. The binding is specific since the presence of the C5 monoclonal MITF antibody which recognizes the N-terminus of Mitf results in a supershift (Ab). (F) The same experiment as in (E), except that the proteins were translated separately and then incubated for 30 minutes in the presence of DNA to allow heterodimerization before performing the mobility shift experiment.

**Figure S6:**
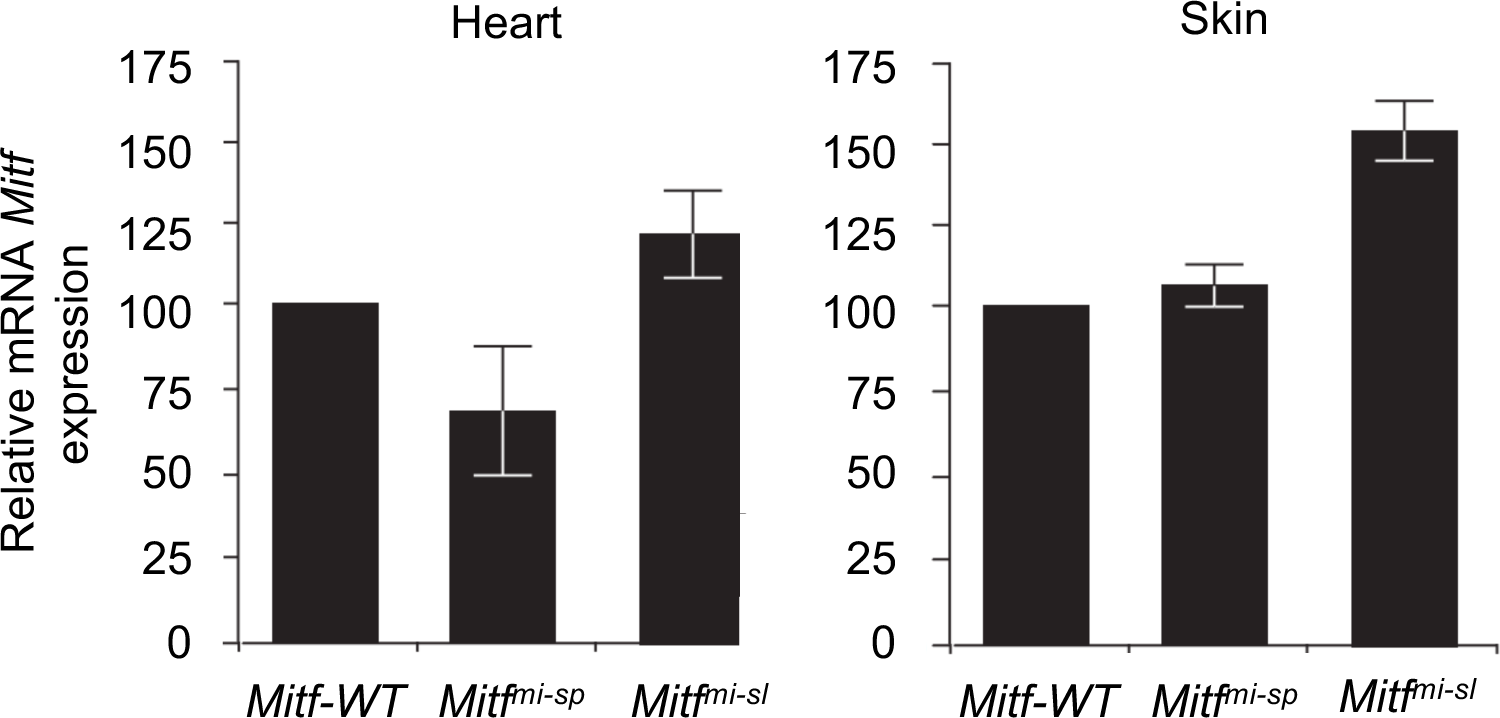
Relative expression of *Mitf* mRNA in wild type, *Mitf^mi-sp^/Mitf^mi-sp^* and *Mitf^mi-sl^*/*Mitf^mi-sl^* heart and skin determining by qRT-PCR.

**Figure S7:**
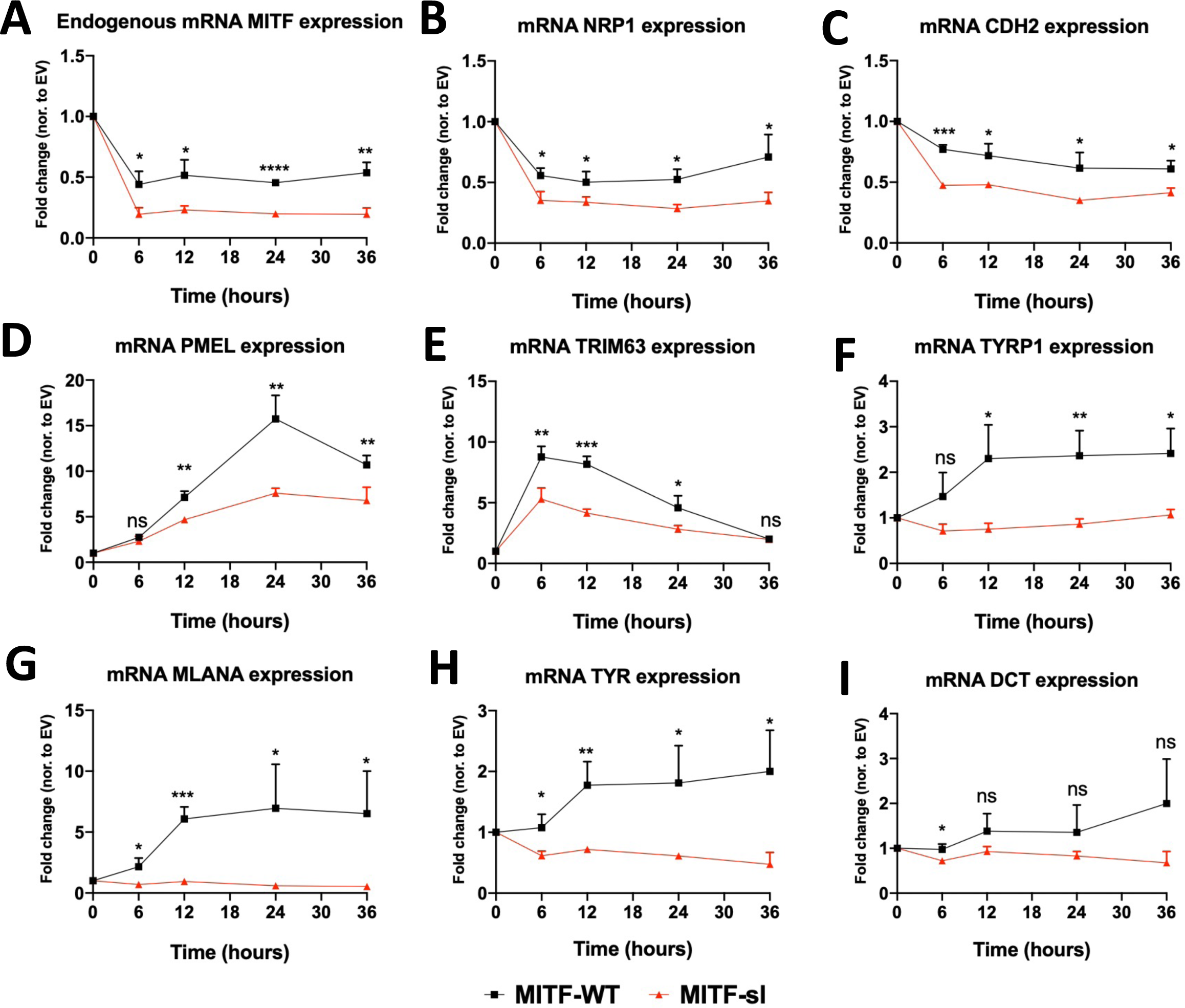
MITF-sl protein is a less potent activator than MITF-WT. RT-qPCR analysis of **(A)** endogenous mRNA MITF and mRNA MITF target genes: **(B)** NRP1, **(C)** CDH2, **(D)** PMEL, **(E)** TRIM63, **(F)** TYRP1, **(G)** MLANA, **(H)** TYR, and (**I)** DCT in the dox-inducibleA375P overexpressing cells. The cells were treated with doxycycline for 6, 12, 24, and 36 hours to induce MITF expression at the same level before harvesting. Expression was normalized to EV-FLAG-HA cell lines and then to the proportion of MITF proteins retained in the nucleus. Error bars represent SEM of at least three independent experiments. Statistically significant differences (Student’s t-test) are indicated by * p< 0.05, *p< 0.05, ** p < 0.01, *** p < 0.001, **** p < 0.00, and ns not significant.

**Figure S8:**
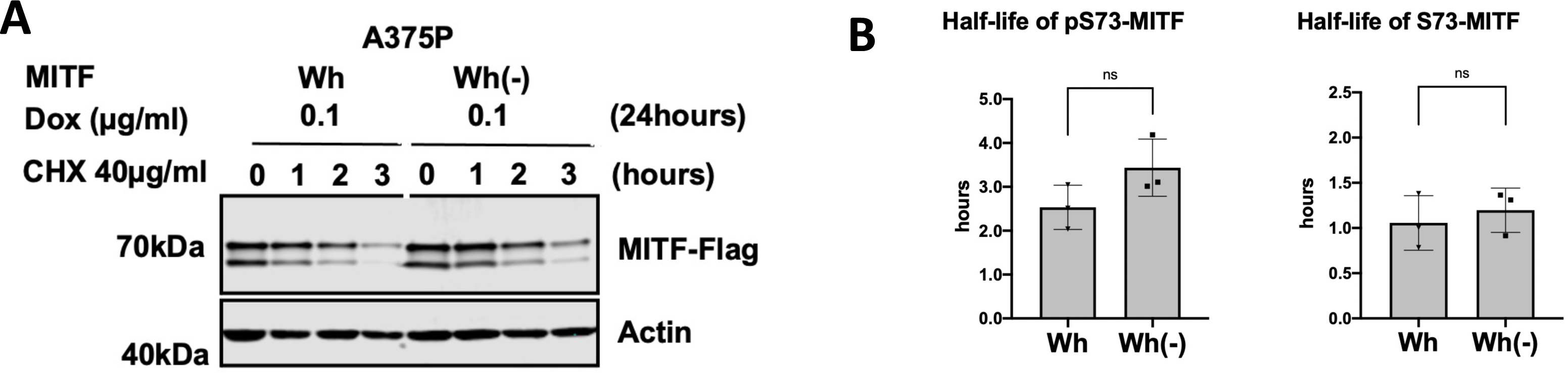
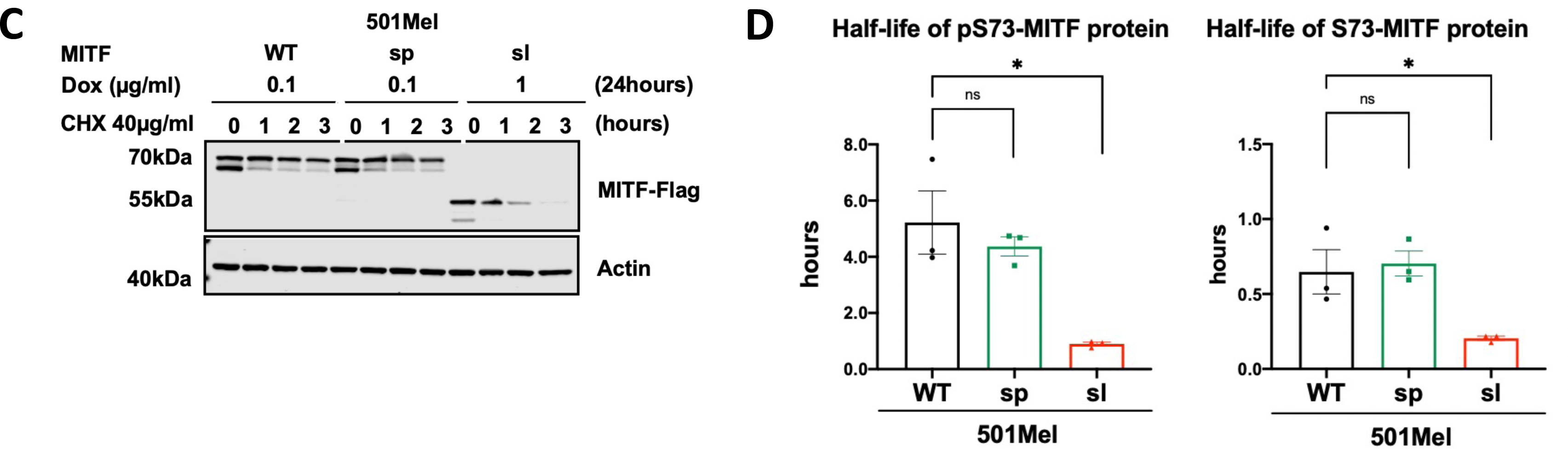
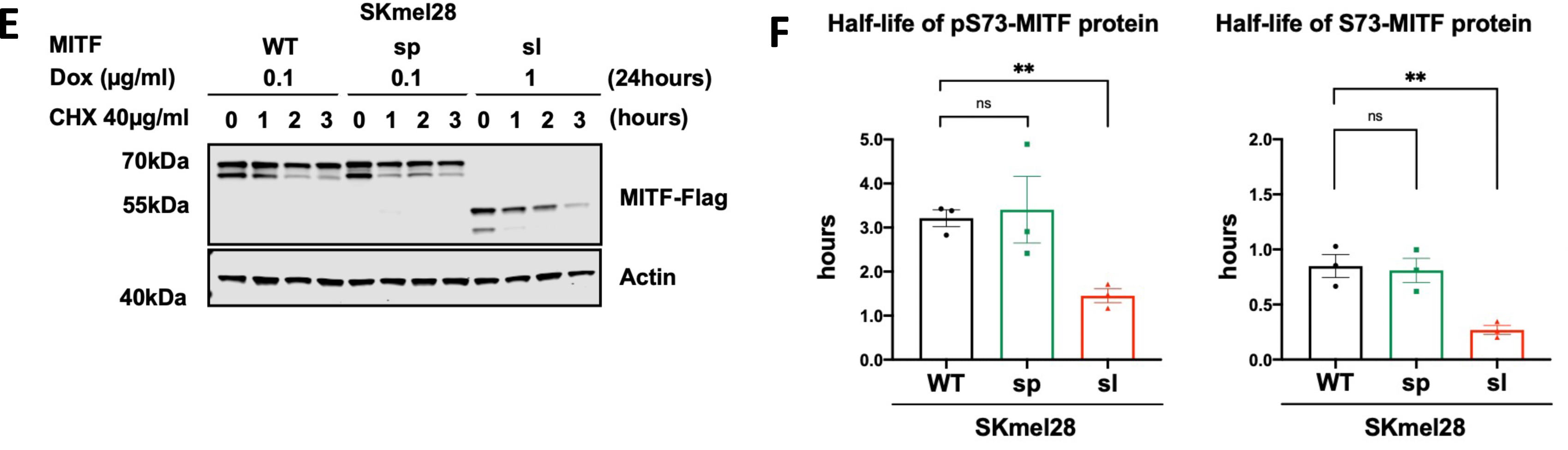
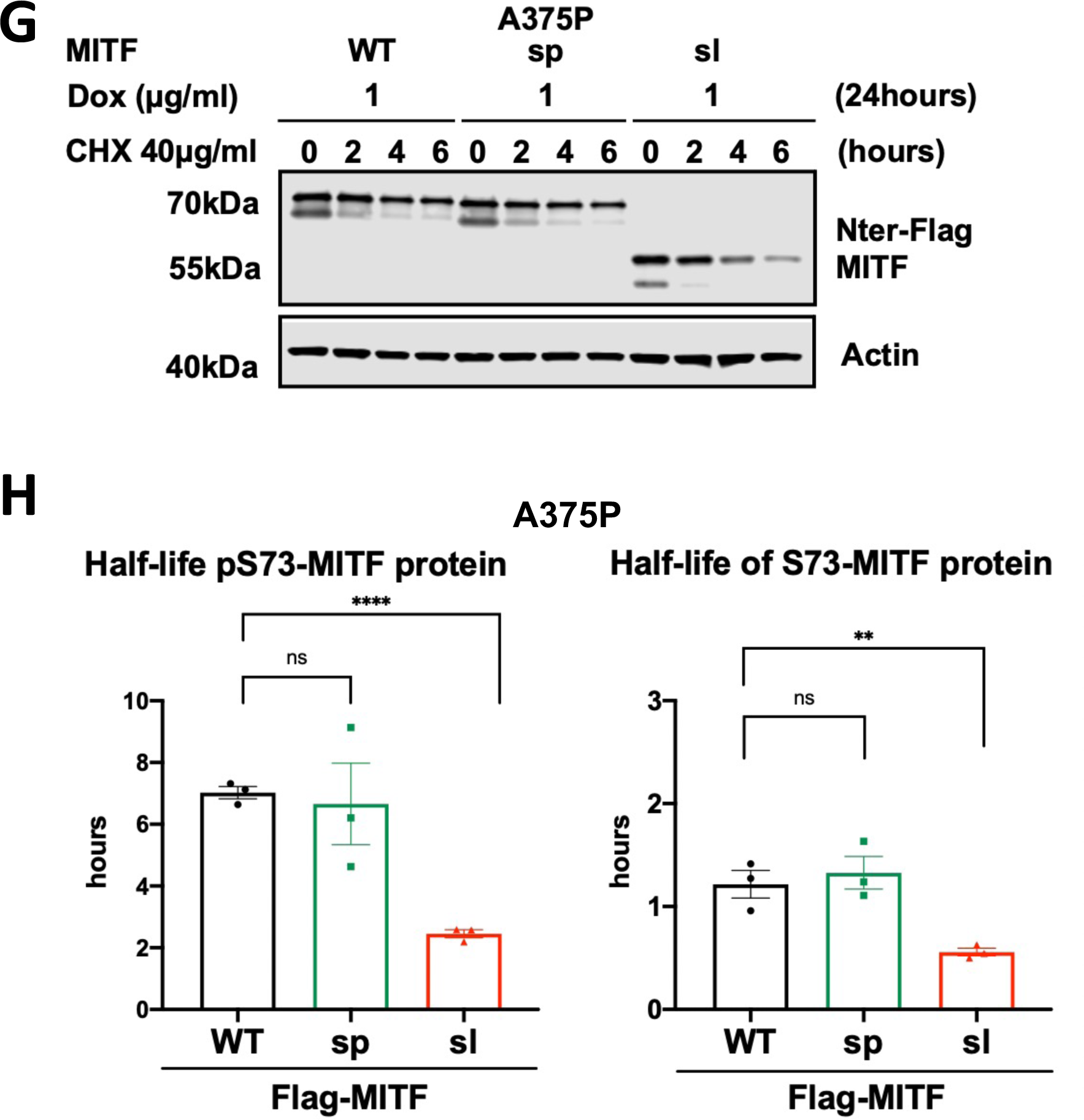
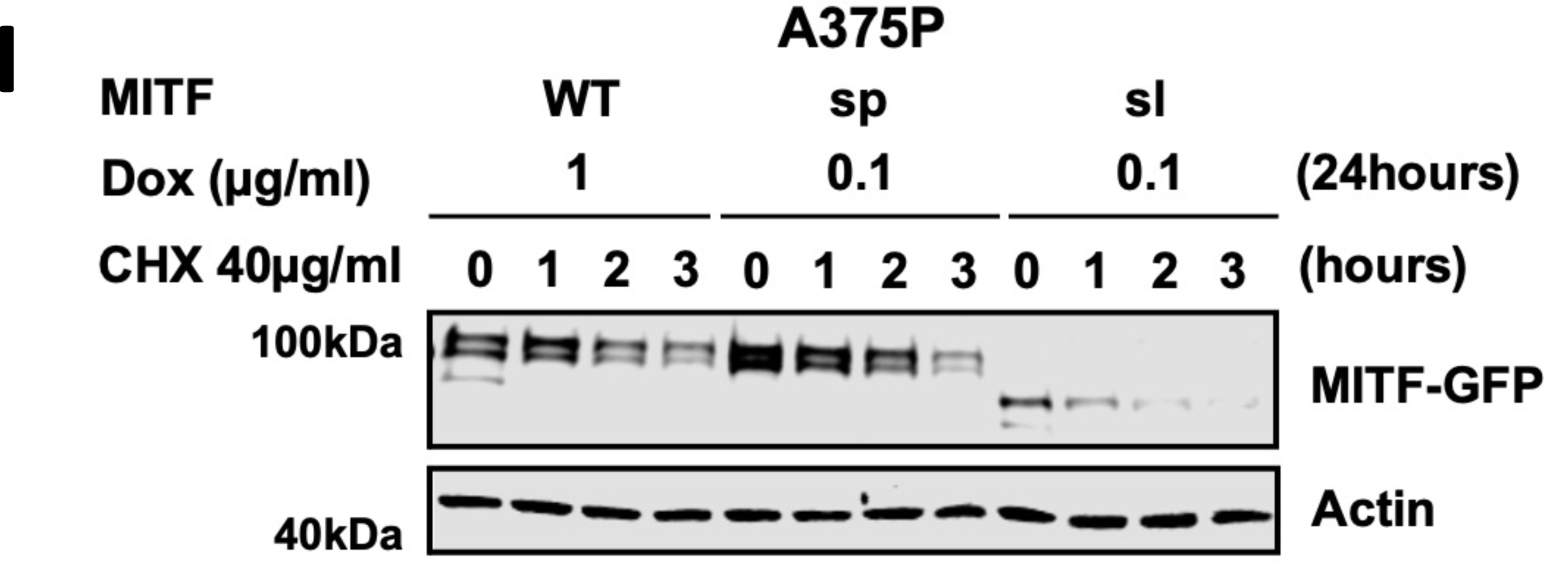
Exon 6A and the tags did not affect MITF stability. **(A), (C), (E), (G),** and **(I)** Western blot analysis of the wild type and mutant MITF proteins with different tags at either N-terminus or C-terminus. The dox-inducible A375P cells were treated with doxycycline for 24h to express the indicated MITF proteins before treating them with 40 µg/ml CHX for 0, 1, 2, and 3 hours. The MITF proteins were then compared by western blot using FLAG antibody. Actin was used as a loading control. The band intensities were quantified using ImageJ software. **(B), (D), (F), (H)** Half-life analysis of the pS73- and S73-MITF proteins over time after CHX treatment. The MITF protein levels relative to T0 were calculated, and non-linear regression analysis was performed. Error bars represent SEM of at least three independent experiments. Statistically significant differences (Student’s t-test) are indicated by * p< 0.05, *p< 0.05, ** p < 0.01, *** p < 0.001, **** p < 0.00, and ns not significant.

**Figure S9:**
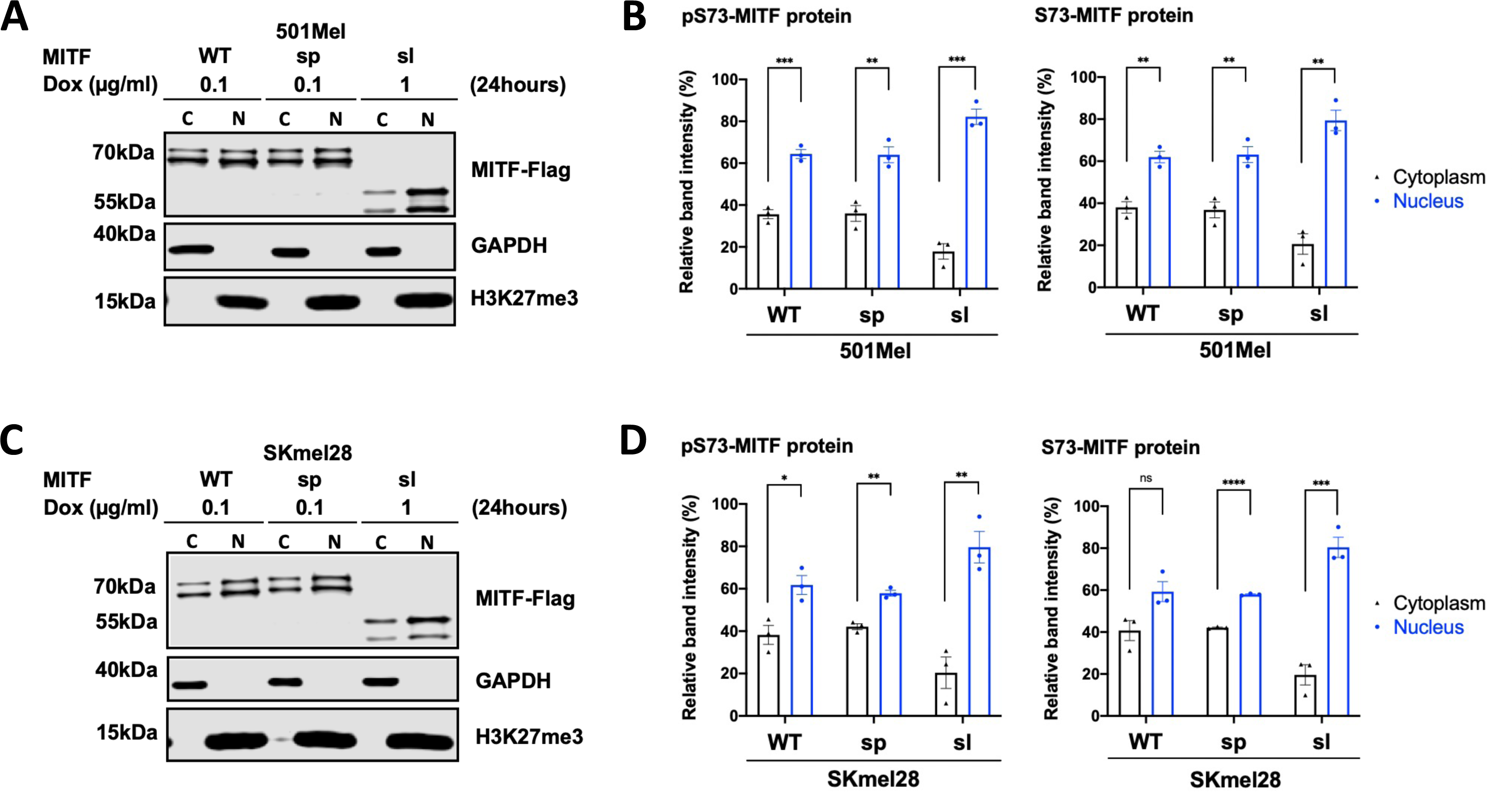
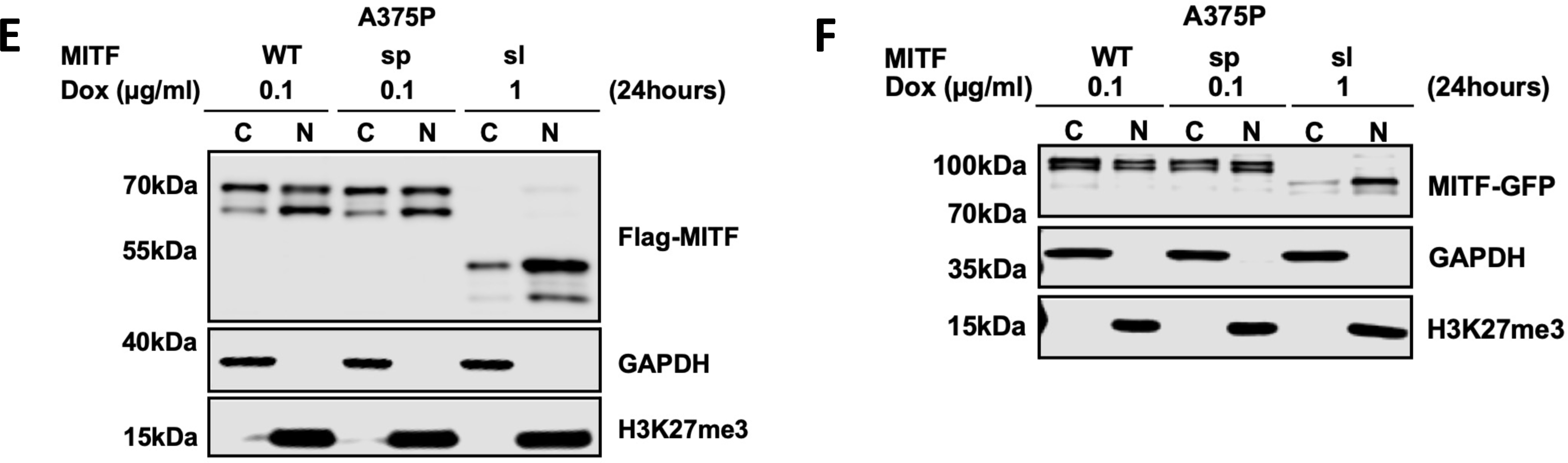
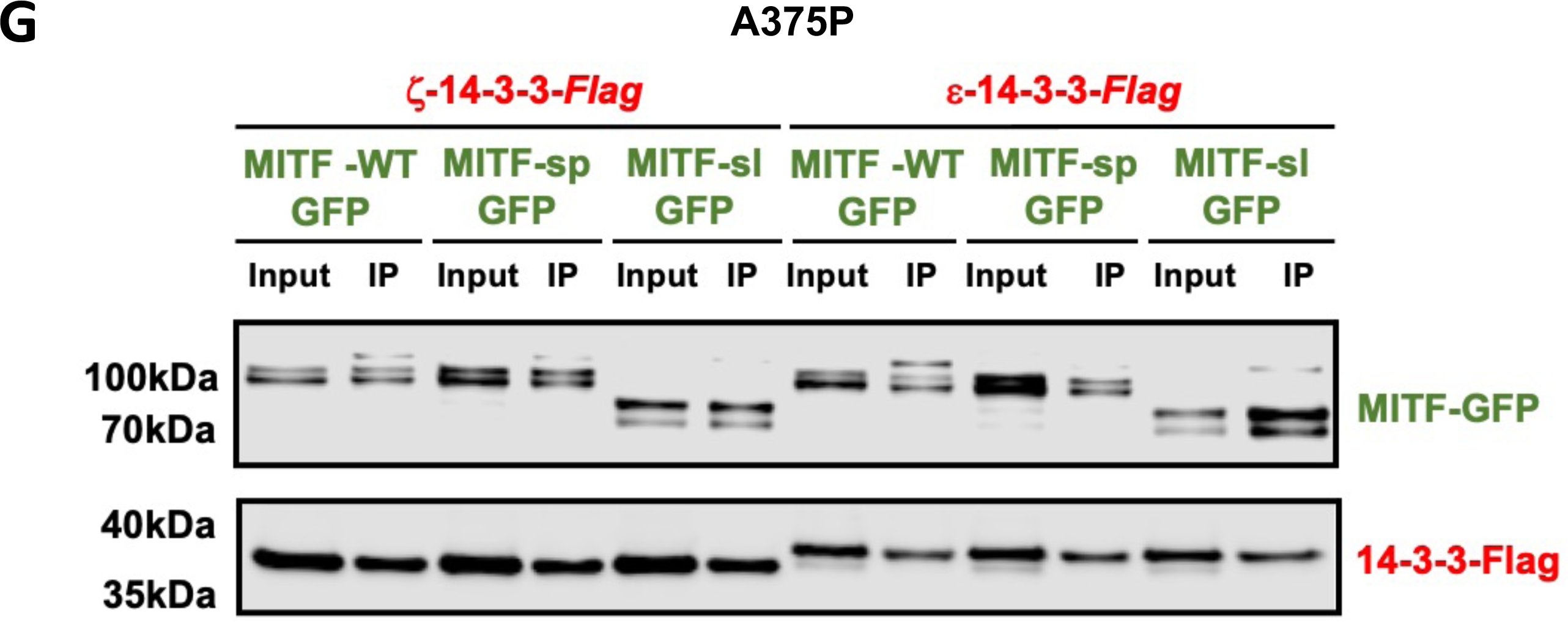
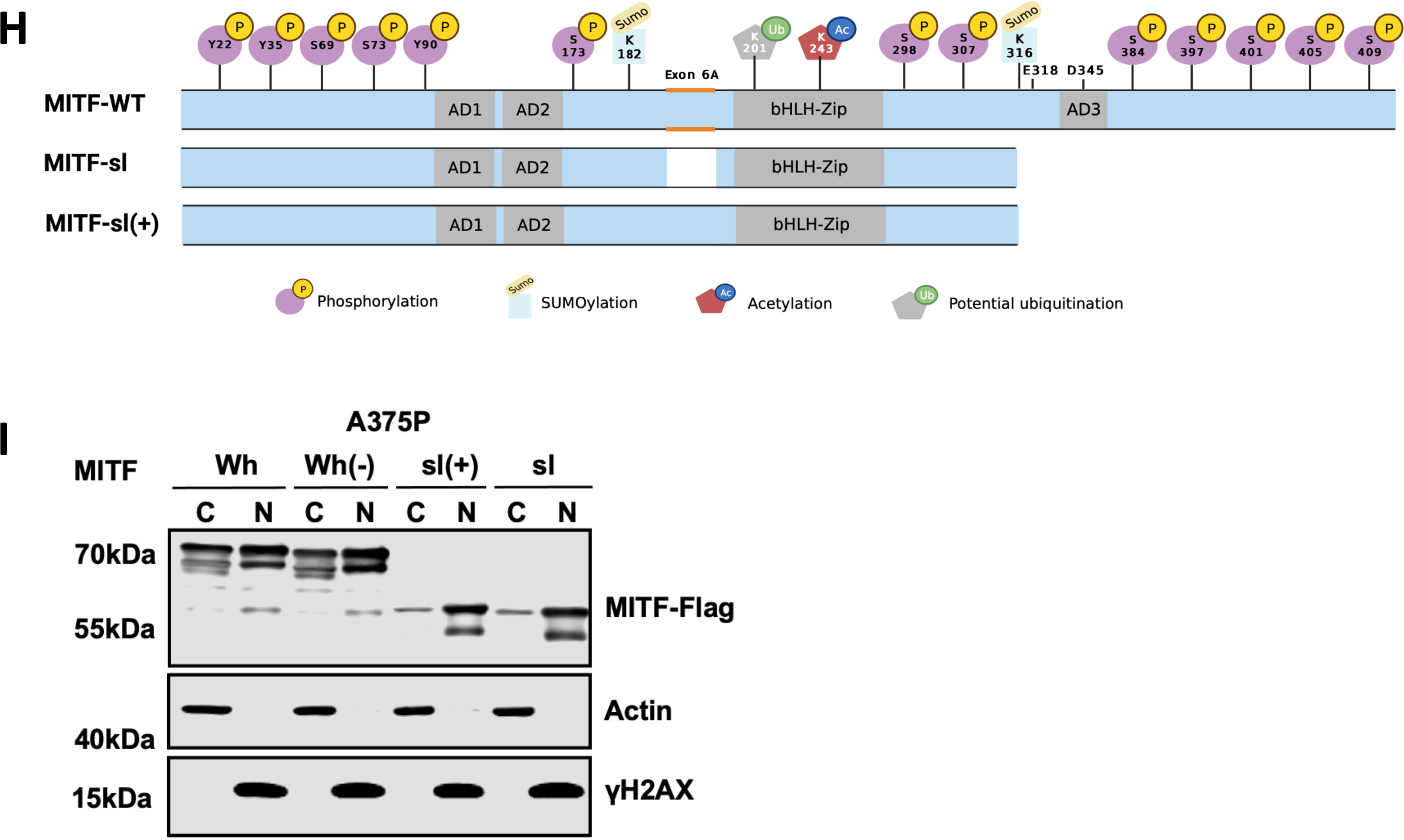
Exon 6A and the tags did not affect MITF nuclear localization. **(A), (C), (E), (F),** and **(I)** Western blot analysis of cytoplasmic (C) and nuclear (N) fractions from either 501Mel (A), SKMel28 (C) or A375P (E, F, G) melanoma cells induced for 24 hours to overexpress the MITF mutant proteins with either Flag-tag at C-terminus (A, C, G) or Flag-tag at N-terminus (E) or GFP-tag at C-terminus (F) visualized using either FLAG or GFP antibody. GAPDH and H3K27me3 were loading controls for cytoplasmic and nuclear fractions, respectively. **(B)** and **(D)** The intensities of the pS73- and S73-MITF proteins from the Western blots in (A) and (C), respectively, were quantified separately with *ImageJ* software and are depicted as percentages of the total amount of protein present in the two fractions. Error bars represent SEM of three independent experiments. Statistically significant differences (Student’s t-test) are indicated by * p< 0.05, *p< 0.05, ** p < 0.01, *** p < 0.001, **** p < 0.00, and ns not significant. **(G)** Western blot analysis showing the results of a co-immunoprecipitation experiment. MITF^mi-sp^-GFP, MITF^mi-sl^-GFP, or MITF-WT-GFP construct were cotransfected with either ε-14-3-3-Flag or ζ-14-3-3-Flag in A375P melanoma cells. Co-immunoprecipitation (co-IP) of the whole cell lysate using FLAG-antibodies was performed, and proteins were visualized using FLAG and GFP antibodies. Input fraction (Input), unbound fraction (Unb), and an immunoprecipitated fraction (IP) are indicated on the Western blot. **(H)** Schematic of the MITF-sl(+) mutant construct, which was generated by introducing exon 6A.

**Figure S10:**
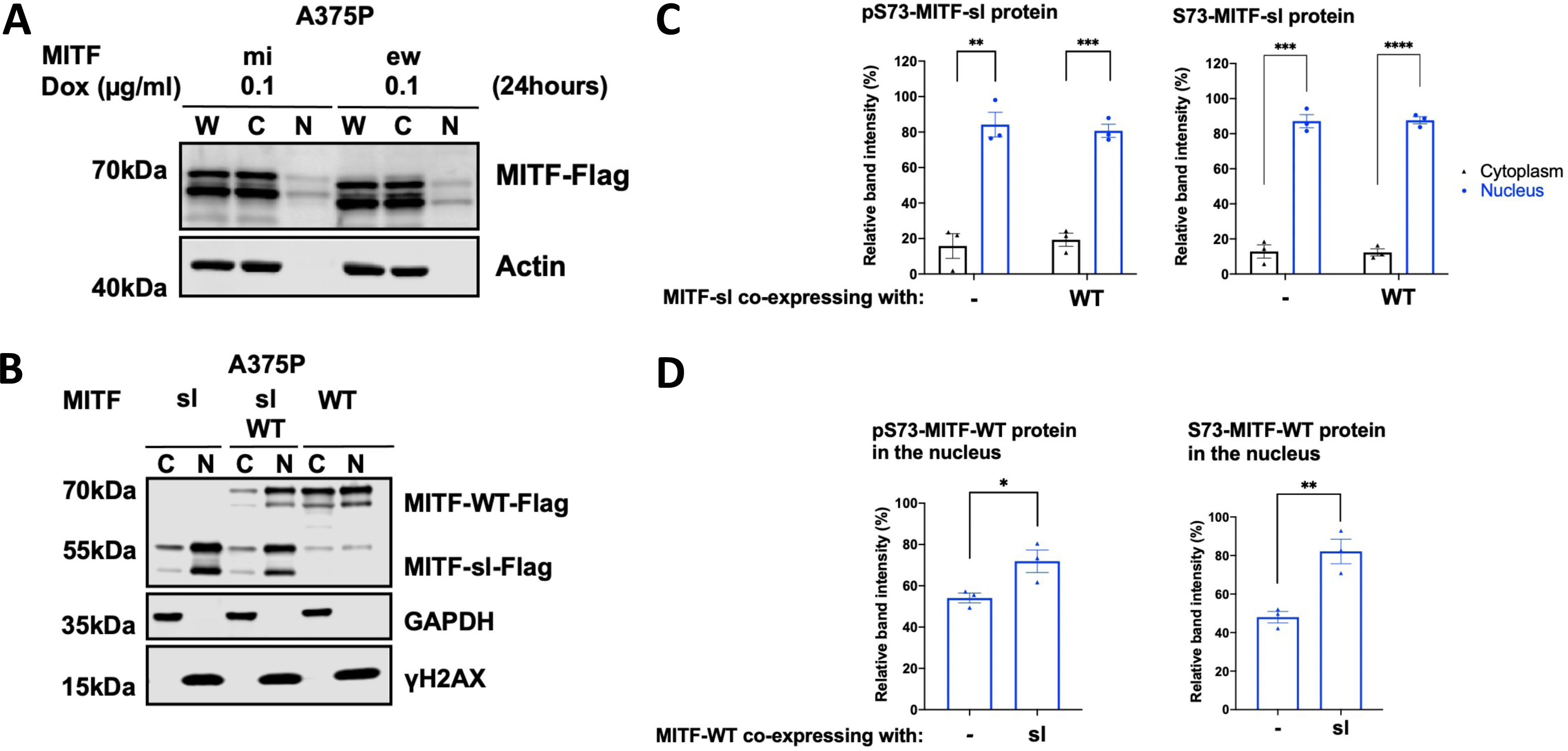
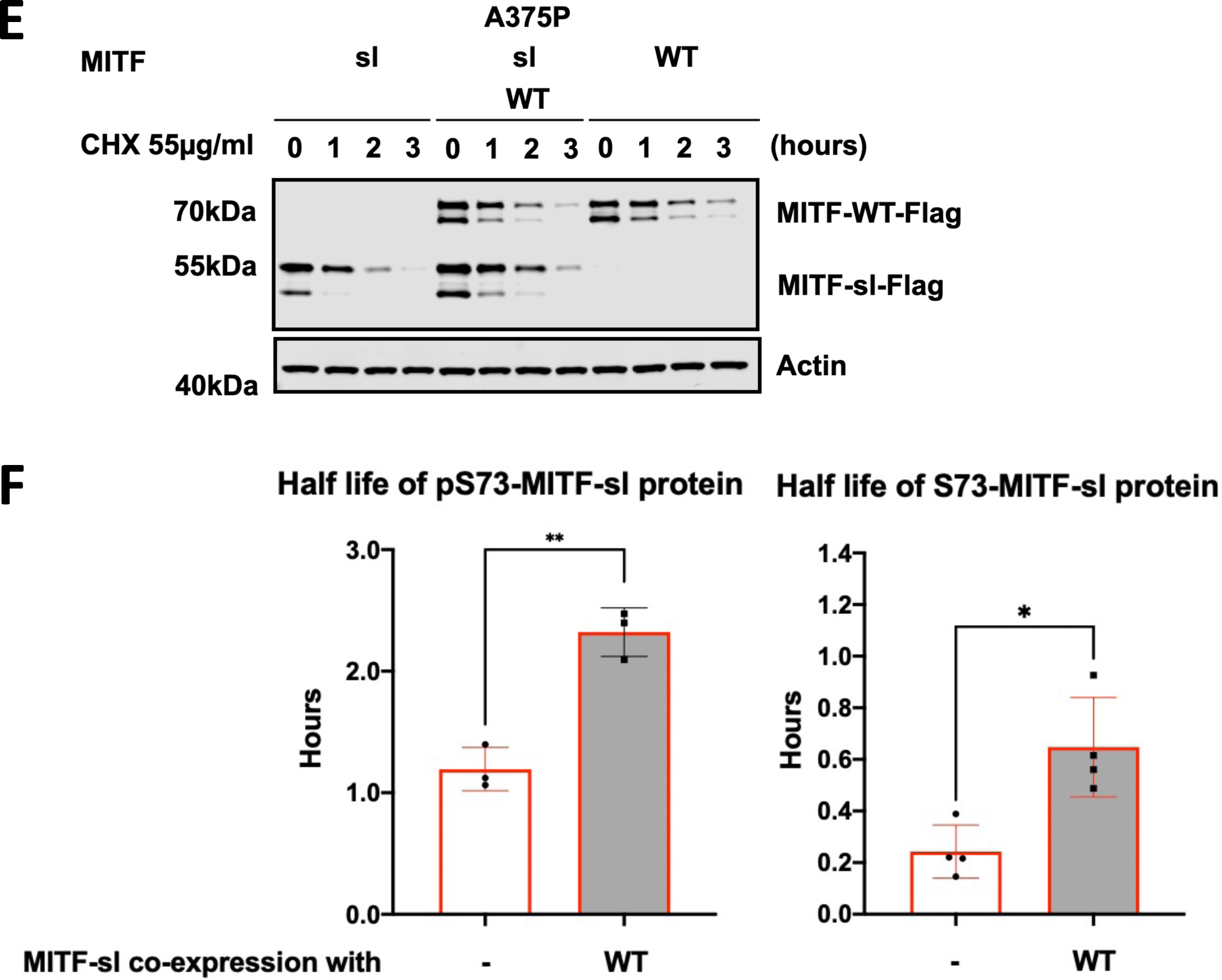
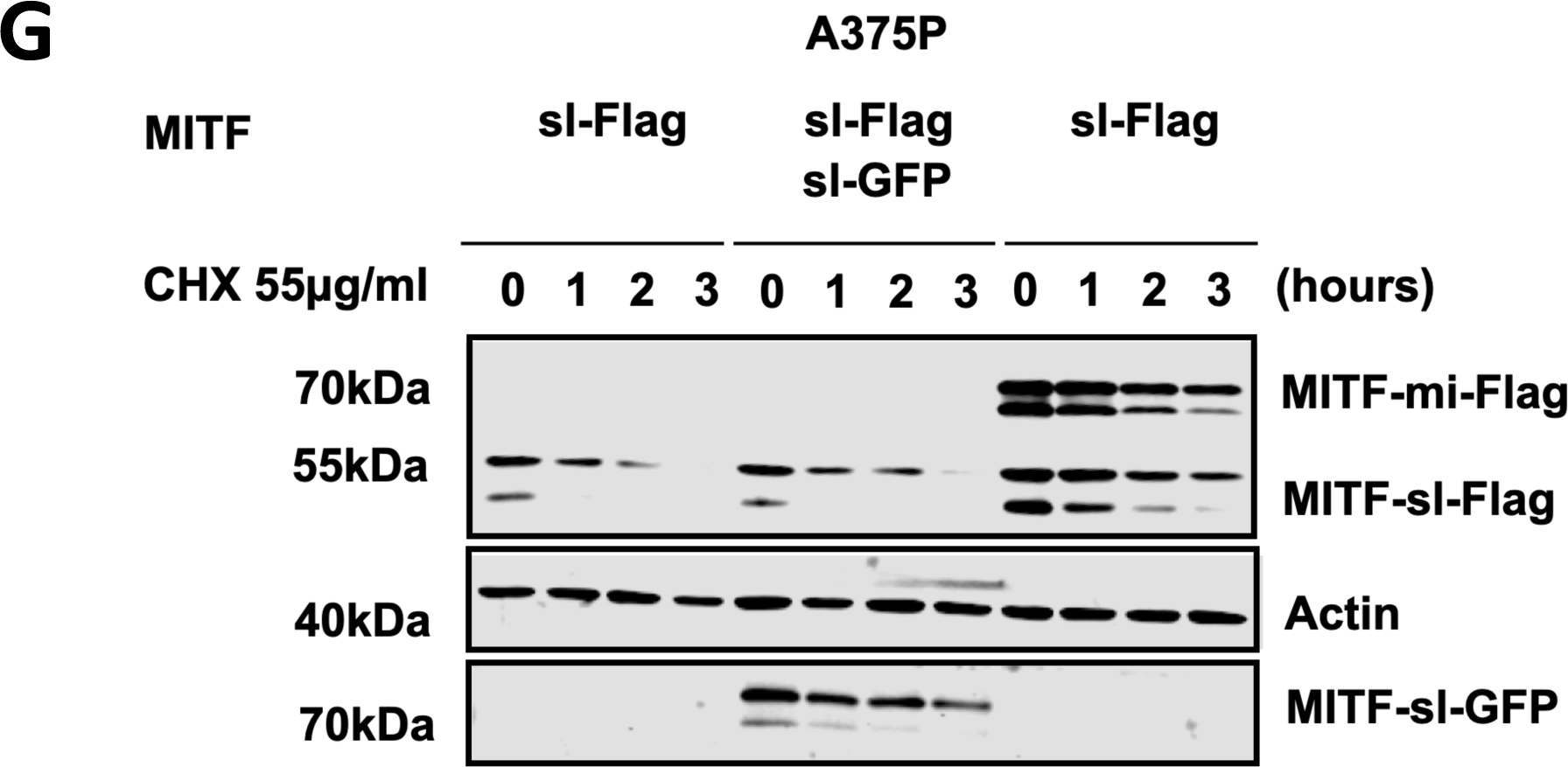
MITF-sl enables nuclear localization of its dimer partner. **(A)** Western blot analysis of subcellular fractions isolated from A375P melanoma cells induced for 24 hours to overexpress different MITF mutant proteins. MITF-mi and MITF-ew proteins in whole cell lysate (W), cytoplasmic (C) and nuclear (N) fractions were visualized using FLAG antibody. Actin was used as a loading control for the cytoplasm. **(B)** Western blot analysis of subcellular fractions isolated from A375P cells transiently co-overexpressing the MITF*-*sl protein with MITF-WT. MITF proteins in cytoplasmic (C) and nuclear (N) fractions were visualized using FLAG antibody. GAPDH and γH2AX were loading controls for cytoplasmic and nuclear fractions, respectively. **(C)** Intensities of the pS73- and S73-MITF-sl proteins in the cytoplasmic and nuclear fractions of A375P cells transiently co-overexpressing the MITF*-*sl protein together with MITF-WT were quantified separately with ImageJ software and are depicted as percentages of the total amount of protein present in the two fractions. Error bars represent SEM of three independent experiments. Statistically significant differences (Student’s t-test) are indicated by * p< 0.05, *p< 0.05, ** p < 0.01, *** p < 0.001, **** p < 0.00, and ns not significant. **(D)** Intensities of the pS73- and S73-MITF-WT proteins in the nuclear fraction of A375P cells transiently co-overexpressing the MITF*-*sl protein together with MITF-WT proteins were quantified with ImageJ software and are depicted as percentages of the total amount of protein present in the two fractions. Error bars represent SEM of three independent experiments. Statistically significant differences (Student’s t-test) are indicated by * p< 0.05, *p< 0.05, ** p < 0.01, *** p < 0.001, **** p < 0.00, and ns not significant. **(E)** Western blot analysis of the MITF-sl protein in the presence of MITF-WT. The A375P cells were transiently co-transfected with MITF-sl and MITF-WT for 24 hours before being treated with 55 µg/ml CHX. The amount of MITF protein was then compared by western blot using FLAG antibody. Actin was used as a loading control. The band intensities were quantified using ImageJ software. **(F)** Non-linear regression (one-phase decay) and half-life analysis of the pS73- and S73-MITF proteins over time after CHX treatment. The MITF protein levels relative to T0 were calculated, and non-linear regression analysis was performed. Error bars represent SEM of at least three independent experiments. Statistically significant differences (Student’s t-test) are indicated by *, p< 0.05. **(G)** Western blot analysis of the MITF-sl protein with Flag-tag at C terminus in the presence of either MITF-sl-GFP tagged at C-end or MITF-mi-Flag at C-end. The A375P cells were transiently co-transfected with MITF-sl-Flag and the MITF mutant proteins for 24 hours before being treated with 55 µg/ml CHX. The MITF protein was then compared by western blot using FLAG antibody. Actin was used as a loading control. The band intensities were quantified using ImageJ software.

**Figure S11:**
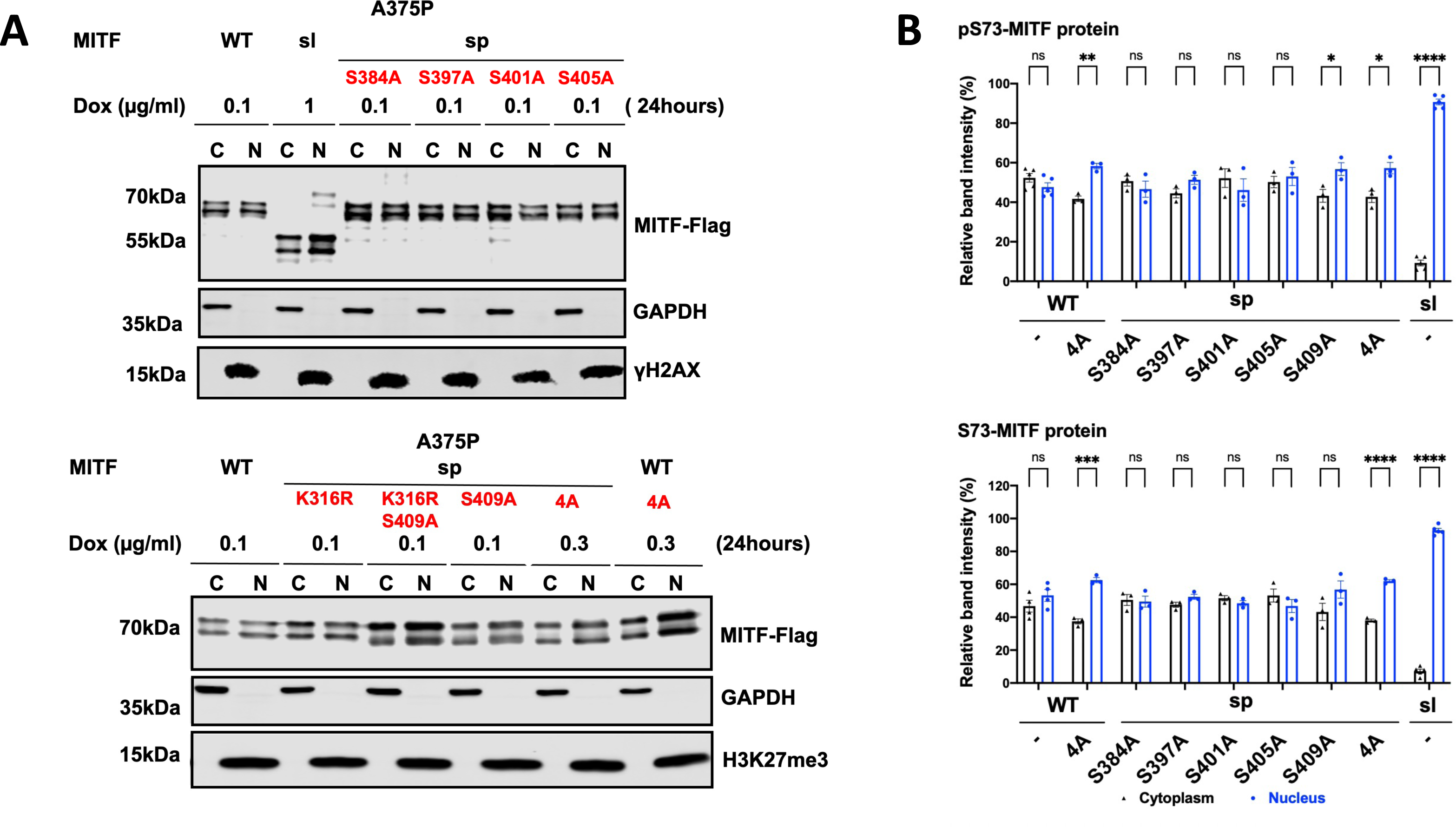
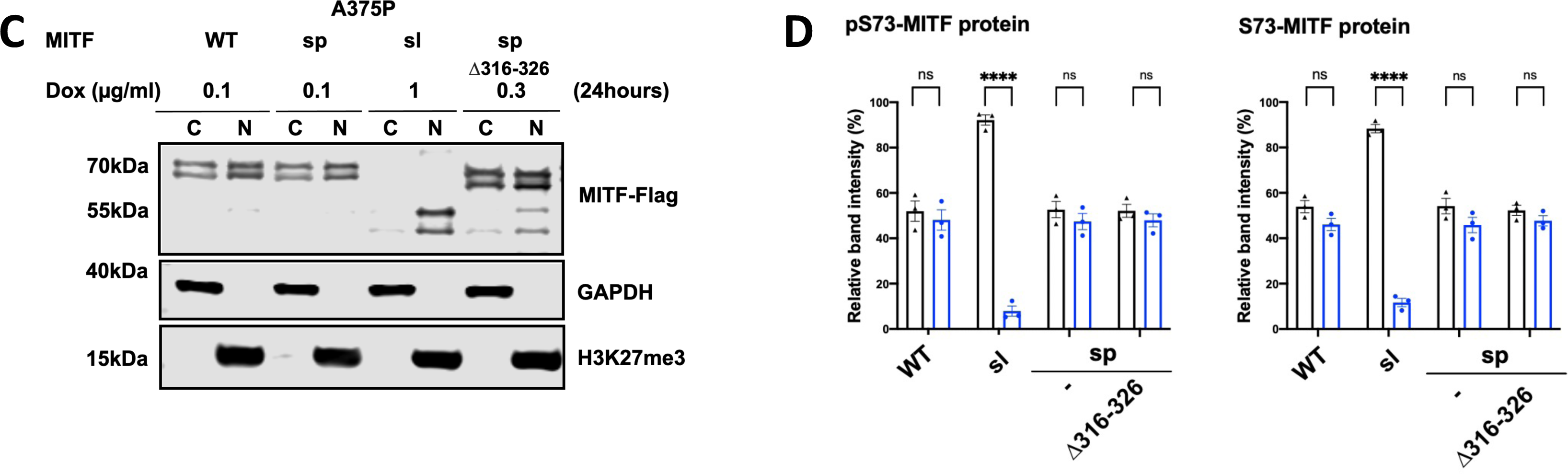
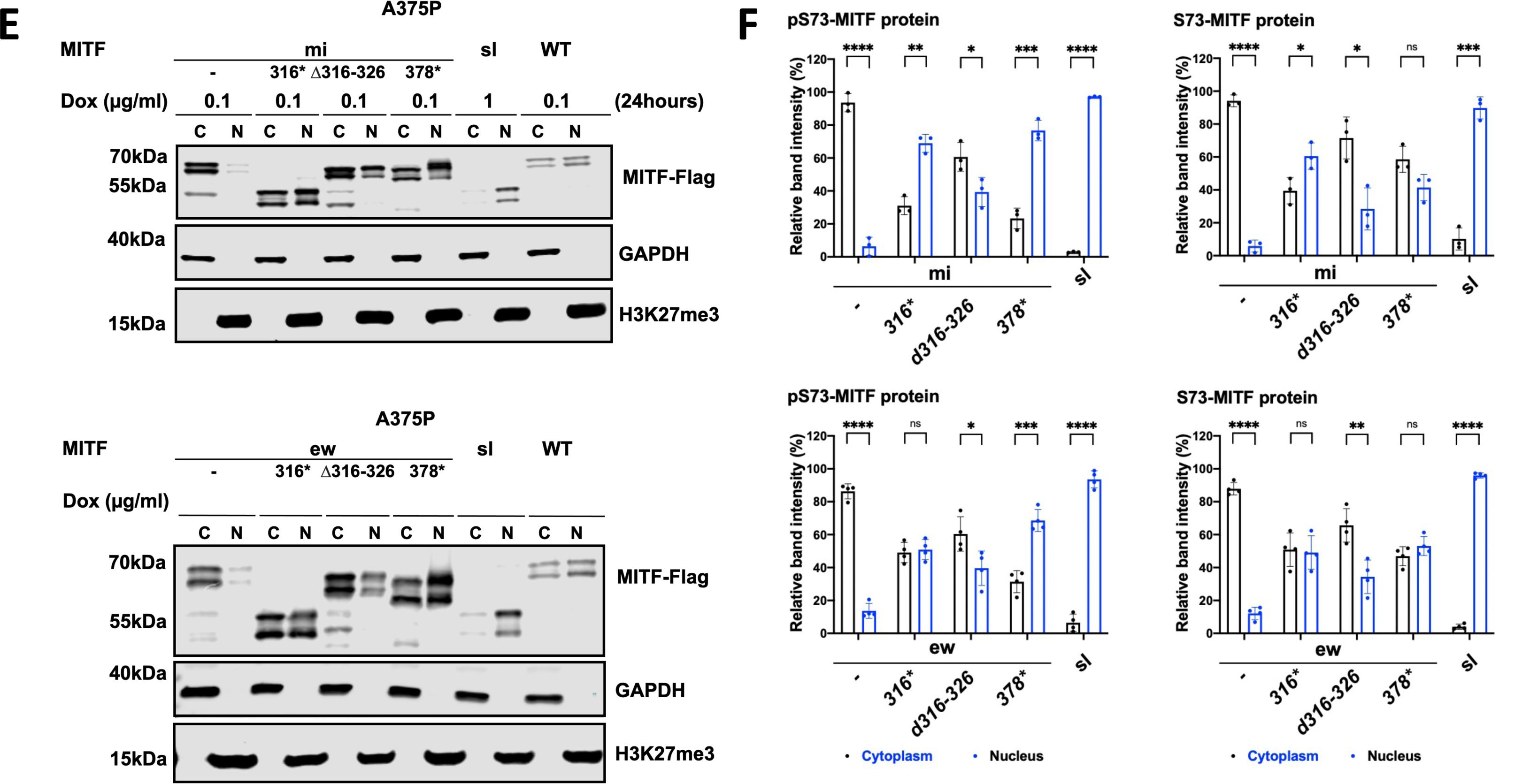
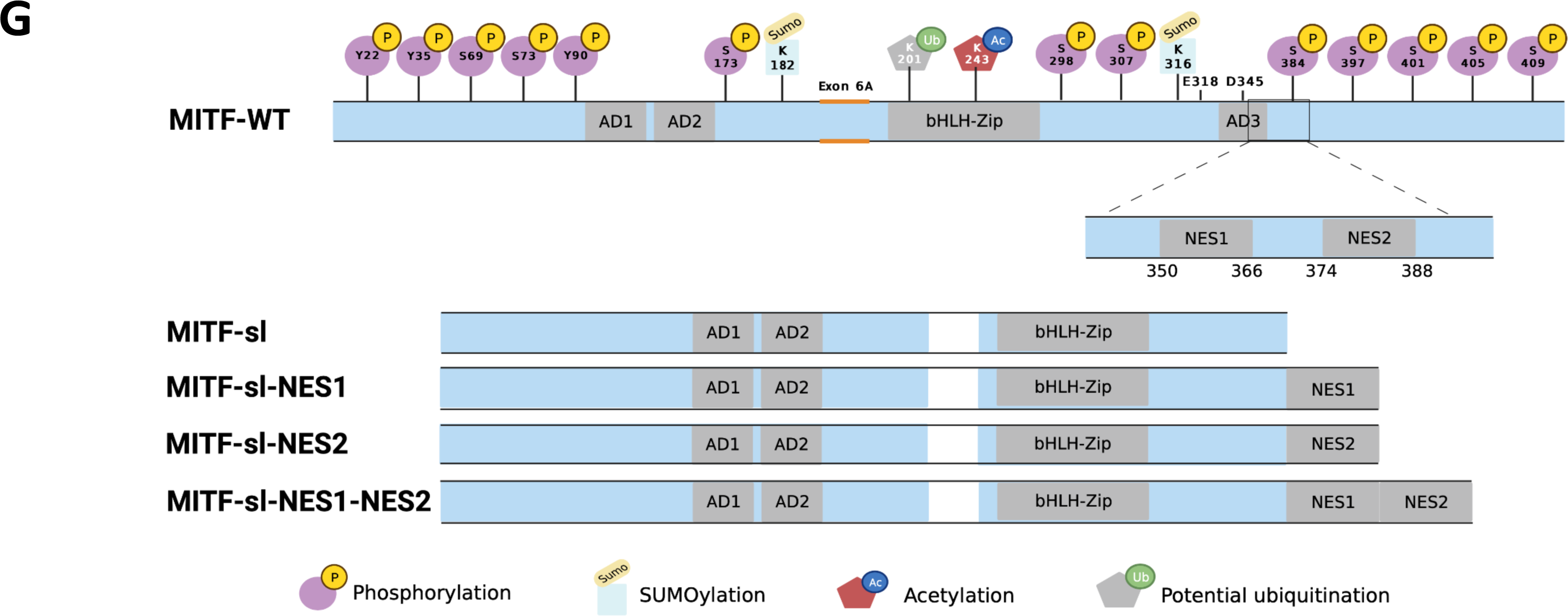
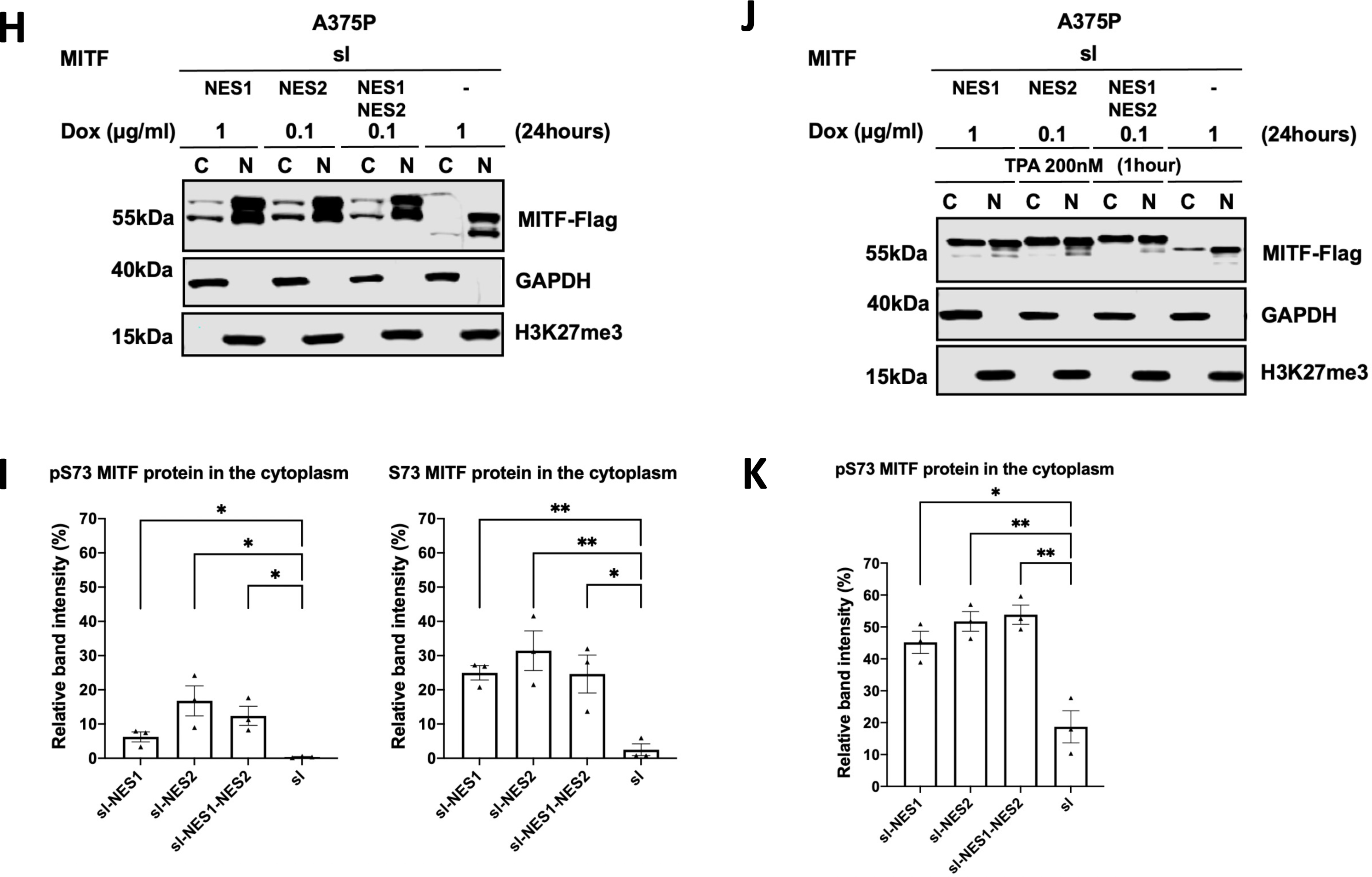
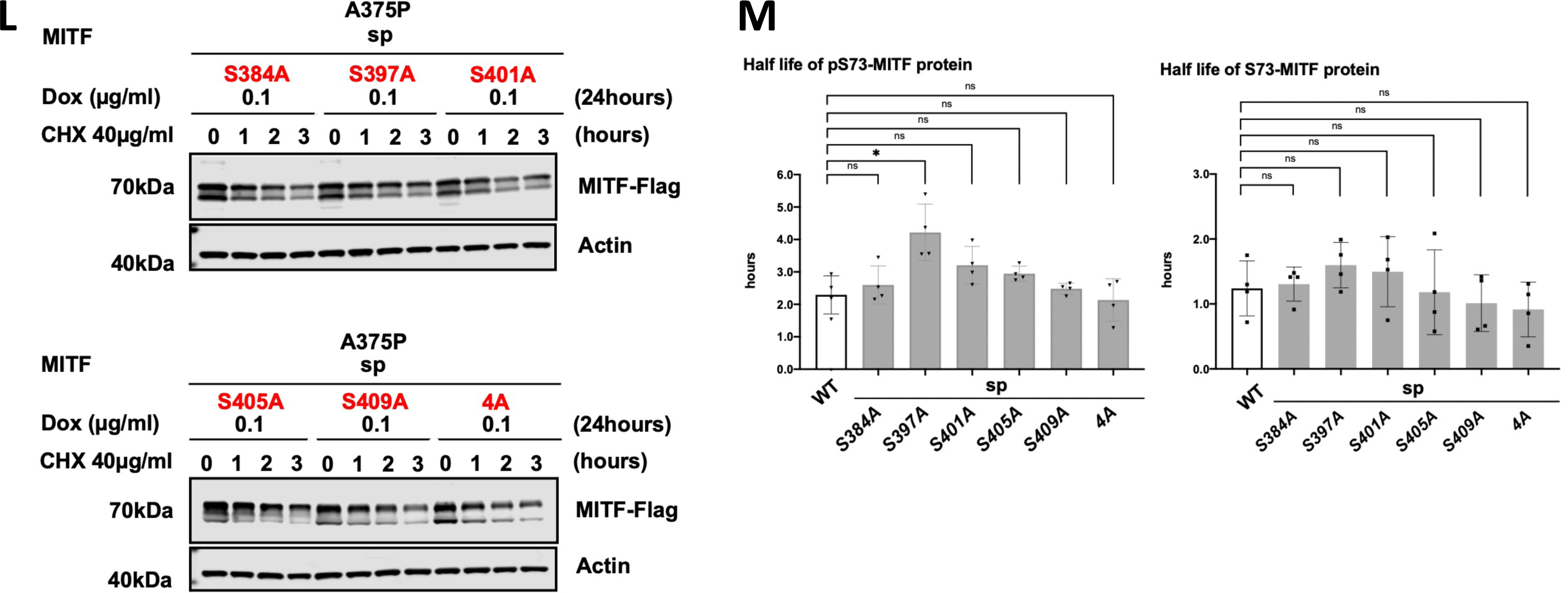
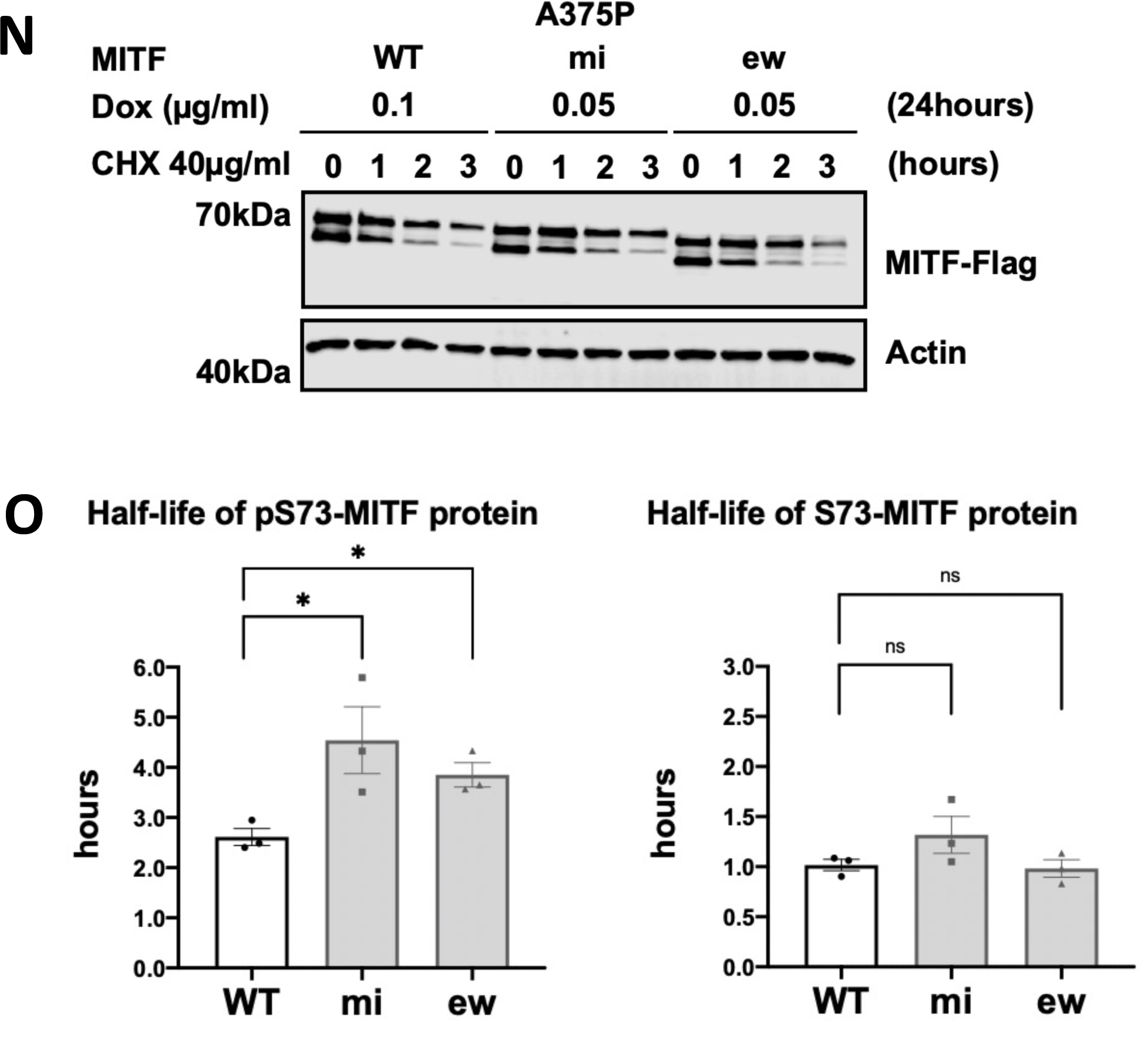
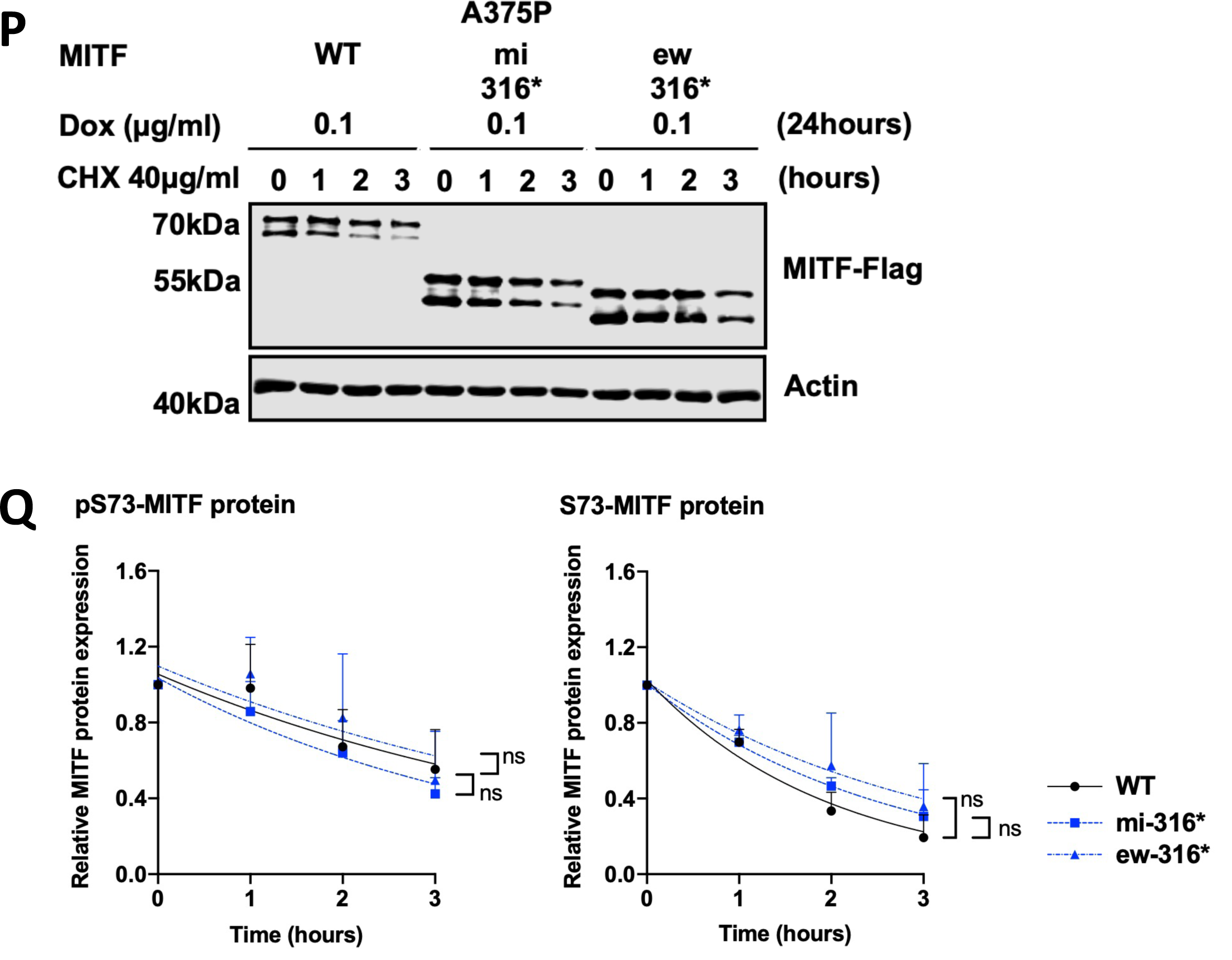
The four Individual phosphorylation sites at C-terminus (S384, S397, S401, and S405) did not affect MITF localization or stability. **(A), (C), (E),** and **(H)** Western blot analysis of cytoplasmic (C) and nuclear (N) fractions from A375P melanoma cells induced for 24 hours to overexpress the indicated MITF mutant proteins visualized using FLAG antibody. GAPDH and H3K27me3 were loading controls for cytoplasmic and nuclear fractions, respectively. **(J)** Western blot analysis of subcellular fractions isolated from A375P melanoma cells induced for 24 hours to overexpress the indicated MITF mutant proteins before treatment with 200nM TPA for 1 hour. MITF was visualized using a FLAG antibody. GAPDH and H3K27me3 were loading controls for cytoplasmic and nuclear fractions, respectively. **(B), (D)**, **(F)**, **(I),** and **(K)** MITF band intensities in the cytoplasmic and nuclear fractions from western blot analysis (A, C, E, H, and J, respectively) were quantified separately with *ImageJ* software and are depicted as percentages of the total amount of protein present in the two fractions. Error bars represent SEM of three independent experiments. Statistically significant differences (Student’s t-test) are indicated by * p< 0.05, *p< 0.05, ** p < 0.01, *** p < 0.001, **** p < 0.00, and ns not significant. **(G)** Graphical depiction of the MITF-WT, MITF-sl, MITF-sl-NES1, MITF-sl-NES2, and MITF-sl-NES1-NES2 proteins. The location of the NES1 and NES2 sequences in MITF-WT are also shown. **(L), (N) and (P)** The dox-inducible A375P cells were treated with doxycycline for 24h to express the indicated MITF proteins before treating them with 40 µg/ml CHX for 0, 1, 2, and 3 hours. The amount of MITF proteins were then visualized by western blot using FLAG antibody. Actin was used as a loading control. The band intensities were quantified using ImageJ software. **(M)** and **(O)** Half-life analysis of the pS73- and S73-MITF proteins over time after CHX treatment. The MITF protein levels relative to T0 were calculated, and non-linear regression analysis was performed. Error bars represent SEM of at least three independent experiments. Statistically significant differences (Student’s t-test) are indicated by * p< 0.05, *p< 0.05, ** p < 0.01, *** p < 0.001, **** p < 0.00, and ns not significant. **(Q)** Non-linear regression (one-phase decay) analysis of the indicated pS73- and S73-MITF proteins over time after CHX treatment in A375P melanoma cells. The relative MITF protein levels to T0 were calculated, and non-linear regression analysis was performed. Error bars represent SEM of at least three independent experiments. Statistically significant differences (Student’s t-test) are indicated by *p< 0.05, ** p < 0.01, *** p < 0.001, **** p < 0.00, and ns not significant.

**Figure S12:**
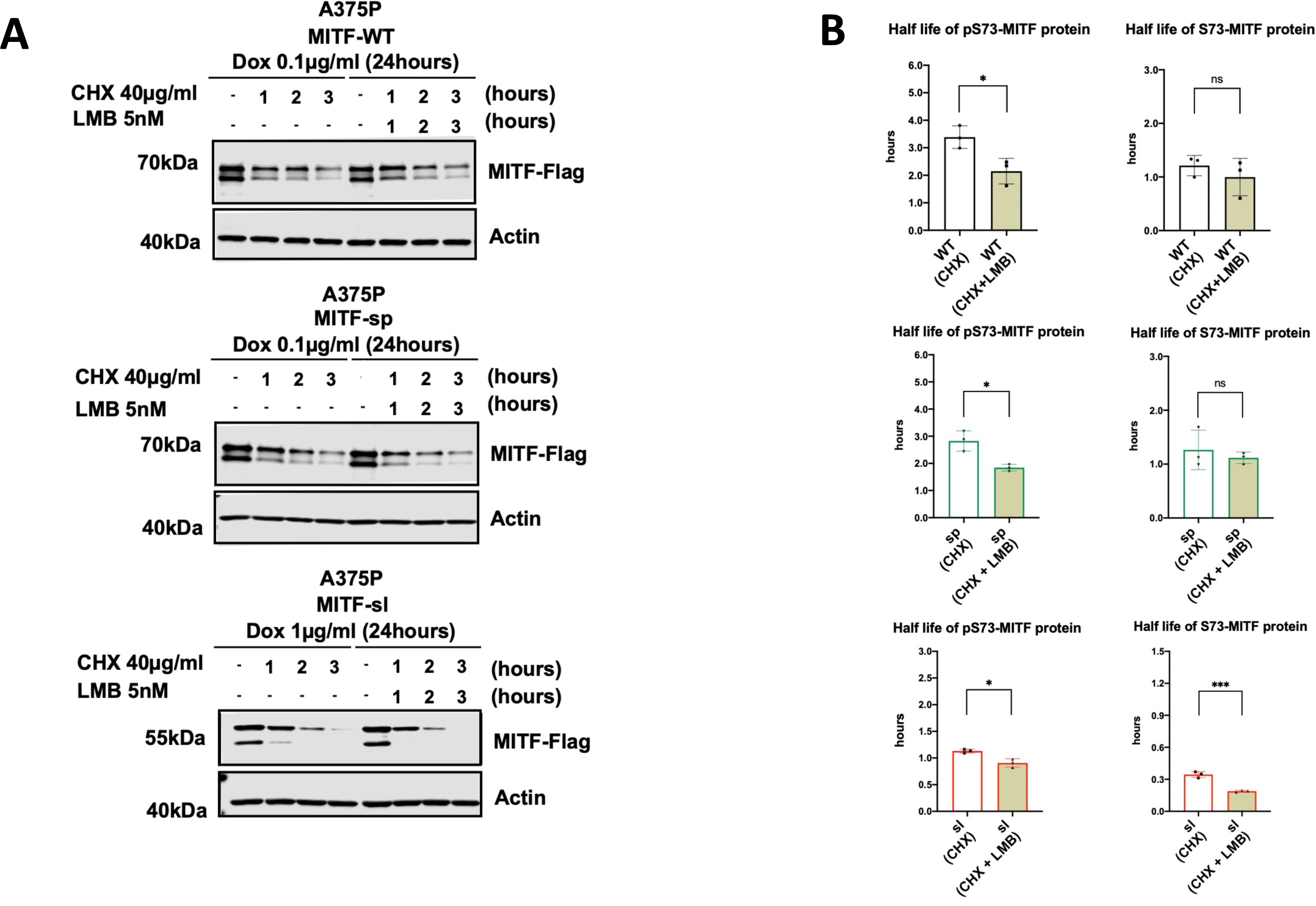
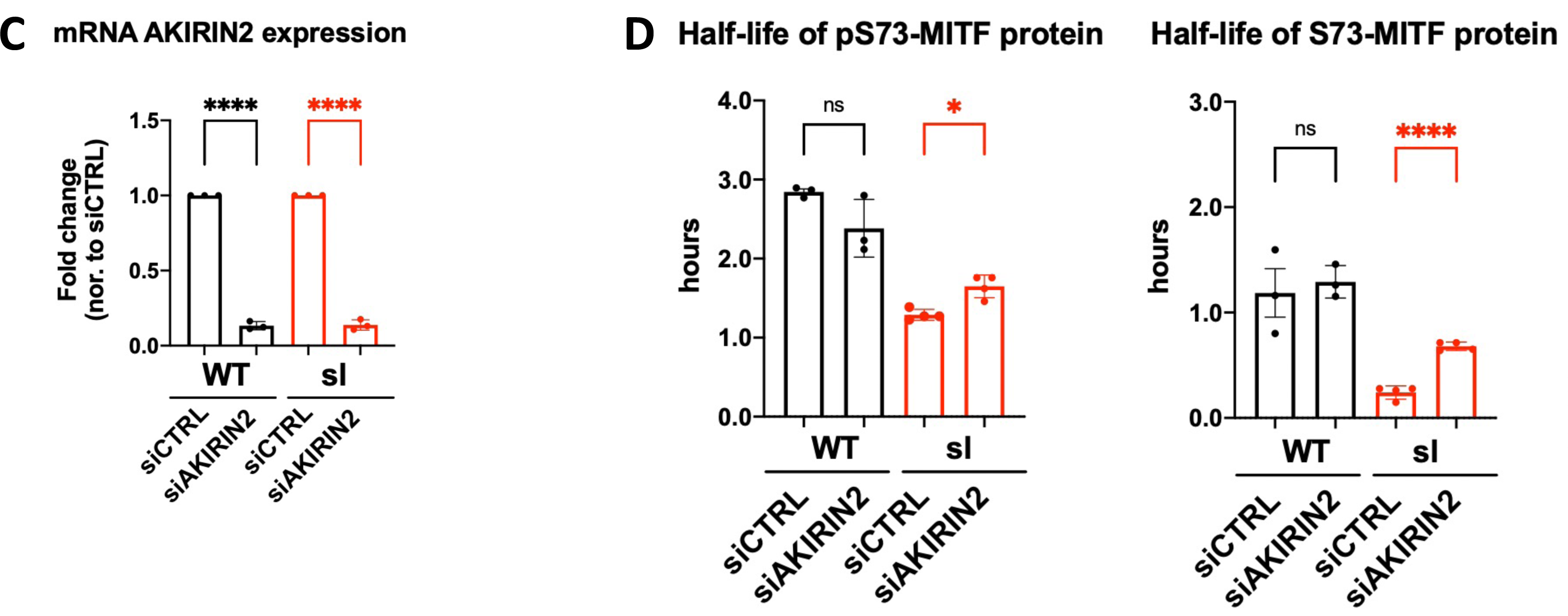
MITF is mainly degraded through the proteasome pathway in the nucleus. **(A)** Western blot analysis of the indicated MITF mutant proteins. The A375P cells were induced for 24 hours and treated with 40 µg/ml CHX in the presence of 5nM LMB for 0, 1, 2, and 3 hours. The MITF protein was visualized using FLAG antibody. Actin was used as a loading control. The band intensities were quantified using ImageJ software. **(B)** Half-life analysis of the pS73- and S73-MITF proteins over time after CHX plus LMB treatment. The MITF protein levels relative to T0 were calculated, and non-linear regression analysis was performed. Error bars represent SEM of at least three independent experiments. Statistically significant differences (Student’s t-test) are indicated by * p< 0.05, *p< 0.05, ** p < 0.01, *** p < 0.001, **** p < 0.00, and ns not significant. **(C)** RT-qPCR analysis of AKIRIN2 gene expression in A375P cells treated with siAKIRIN2 for 24 hours and then induced by dox to induce MITF overexpression for 6 hours. The expression was normalized to siCTRL-treated cells. Error bars represent SEM of at least three independent experiments. Statistically significant differences (Student’s t-test) are indicated by * p< 0.05, *p< 0.05, ** p < 0.01, *** p < 0.001, **** p < 0.00, and ns not significant. **(D)** Half-life analysis of the pS73- and S73-MITF proteins in siAKIRIN2-treated A375P cells which are induced to overexpress MITF. The MITF protein levels relative to T0 were calculated, and non-linear regression analysis was performed. Error bars represent SEM of at least three independent experiments. Statistically significant differences (Student’s t-test) are indicated by * p< 0.05, *p< 0.05, ** p < 0.01, *** p < 0.001, **** p < 0.00, and ns not significant.

**Figure S13:**
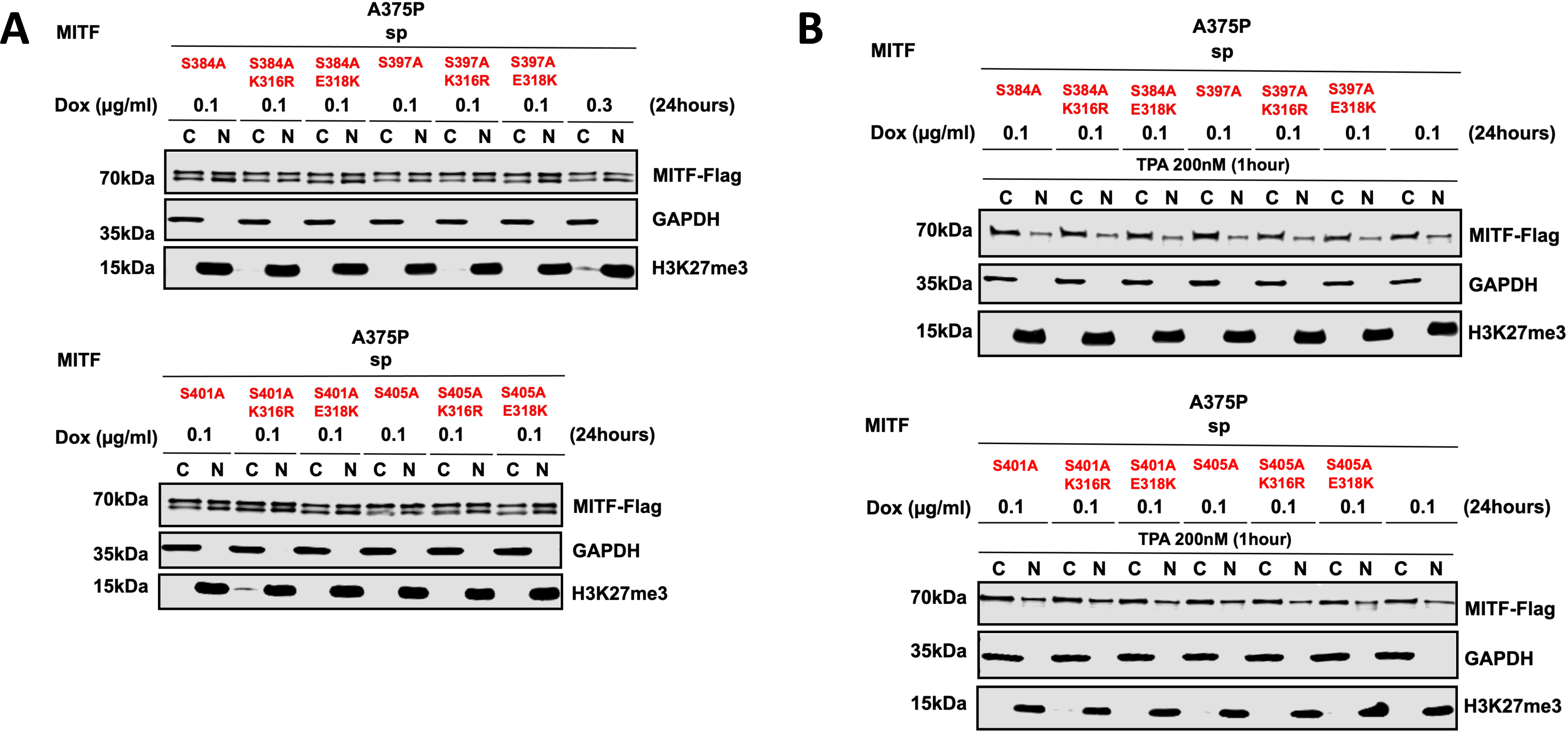
Mutation at four individual phosphorylation sites at C-end (S384, S397, S401, and S405) in combination with K316R and E318K do not affect MITF localization. **(A)** Western blot analysis of subcellular fractions isolated from A375P melanoma cells induced to overexpress the indicated MITF mutant proteins. MITF protein in cytoplasmic (C) and nuclear (N) fractions were visualized using FLAG antibody. GAPDH and H3K27me3 were loading controls for cytoplasmic and nuclear fractions, respectively. **(B)** Western blot analysis of subcellular fractions isolated from A375P melanoma cells induced to overexpress the indicated MITF mutant proteins before treatment with TPA at 200nM for 1 hour. MITF protein in cytoplasmic (C) and nuclear (N) fractions were visualized using FLAG antibody. GAPDH and H3K27me3 were loading controls for cytoplasmic and nuclear fractions, respectively.

## ACKNOWLEDGMENTS

We thank Joanne Dietz, Fran Dorsey, Latasha Crawford, Melanie Gasper, and Anjalie Parekh and the NINDS Animal Health and Care Section for maintaining mutant mouse lines, Linda S. Cleveland, Susan Skuntz, Christian Praetorius, Bryndís K Gísladóttir and Aðalheiður G. Hansdóttir for expert technical assistance, and the NINDS sequencing facility for support. We also thank Colin Goding for his critical comments on the manuscript. This work was supported by grant 217768 from the Icelandic Research Council (ES), the Icelandic Cancer Society (POH), and the University of Iceland Ph.D. Student Fund and by the intramural research program of the NIH, NCI, and NINDS.

## AUTHOR CONTRIBUTIONS

NHV, SS, KB, JHH, LL and ES designed and performed all molecular experiments. ES, DAS, NAJ and NCG conceptualized and performed the suppressor screen. HA, JD and KB studied developmental effects. MMV and POH performed smFRET analysis. ES supervised the study. The manuscript was written by NHV and ES with contributions from all authors.

